# Guided multi-agent AI invents highly accurate, uncertainty-aware transcriptomic aging clocks

**DOI:** 10.1101/2025.09.08.674588

**Authors:** Vinayak Agarwal, Orion Li, Christopher A. Petty, Timothy Kassis, Paul W.K.Rothemund, David A. Sinclair, Ashwin Gopinath

**Affiliations:** Blavatnik Institute, Dept. of Genetics, Paul F. Glenn Center for Biology of Aging Research at Harvard Medical School, Boston, MA 02115 USA; Department of Mechanical Engineering, Massachusetts Institute of Technology, Cambridge, MA 02139, USA; Department of Bioengineering, California Institute of Technology, Pasadena, CA 91125; Biostate.AI, 380 Portage Ave Palo Alto, CA 94306 USA; Bayoshiti AI, Koramangala, Bengaluru, Karnataka 560095, India

## Abstract

Scientific discovery has long relied on human creativity, with computation limited to analysis. Here we report an AI-guided system, K-Dense, that accelerates hypothesis testing and delivers robust scientific discoveries. Trained on ARCHS4 (57,584 samples, 28 tissues, 1,039 cohorts, ages 1–114), the unified ensemble clock achieved *R*^2^ = 0.854 and MAE = 4.26 years—while uniquely providing calibrated confidence intervals. This self-aware design flags predictions made at transitional or extreme ages, where biological heterogeneity peaks, suggesting clinical utility for uncertainty itself. Development revealed stage-specific markers, including CDKN2A/p16 (senescence), AMPD3 (muscle wasting), MIR29B2CHG (progeroid traits), and SEPTIN3 (neurodegeneration resistance). Sliding-window analysis across 85 overlapping ranges uncovered wave-like shifts in gene importance, showing that transcriptomic aging signatures evolve continuously, not discretely. By transforming biological age assessment from static point estimates to calibrated predictions with explicit uncertainty, this approach establishes reliable and interpretable clocks. Beyond the clock itself, K-Dense demonstrates how guided AI can compress months of exploration into weeks, pointing toward a scalable framework for accelerated scientific discovery.

## Introduction

The exponential growth of scientific data has not been matched by a proportional rise in tranformative discoveries. Each year, 2.5-3.0 million academic publications are produced, yet the pace of breakthrough innovation has plateaued [1]. Although vast amouns of data, from real-time sensors to large-scale laboratory assays, are continuously collected, their systematic use for optimization or causal inference remains uncommon [2]. Advances in artificial intelligence and big data analytics have demonstrated transformative potential, exemplified by AlphaFold in protein structure prediction [3]. However, despite dramatic improvements in robotic systems and high-throughput data generation, the human capacity to generate hypotheses, integrate results and synthesize new insights has become the principal bottleneck in scientific progress.

Recent computational frameworks, including Google’s Co-Scientist [4], Sakana’s AI Scientist [5] and ChemCrow [6], highlight the potential of AI to help with individual research tasks such as locating literature, generating hypothesis, or performing targeted data analysis. However, these systems address only a fraction of the broader research cycle. We argue that accelerating discovery requires automating this cycle in a more integrated manner, spanning literature survey and hypothesis formulation to data analysis and synthesis of results. In this work, we demonstrate that such a cycle can be automated for experimental data interpretation under human supervision, focusing on the interpretive core of research rather than data generation, which is already advancing rapidly through robotics and high-throughput platforms.

We developed K-Dense (Knowledge Dense), an integrated multi-agent research system built on Google’s Gemini 2.5 Pro foundation model that amplifies human research capabilities through rapid implementation and testing of researcher-directed hypotheses. Unlike tools limited to isolated analyses or single tasks, K-Dense coordinates multiple specialized agents that collectively implement the broader research cycle. When guided by researchers, the system can aggregate and analyze literature to identify knowledge gaps, generate and refine hypotheses, apply and compare analytical approaches, perform multilevel data interpretation, contextualize results, and assist in documentation and manuscript preparation. These capabilities are distributed among four coordinated agents, Literature Reviewer, Research Planner, Coder & Analyst, and Writer & Summarizer, that operate through iterative refinement of hypotheses and methodologies. Critically, K-Dense incorporates reflexion loops that detect and correct errors, reduce hallucinations, and prevent AI manipulation, with the aim of ensuring that the output remains scientifically reliable. To the best of our knowledge, these measures minimize unintended errors, but we acknowledge that some leakage may persist despite extensive human supervision. Therefore, additional safeguards and future work are needed to further strengthen robustness and reproducibility (Figure 1a; see **Supplementary Note 1** for technical architecture).

**Figure 1.**
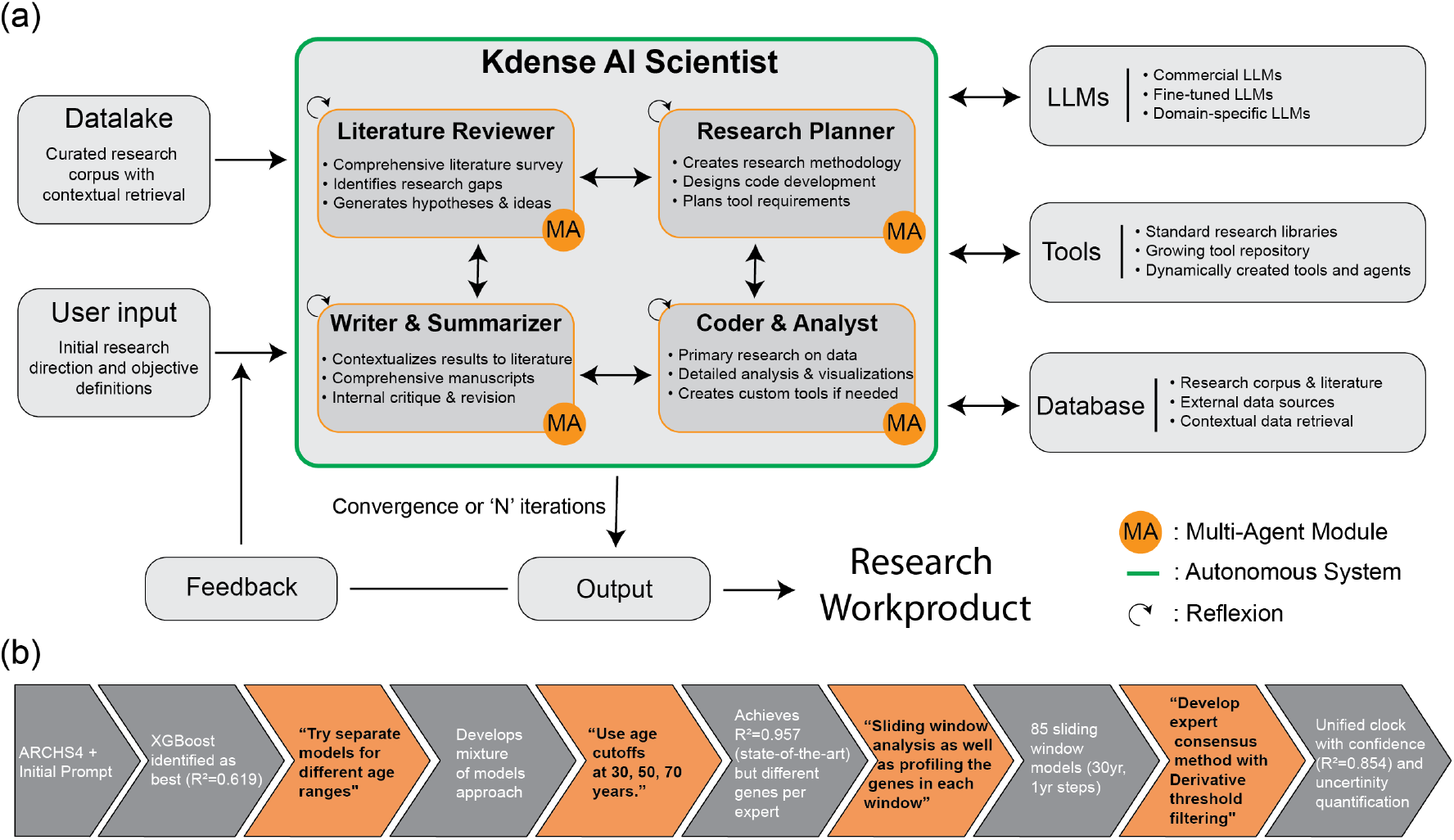
K-Dense Multi-Agent Research System. Schematic representation of the multi-agent system for accelerated scientific research under principal investigator supervision. (A) General framework showing coordination of four specialized computational agents through PI-directed iterative cycles: Literature Reviewer (literature aggregation and knowledge synthesis), Writer & Summarizer (documentation and manuscript assistance), Research Planner (hypothesis generation and experimental design support), and Coder & Analyst (sophisticated data analysis and implementation). External components include tool integration, database access, and system control for resource management. The PI determines research direction, validates hypotheses, and interprets results at each iteration.[7] (B) Aging research implementation workflow demonstrating the iterative development process. Orange arrows indicate decision points where researcher expertise guided methodological choices based on K-Dense’s analytical outputs. Gray arrows show autonomous system progression. The workflow progressed from initial ARCHS4 exploration through hypothesis testing (“Try separate models for different age ranges”) to methodological refinement (sliding window analysis) and final implementation (unified clock with uncertainty quantification). See **Supplementary Note 1 & 2** for detailed prompts and system outputs at each intervention point.

To demonstrate the value of this supervised multi-agent framework, we applied it to biological aging, a domain that requires complex data integration and recognition of subtle molecular patterns. Biological age clocks quantify the deviation between molecular aging profiles and chronological age, serving as critical tools in geroscience to evaluate the efficacy of interventions and to support personalized medicine applications [8]. The field began with DNA methylation-based predictors such as Horvath’s pan-tissue clock [9] and Hannum’s blood-specific clock [10], which achieved remarkable accuracy (*r >* 0.9). Subsequent generations, including PhenoAge[11] and GrimAge[12], extended these models to predict morbidity and mortality rather than chronological age [13]. However, methylation clocks function primarily as statistical correlates, with CpG sites often lacking clear functional relationships to established aging pathways. This black-box nature limits their utility for understanding mechanisms or identifying targetable interventions[14]. While proteomic clocks have emerged as a more mechanistically grounded alternative, since circulating proteins reflect functional states of tissues and pathways involved in aging [15]. These clocks provide biological insight beyond correlation, but their development is constrained by limited datasets[16], incomplete proteome coverage in current assays[17], and relatively high measurement costs[18]. For these reasons, we chose not to focus on proteomic clocks in this study and instead focus on transcrptomic clocks that provide a more accessible and mechanistically interpretable option, as gene expression directly reflects cellular activity and can be mapped onto specific biological pathways [19].

Unlike static methylation marks, transcriptome capture dynamic responses to environmental stimuli, disease states, and therapeutic interventions. Recent transcriptomic clocks have achieved respectable accuracy, with RNAAgeCalc reporting a mean absolute error (MAE) of 7.8 years [20] and more recent approaches reaching 4.82 years [21]. However, there is no consensus on optimal gene sets, with different studies identifying largely non-overlapping aging signatures. Current approaches also tend to treat aging as a uniform process across the lifespan despite evidence that distinct biological processes govern growth, maintenance, and decline [14]. This leads to missed signals such as the fundamental shift from anabolic to catabolic dominance that characterizes aging trajectories. In addition, most transcriptomic clocks are trained on homogeneous cohorts and show degraded performance when applied to diverse populations or technical platforms, which limits their clinical utility.

We directed K-Dense to analyze the ARCHS4 database [22], which uniformly reprocesses RNA-seq data from thousands of Gene Expression Omnibus studies. This choice was intentional: we sought to demonstrate that K-Dense can reuse and integrate existing data at scale, rapidly constructing cohorts and performing analyses that would otherwise require extensive manual curation. At this stage, focusing on interpretation highlights the system’s ability to make data processing seamless, unlocking insights that remain hidden in fragmented datasets. In principle, this interpretive core can also be coupled downstream with automated data generation platforms, creating a fully closed research loop, but that lies beyond the scope of the present study. The ARCHS4 collection, with 57,584 samples spanning 1,039 independent studies across diverse tissues, protocols, and populations, provides a rigorous test case. Although harmonization mitigates some computational variation, the main sources of batch effects, including sample collection methods, RNA extraction, library preparation, and sequencing platform differences, remain [23]. Achieving strong performance under these conditions requires the capture of robust biological signals rather than study-specific artifacts, making ARCHS4 an ideal benchmark to evaluate both transcriptomic clocks and K-Dense capabilities.

We initiated K-Dense with an exploratory objective: *‘Explore the relationship between aging and transcriptomics*.*’* Through iterative human-AI collaboration, we directed the system to test specific hypotheses about age-stratified modeling, with K-Dense executing comprehensive analyses and identifying patterns that informed our next research decisions (Figure 1b, orange arrows indicate human intervention points; see **Supplementary Note 1** for detailed PI-system interaction protocol), ultimately developing three key methodological innovations. First, a Multi-Model Architecture of age-stratified XGBoost models that capture distinct molecular processes across life stages. Second, a sliding-window ensemble spanning 85 overlapping age windows, while sliding-window approaches have been applied in genomic variation detection [24] and time-series biomarker discovery [25, 26], this represents their first application to transcriptomic age modeling, revealing continuous gene importance transitions across the lifespan. Third, a consensus prediction framework that adapts ensemble methods successful in weather forecasting [27] to biological age assessment. Unlike traditional ensemble approaches that simply average predictions, our framework uses derivative-threshold filtering to identify stable expert consensus, transforming model disagreement into a biological signal where low confidence indicates heterogeneous aging patterns or phase transitions.

This supervised exploration yielded transformative insights beyond the methodological advances. The system identified novel aging biomarkers including CHAMP1 (chromosomal alignment), MIR29B2CHG (ECM remodeling and aging regulation), and SEPTIN3 (synaptic vesicle trafficking, AD-associated) alongside validation of established markers like CDKN2A/p16. Most strikingly, the analysis quantified a fundamental reversal in pathway activation across the lifespan, with proliferation pathways declining from youth to senescence while stress-response pathways show the inverse trajectory. In the following section, we detail how this computational exploration of 57,584 samples across 1,039 studies led to a transcriptomic clock that not only achieves exceptional accuracy but also provides mechanistic insights into the aging process.

### Development of Transcriptomic Aging Clock

The development of a transcriptomic clock from our openended directive unfolded through successive hypothesis tests and refinements. Initially, K-Dense approached the ARCHS4 database much as earlier studies had, by seeking a single, universal model to predict age across all samples. Benchmark comparisons across algorithms (XGBoost, LightGBM, LinearSVR, Ridge, ElasticNet) confirmed that XGBoost consistently outperformed other methods, though even the best single global model achieved only moderate accuracy. XG-Boost’s architecture may itself be biologically relevant: its cascading decision trees parallel how aging unfolds through sequential molecular events, where early damage creates errors that subsequent processes must compensate for, and tree-based thresholds naturally capture phenomena such as senescence activation when damage exceeds repair capacity. Yet despite this advantage, the single-model approach showed systematic degradation at the extremes of age, particularly among very young and very old samples.

This heterogeneity raised a fundamental question: what if aging is not a single continuous process, but a succession of distinct biological processes? Guided by this hypothesis and upon explicit instruction from the investigators (see Supplementary Note 1 for detailed prompts and system outputs), K-Dense tested age-windowed models. We further directed the system to optimize toward four distinct windows, yielding the Multi-Model Architecture (MMA). The final design employed four XGBoost models covering deliberately simple age ranges (Young: 1-30, Early Middle: 30-50, Late Middle: 50-70, Elderly: 70+ years). These cutoffs were not optimized, but chosen to demonstrate the feasibility of age-stratified modeling. Each model captured stage-specific biology with moderate performance (*R*^2^ ≈ 0.68 − 0.74), while their ensemble combination achieved *R*^2^ = 0.957 and mean absolute error of 3.7 years, slight improvement over the state-of-the-art transcriptomic clock (MAE 4.82 years) reported by Qi and colleagues [21].

When compared across multiple machine learning methods (Figure 2a-b), the MMA demonstrated superior coefficient of determination (*R*^2^ = 0.957) versus single global XGBoost (*R*^2^ = 0.619), LightGBM (*R*^2^ = 0.604), LinearSVR (*R*^2^ = 0.574), Ridge (*R*^2^ = 0.539), and ElasticNet (*R*^2^ = 0.310). Age-stratified prediction analysis across 11,517 test samples (Figure 2c) revealed tight clustering around the perfect prediction diagonal, with age group-specific performance (Figure 2d) maintaining MAE below 5 years from ages 20-80. Performance remained stable across most of the lifespan, with mean absolute error below 5 years from ages 20-80. Predictability degraded in nonagenarians (MAE up to 12.1 years), reflecting both sparse data and the biological heterogeneity of extreme aging. Notably, the ARCHS4 dataset itself exhibited a bimodal age distribution (peaks at 21 and 71 years; Supplementary Fig. S2c, which likely accentuated these edge effects.

**Figure 2.**
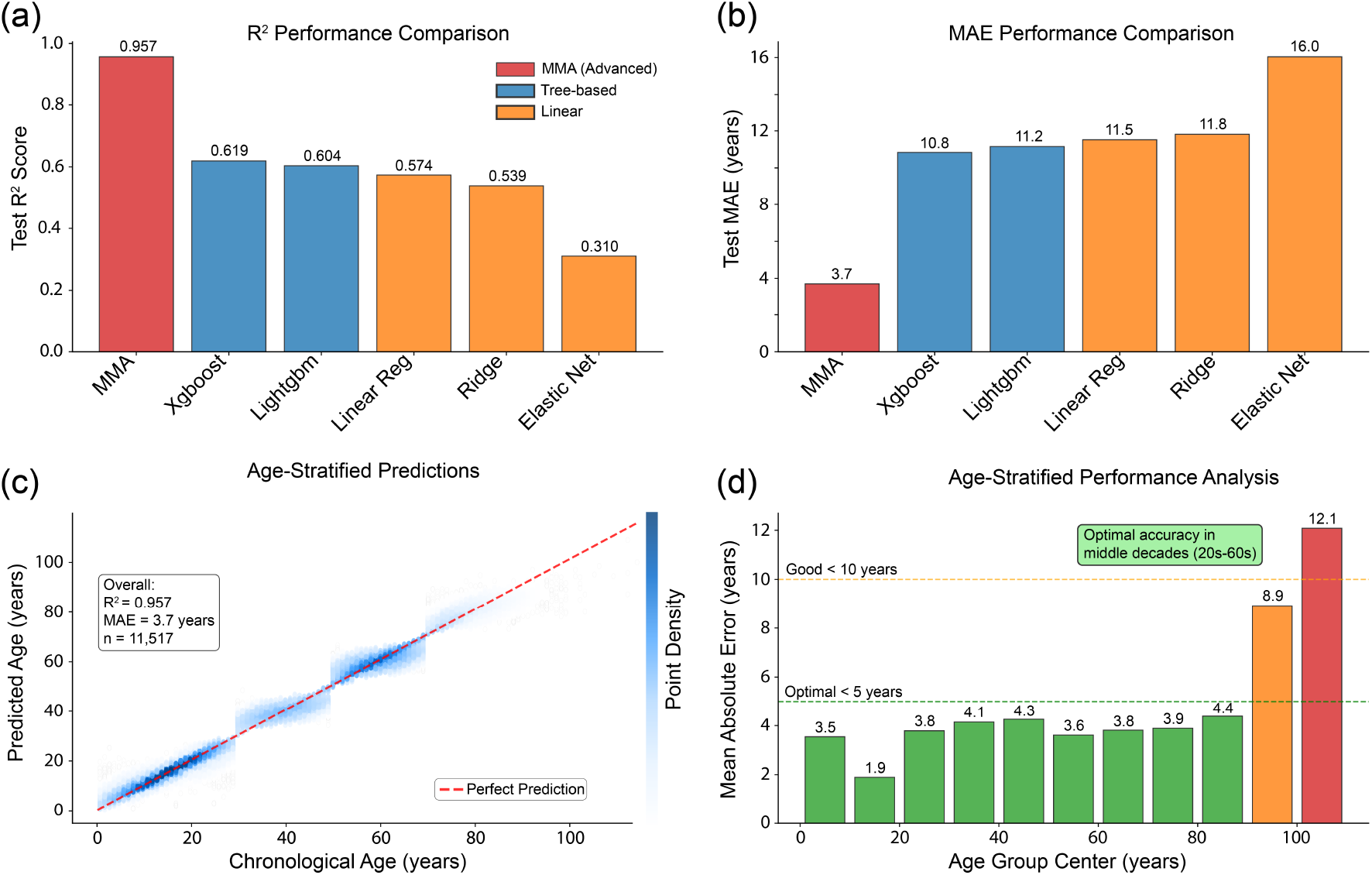
Model Performance and Comparative Analysis. K-Dense ‘s development of state-of-the-art transcriptomic aging clock. (A) Coefficient of determination (R_2_) comparison across machine learning approaches showing MMA architecture superiority (R_2_ = 0.957) over XGBoost (0.619), LightGBM (0.604), LinearSVR (0.574), Ridge (0.539), and ElasticNet (0.310). (B) Mean Absolute Error analysis confirming MMA model’s lowest prediction error (3.7 years) compared to traditional approaches (10.8-16.0 years). (C) Age-stratified prediction accuracy across 11,517 samples showing excellent correlation between predicted and chronological ages across four age cohorts with tight clustering around perfect prediction diagonal. (D) Age group-specific performance revealing optimal accuracy in middle decades (MAE < 5 years) with degradation at extremes reaching 12.1 years in nonagenarians.

The methodological innovations developed through this iterative process included three key components: age-stratified specialization dividing samples into biologically meaningful ranges; density-weighted regression to correct sampling imbalances; and heteroscedastic uncertainty quantification to capture increasing transcriptomic variability with age. Technical validation confirmed robust data quality across 5,000 genes and consistent sequencing coverage (Supplementary Fig. S3).

Despite strong predictive performance, the MMA has a critical limitation: it requires chronological age input for model selection, preventing its use as a true standalone biological clock. We repeatedly attempted to train a gating function that could infer the correct window directly from transcriptomic profiles, but these efforts failed to achieve reliable accuracy. As a result, predictions near age-window boundaries were especially unstable, with errors depending on which expert was selected. This dependency on external age input restricts the MMA’s direct clinical utility, even as it provides valuable insight into stage-specific aging processes. We present it here not as our final solution but as an intermediate finding that revealed important biological insights. These stage-specific patterns directly motivated our development of the age-agnostic ensemble approach, demonstrating how methodological limitations can illuminate biological understanding.

### Biological Interpretation Framework

Having established that the Multi-Model Architecture achieves state-of-the-art accuracy, we next examined which genes were driving predictions within each life stage. Previous clocks have typically identified markers through regression coefficients or penalized feature selection [11, 12, 21]. In contrast, we applied model-based feature importance using the XGBoost gain metric [28], an approach with precedent in biomarker discovery, where tree-based methods have been used to classify cancer subtypes [29, 30], predict drug response [31, 32], and uncover disease-associated genes [33, 34, 35, 36]. This method quantifies each gene’s contribution to reducing prediction error in the context of all other genes, inherently capturing gene–gene interactions and non-linear relationships. While highly informative, we emphasize that feature importance reflects correlation rather than causality: a high score indicates association with aging, not necessarily mechanistic involvement.

Analysis of the four age windows revealed distinct, stagespecific signatures (Figure 3). In the **young** cohort (1–30 years), MIR29B2CHG was the most important predictor, consistent with its role in tissue maintenance and progeroid phenotypes when deficient. CDKN2A/p16 was already detectable in this window, suggesting senescence processes begin earlier than often assumed. In **early middle age** (30–50 years), RBP1 emerged as most predictive, with AMPD3 rising in prominence, a muscle metabolism gene whose activity reflects pre-sarcopenic changes. In the **late middle** window (50–70 years), CHAMP1 became dominant, highlighting genomic stability pathways, while AMPD3 maintained high importance. Finally, in the **elderly** (70+ years), SEPTIN3 was the strongest predictor, a neuronal protein associated with synaptic vesicle trafficking and Alzheimer’s disease susceptibility [37].

**Figure 3.**
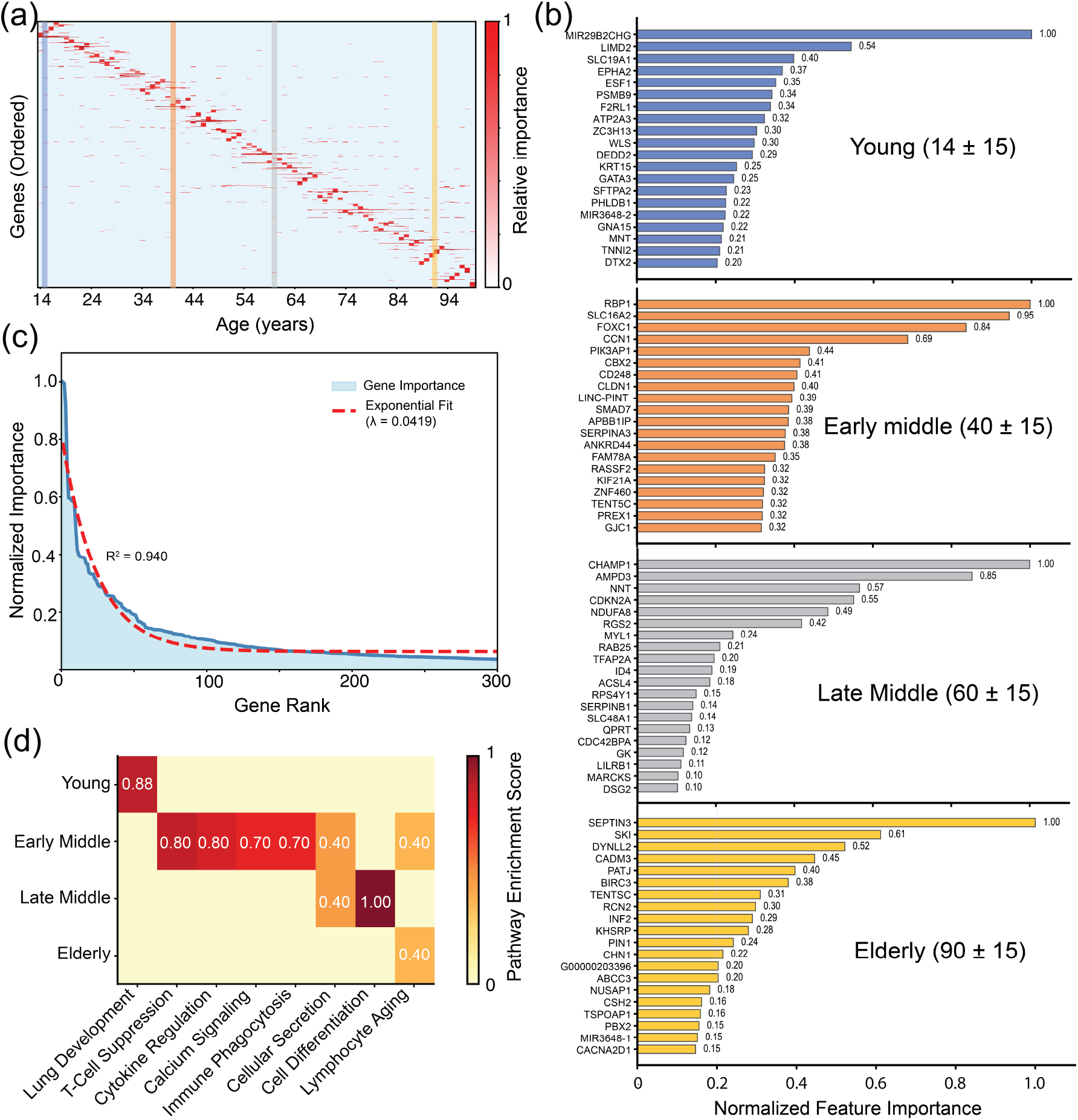
Stage-Specific Gene Signatures and Pathway Dynamics Across the Lifespan. (a) Sliding-window gene importance heatmap across ages 14-94 years displaying temporal dynamics of 5,000 genes ordered by weighted peak importance age. Column-normalized visualization (red=high, white=low importance) reveals wave-like horizontal bands with distinct temporal patterns. Vertical lines mark age boundaries used in the Multi-Model Architecture. (b) Top 20 age-predictive genes for four life stages showing stage-specific molecular signatures. Young (14±15): MIR29B2CHG (1.00), LIMD2 (0.54), SLC18A1 (0.40); Early middle (40±15): RBP1 (1.00), SLC10A2 (0.95), FOXC1 (0.84); Late middle (60±15): CHAMP1 (1.00), AMPD3 (0.85), NNT (0.57); Elderly (90±15): SEPTIN3 (1.00), SKI (0.61), DYNLL2 (0.52). Note CDKN2A/p16 begins to appear in importance rankings, though not yet among the top predictors (Supplementary Data). (c) Gene importance distribution across top 300 genes following exponential decay (*λ*=0.0419, *R*^2^=0.940), demonstrating concentrated predictive power in limited gene sets rather than diffuse transcriptome-wide signals. (d) Pathway enrichment scores across four age groups revealing quantitative reversal of biological processes. Proliferation pathways (Lung Development) show maximal enrichment in youth (0.88) declining to elderly (0.40), while stress-response pathways (Lymphocyte Aging) demonstrate inverse pattern (0.40 in youth to 1.00 in elderly). Numbers indicate composite enrichment scores integrating gene counts, importance values, expression changes, and statistical significance. This quantifies the theoretical anabolic-to-catabolic shift in human aging. Note: White regions indicate zero or negligible importance values. Gene ordering data available in Supplementary Data `file gene_ordering_AllTissues_data_sliding_window.csv`.

Markers such as CDKN2A and AMPD3 appeared across multiple windows with varying importance, suggesting persistent but context-dependent contributions to aging trajectories. At the same time, the near-complete turnover of top-ranking genes between stages underscores that no single universal biomarker set exists, validating the multi-model approach.

Beyond individual genes, the distribution of importance scores followed an exponential decay (Figure 3c; *λ* = 0.0419, *R*^2^ = 0.940), indicating that aging signals concentrate within a limited set of key pathways rather than being broadly diffuse. Pathway enrichment analysis (Figure 3d) revealed a striking reversal: proliferation-related pathways were maximally enriched in youth (0.88) and declined steadily with age, while stress-response and senescence pathways increased, reaching maximal enrichment in the elderly. This quantitative flip provides direct empirical support for the long-hypothesized anabolic-to-catabolic transition in human aging [38].

Together, these findings demonstrate that accurate age prediction relies on distinct sets of genes across life stages, with little overlap between cohorts, and that aging reflects shifting molecular priorities rather than a static process. Detailed analyses of the top 20 genes in each age window, including functional annotations and stage-specific roles, are provided in Supplementary Data. The sharp boundaries of our four-window model, however, raised a critical question: do these molecular signatures change gradually rather than discretely across life stages? This observation motivated our next analysis, where we moved beyond fixed windows to explore aging biomarkers in a continuous sliding-window framework spanning the entire lifespan.

### Mapping Age-Progressive Gene Dynamics

While the four fixed windows of the Multi-Model Architecture revealed distinct sets of age-predictive genes, their sharp boundaries risked missing gradual molecular transitions across the lifespan. To overcome this limitation, we developed a continuous analysis using 85 overlapping 30-year sliding windows advancing in 1-year increments (ages 1-30, 2-31, …, 85-114). For each window, we trained an XGBoost model and computed feature importance scores using the gain metric, retaining the top 5,000 genes. Genes were then ordered by their weighted peak importance age, calculated as the weighted average of windows where each gene exceeded 30% of its maximum importance, with similar-peaking genes clustered to preserve pattern continuity. This approach created a temporally resolved heatmap of age-gene importance (Figure 3a).

This analysis revealed striking “wave-like” patterns: horizontal bands of genes rising in importance at specific life stages and declining thereafter. Genes were not uniformly predictive across the lifespan; rather, most contributed strongly for a limited interval, typically 20-30 years, before fading, while others emerged later. For example, youth-associated genes dominated in early decades, giving way to metabolic and immune regulators in middle age, and finally to stress-response and synaptic maintenance genes in late life. The continuous progression of these bands demonstrates that aging signatures evolve smoothly rather than switching abruptly between discrete cohorts.

The wave structure further emphasizes that no single universal biomarker set exists; instead, accurate prediction depends on shifting molecular drivers across time. This observation also suggested a natural strategy for integrating predictions across models without requiring chronological age input, a concept that underlies the ensemble approach described later in this paper (Unified Transcriptomic Clock section).

Having established that transcriptomic aging signatures evolve continuously across life, we next asked whether this temporal structure is universal across tissues. The crosstissue analysis addressed this question directly and revealed a surprising finding: while all tissues exhibit the same temporal wave pattern, each employs distinct gene sets to implement it.

### Tissue-Specific Aging process

The wave-like age signatures identified in the wholetranscriptome analysis raised an important question: is this temporal progression universal across tissues, or does each organ follow its own trajectory? To address this, we next asked whether this temporal pattern holds within individual tissues. We first screened 28 tissues for sample availability and age coverage (Supplementary Data). Only **blood** (*n* ≈ 4,285) and **colon** (*n* ≈ 1,254) provided sufficient coverage across the lifespan to support stable sliding-window models; therefore, detailed tissue-specific maps are shown for these two organs.

Across the two tissues, we observed the same temporal wave structure: distinct bands of genes rising and falling in predictive importance across the lifespan (Figure 4b). This striking conservation demonstrates that aging proceeds through shared temporal phases across the human body. However, the genes responsible for these patterns were largely non-overlapping between tissues. For example, blood and colon both exhibited nearly identical temporal trajectories, yet their predictive gene sets showed minimal intersection (Figure 4b, bottom panel).

**Figure 4.**
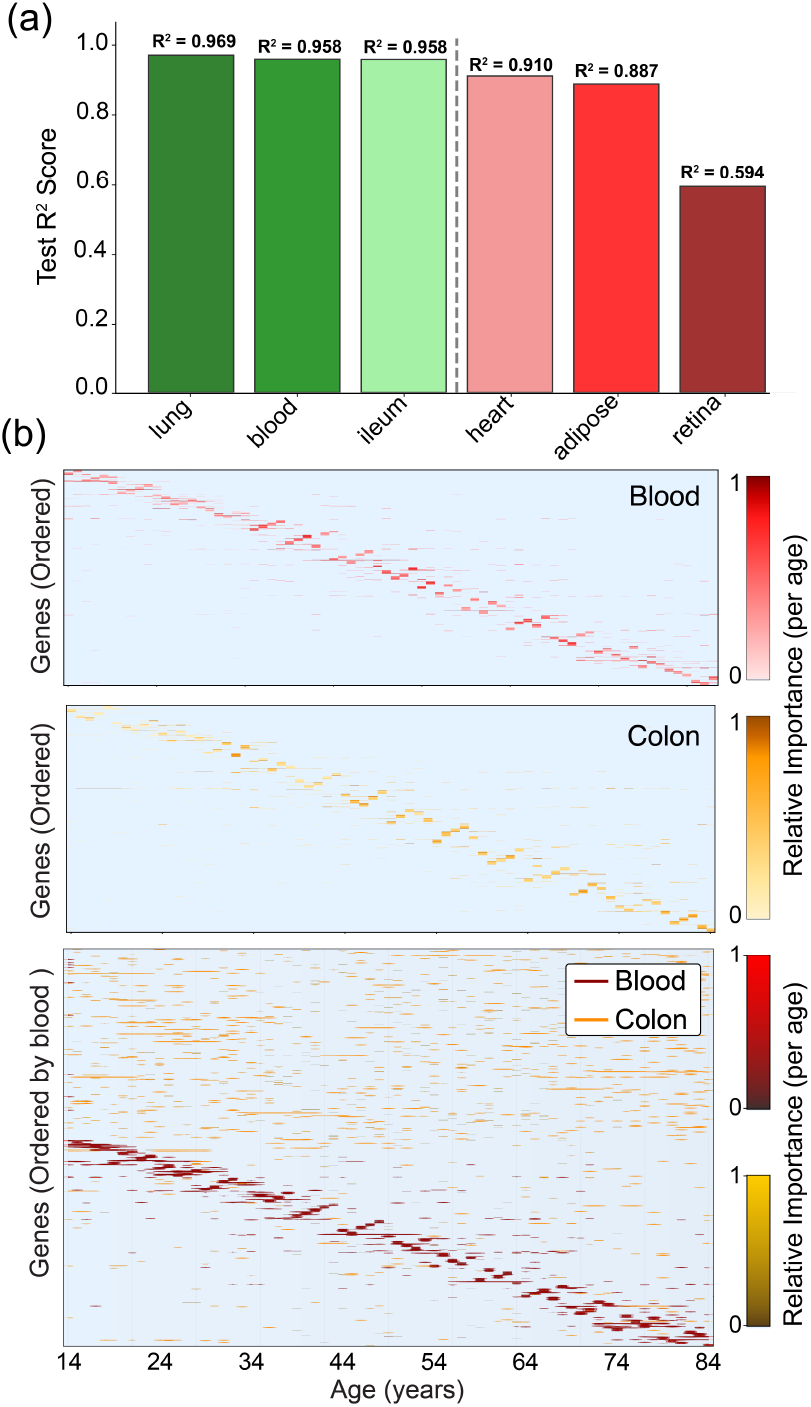
Tissue-Specific Performance and Aging Patterns. (a) Organ-specific performance analysis displaying three best (lung: *R*^2^ = 0.969, *n* = 1, 141; blood: *R*^2^ = 0.958, *n* = 4, 264; ileum: *R*^2^ = 0.958, *n* = 148) and three worst performing tissues (heart: *R*^2^ = 0.910, *n* = 204; adipose: *R*^2^ = 0.887, *n* = 109; retina: *R*^2^ = 0.594, *n* = 100) from comprehensive 28-organ analysis. (b) Sliding-window gene importance patterns for blood and colon tissues across ages 14-94 years. Top panels show individual tissue patterns with characteristic wave structure. Bottom panel overlays blood (blue) and colon (red) importance patterns, revealing non-overlapping gene sets despite identical temporal trajectories. Tissue-specific gene ordering data available in Supplementary Data files.

Performance analysis revealed both strong and weaker tissues (Figure 4a). Lung (*R*^2^ = 0.969, *n* = 1,141), blood (*R*^2^ = 0.958, *n* = 4,264), and ileum (*R*^2^ = 0.958, *n* = 148) achieved the highest accuracies, despite large differences in sample size. Heart (*R*^2^ = 0.910, *n* = 204), adipose (*R*^2^ = 0.887, *n* = 109), and retina (*R*^2^ = 0.594, *n* = 100) ranked among the lowest, with retina as an outlier likely due to limited samples and its unique developmental origin. Importantly, even the weakest-performing tissues (excluding retina) showed *R*^2^ *>* 0.88, confirming robust agerelated transcriptional changes across the body. These results, however, should be interpreted with caution: performance is influenced by the number of available samples, and some tissues (e.g., liver, pancreas) may also show degraded signal quality due to increased RNA damage or variable preanalytical handling.

These findings suggest that aging operates as a coordinated biological process: all tissues progress through common temporal phases, yet each implements them with distinct molecular components. This duality, universal timing with tissuespecific execution, provides a plausible explanation for why transcriptomic clocks trained on one tissue often fail to generalize to others. It also implies that while systemic interventions might align with shared temporal phases, tissue-targeted strategies may be required to modulate the molecular mechanisms underlying those phases.

### Unified Transcriptomic Clock with Uncertainty Quantification

The stage-specific models of the Multi-Model Architecture demonstrated that aging is best captured by distinct predictors across life stages, but their use as a “clock” was limited: chronological age had to be provided to select the appropriate model, and predictions were unstable near window boundaries. The sliding-window analysis confirmed that aging signatures evolve continuously rather than discretely, motivating an ensemble strategy to integrate predictions across the entire lifespan without requiring prior age input.

To address this, we trained 85 overlapping models spanning 30-year windows (ages 1–30, 2–31, …, 85–114) and combined their outputs through a derivative-threshold filtering method. Rather than 85 independent models operating simultaneously, this ensemble employs consensus prediction where each 30-year window contributes only within its stable derivative range. For each sample, predictions from all models are generated, smoothed, and grouped into expert windows. Only regions where the derivative of predictions fell below a stability threshold were retained, effectively creating a voting mechanism where models with high confidence for a given sample determine the final biological age estimate (Figure 5a,b). This consensus strategy captures continuous aging dynamics while eliminating the need for chronological age as input, yielding a unified transcriptomic clock that adapts seamlessly across the lifespan.

**Figure 5.**
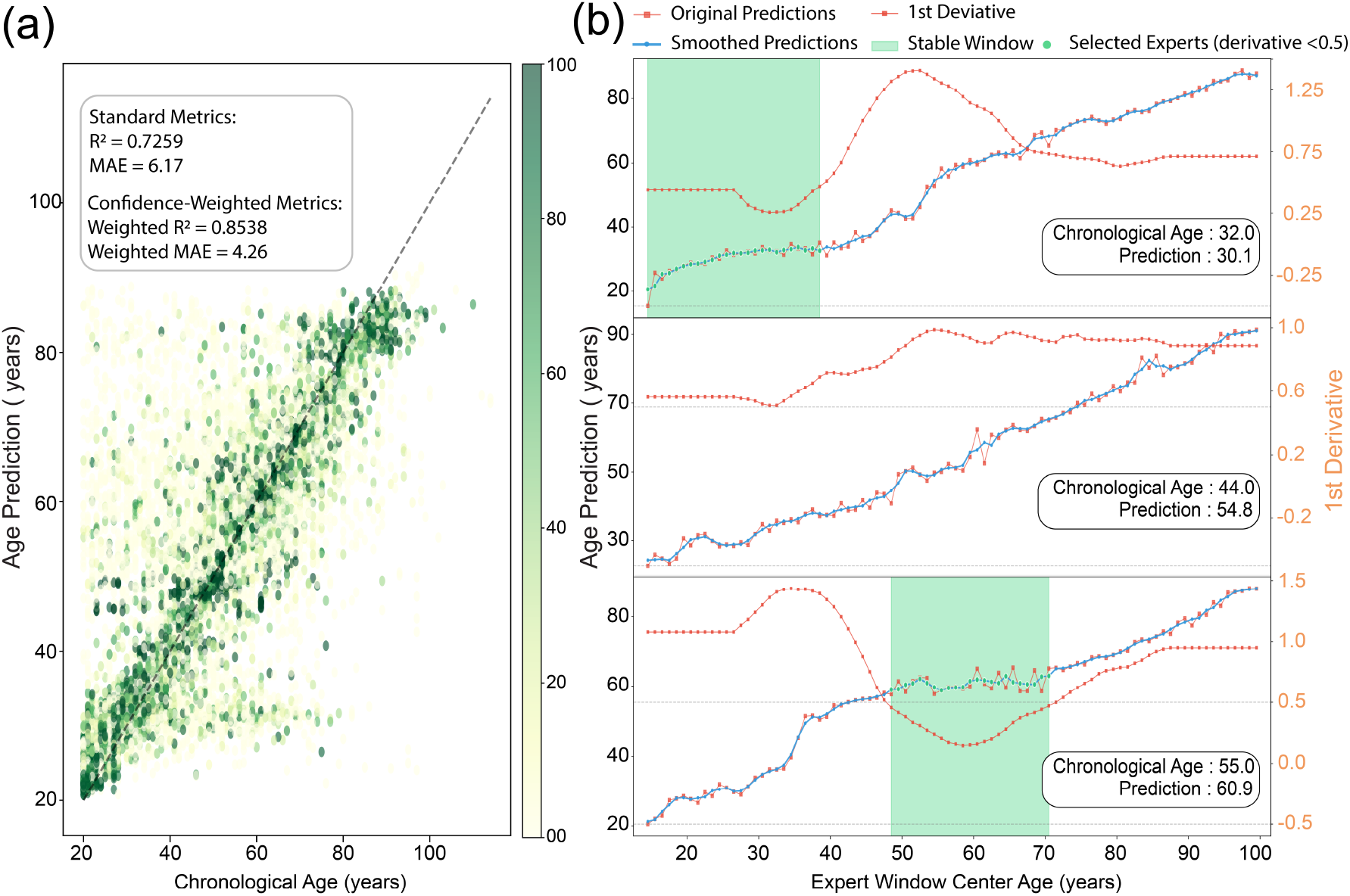
Final Ensemble Clock Performance. (a) Predicted versus chronological age for test samples (ages 20-80) colored by confidence scores. The ensemble achieves *R*^2^ = 0.726 and MAE = 6.17 years for all predictions, with confidence-weighted metrics showing improved performance (*R*^2^ = 0.854, MAE = 4.26 years). (b) Example predictions for three individuals showing how the derivative-threshold method selects stable expert windows. Top panel: 32-year-old with high-confidence prediction (30.1 years) based on stable expert consensus. Middle panel: 44-year-old with moderate confidence prediction (54.8 years) showing some model disagreement. Bottom panel: 55-year-old with high-confidence accurate prediction (60.9 years). Green shading indicates selected stable windows (derivative < 0.5), while the orange line shows the first derivative of smoothed predictions. Selected experts (green dots) contribute to the final prediction.

A key innovation of this framework is explicit **uncertainty quantification**. In machine learning more broadly, models that provide calibrated confidence estimates are considered safer and more reliable, particularly in high-stakes domains such as medicine [39, 40, 41]. Bringing this capability into biological clocks represents a conceptual advance: instead of offering only a point estimate, the model also reports how certain it is about that estimate.

The impact of uncertainty quantification is visible in Figure 5. In the aggregate scatterplot (Fig. 5a), samples with high confidence (green) fall tightly along the diagonal, while low-confidence samples deviate more strongly, especially at the extremes of the age range. Confidence weighting improved accuracy from *R*^2^ = 0.726 (MAE = 6.17) to *R*^2^ = 0.854 (MAE = 4.26), demonstrating that uncertainty itself is predictive of reliability. The three individual examples (Fig. 5b) illustrate this clearly: a 32-year-old is predicted almost exactly (30.1 years) with high confidence based on stable windows; a 44-year-old shows no stable regions, leading to an inaccurate prediction (54.8 years) that is correctly flagged as low confidence; and a 55-year-old again achieves an accurate, high-confidence prediction (60.9 years). These examples highlight that the model “knows when it does not know,” a crucial property for clinical deployment.

In our context, uncertainty quantification serves both as a safeguard and as a biological signal. It provides a quality measure for each prediction, flagging when outputs should be interpreted cautiously, while also revealing biological heterogeneity: low-confidence predictions are enriched in transitional phases and in extreme ages where transcriptomic variability is highest. Thus, the unified ensemble clock is not only more accurate but also self-aware, offering calibrated predictions that integrate seamlessly into clinical or research settings.

### Discussion

This work demonstrates how human-AI collaboration using advanced foundation models like Gemini 2.5 Pro can compress research timelines from months to weeks, with AI systems rapidly implementing and testing researcher-formulated hypotheses while identifying patterns that inform subsequent human decisions, from literature synthesis and hypothesis generation to modeling, interpretation, and documentation, under human supervision. Unlike prior frameworks focused on isolated tasks, K-Dense was directed at an open-ended objective (“explore the relationship between aging and transcriptomics”) and, through iterative researcher guided refinement, produced methodological advances, state-of-the-art clocks, and biological insights. The system identified limitations in the MMA and, when prompted, proposed and implemented methodological alternatives including the sliding-window solution. This achievement, completed in weeks rather than the months typically required for such analyses, illustrates the acceleration possible through guided AI collaboration. The result is not yet an autonomous scientist, but a tightly coupled collaborator that shifts the bottleneck from computation to conceptual oversight. Researcher supervision grounded the system’s choices in biological plausibility and checked artifacts. This human-in-the-loop process is the practical near-term paradigm: automation explores broadly; human judgment decides what matters and why.

Our findings support a view of aging as a progressive, temporally structured process rather than a single uniform process. Wave-like, age-progressive gene-importance patterns (Fig. 3a) and stage-specific marker sets (MIR29B2CHG, RBP1, CHAMP1, SEPTIN3) align with modern frameworks that treat aging as coordinated shifts across hallmarks and pathways, not isolated damage events [42, 43]. Prior population-scale work has similarly reported nonlinear, stage-like changes in the plasma proteome distinct “waves” across life [44] and nonlinear multi-omics trajectories [45], consistent with our transcriptomic waves. At the same time, recurring signals such as CDKN2A/p16 and AMPD3 suggest continuous processes (senescence, metabolic stress) accumulating across decades. These observations are consistent with various hypotheses about aging mechanisms, ranging from purely stochastic damage accumulation [46, 43], loss of epigenetic information [47], integrative models combining stochastic damage with deterministic processes [48] to quasi-programmatic perspectives (e.g., hyperfunction or development-derived theories) that suggest age-linked biological activity patterns emerge from evolutionary constraints rather than direct genetic programming [49, 50].

Together, they offer a coherent picture: conserved temporal phases, stage-specific molecular drivers, and tissue-specific implementation. Our approach offers mechanistic advantages over methylation-based clocks, which function primarily as statistical correlates with CpG sites often lacking clear functional relationships to aging pathways. Gene expression directly reflects cellular function, providing interpretable insights into aging mechanisms and enabling realtime assessment of intervention efficacy. Blood and colon each recapitulated the same temporal wave structure but with largely non-overlapping genes (Fig. 4b), indicating a systems-level timing with organ-specific execution. This duality explains why single-tissue clocks often generalize poorly and underscores the need for tissue-aware biomarkers that still respect shared temporal phases.

Three technical innovations enhanced model performance: density-weighted regression addressed ARCHS4’s bimodal age distribution through inverse probability weighting, heteroscedastic uncertainty quantification captured increasing transcriptomic variability with age, and the XGBoost architecture itself may have biological relevance, its cascading decision trees parallel how aging unfolds through sequential molecular events, with thresholds naturally capturing phenomena like senescence activation when damage exceeds repair capacity.

Our unified ensemble provides explicit uncertainty quantification (UQ). In ML for medicine, calibrated uncertainty is a prerequisite for reliability and safe deployment (e.g., Bayesian approaches, deep ensembles, conformal prediction) [39, 40, 41, 51, 52]. In Fig. 5a–b, high-confidence predictions cluster on the diagonal and low-confidence cases flag transitional/extreme ages, periods of higher heterogeneity. Confidence-weighted evaluation improves performance (*R*^2^ from 0.726 to 0.854; MAE from 6.17 to 4.26), and UQ itself becomes a biological signal (where the clock is least certain, biology is most variable).

The ensemble clock removes the need to input chronological age, provides point estimates with calibrated confidence, and operates on accessible tissues (e.g., blood), making it practical for longitudinal healthspan monitoring. In the broader biomarker landscape, clinically oriented clocks have emphasized outcomes and pace (e.g., PhenoAge, Grim-Age, DunedinPACE) [11, 12, 53]; our contribution is a transcriptome-based approach that is both competitive and interpretable, with UQ to flag low-reliability cases, an attribute valued in clinical AI [51, 52]. These properties support nearterm use in intervention studies and real-world monitoring, with UQ guiding confirmatory testing or orthogonal assays.

### Limitations and Future Directions

Feature importance is correlative, not causal; highlighted markers may track aging without driving it. While ARCHS4 provides de facto cross-study validation across 1,039 independent datasets, systematic biases remain, underscoring the need for validation in independent cohorts. The dataset introduces sampling biases (e.g., a bimodal age distribution with peaks at 21 and 71 years), and per-tissue sliding-window analyses require sufficient coverage, which restricted detailed maps to blood and colon. The stage-specific MMA requires chronological age for model selection; our attempts to learn a transcriptome-only gating function were unsuccessful, motivating the age-agnostic ensemble presented here. Future work should test generalization under distribution shift and calibrate UQ across independent cohorts (an established concern in clinical ML [41, 51]).

Three directions follow naturally. (i) Mechanism: integrate multi-omics (epigenome, proteome, metabolome) to map how temporal transcriptomic processes couple to hallmarks and pathways [43, 45]. (ii) Causality: pair clocks with perturbation/longitudinal designs to distinguish correlates from drivers and to test whether stage-specific markers predict intervention responses (e.g., senescence-targeted therapies in late life). (iii) Translation: prospective studies in clinical settings, using UQ to gate decisions (e.g., lowconfidence predictions trigger confirmatory tests), analogous to best practices for trustworthy clinical AI [41, 51].

### Conclusion

By directing a guided AI scientist system at an open-ended objective, we delivered transcriptomic clocks that match state-of-the-art accuracy while revealing how aging proceeds through conserved temporal phases executed via tissue-specific process. The unified, uncertainty-aware ensemble turns a clock into a reliable instrument, one that knows when its predictions warrant caution, bridging discovery and deployment. The K-Dense approach demonstrated here could accelerate discovery in any data-rich domain with established analytical frameworks, from molecular biology to climate science, while maintaining human oversight. Extending this framework to multi-omics, longitudinal cohorts, and perturbation studies will deepen mechanism and accelerate translation toward interventions that extend healthspan.

## Methods

### Data Source and Inclusion Criteria

We used the ARCHS4 database [22], which contains uniformly reprocessed RNA-seq data from over 10,000 GEO studies. From this resource, we selected 1,039 human studies that met the following criteria: (i) valid chronological age annotations, (ii) minimum sequencing depth of 10 million reads, (iii) RIN scores ≥ 6 when available, (iv) clear tissue/organ annotations, and (v) no cell line or in vitro samples. Disease samples were excluded if the condition directly confounded aging (e.g., cancers), but healthy controls from such studies were retained. The final dataset comprised 57,584 samples spanning ages 1–114 years across 28 tissues. While ARCHS4 applies uniform computational processing via Kallisto alignment, pre-analytical variations (tissue collection, RNA extraction, library preparation, sequencing platforms) remain intact across studies, providing de facto biological and technical diversity. For analysis, the 5,000 most variable genes (by variance across all samples) were selected.

### Quality Control

Age values encoded as ranges were converted to midpoints. Samples with missing or ambiguous annotations were removed. Expression data were normalized to 10,000 counts per sample and log_1*p*_-transformed. Technical validation confirmed consistent coverage across genes and tissues (Supplementary Fig. S3).

### Multi-Model Architecture (MMA)

To test the hypothesis that aging is stage-specific, we trained four age-stratified XGBoost models covering 1–30, 30–50, 50–70, and 70+ years. These boundaries were deliberately kept simple and not optimized, as our focus was demonstrating feasibility of age-stratified modeling. Models were trained independently with the following hyperparameters (optimized by 5-fold cross-validation within each window): n_estimators = 300 (early stopping at 250-280 rounds), max_depth = 8, learning_rate = 0.05, subsample = 0.8, colsample_bytree = 0.8, objective = ‘reg:squarederror’, tree_method = ‘hist’, random_state = 42. Each model was evaluated on held-out test data (80/20 split with age stratification), with ensemble predictions obtained by combining outputs across windows.

### Sliding-Window Analysis

To capture continuous dynamics, we trained 85 overlapping models spanning 30-year age windows advancing in 1-year increments (1–30, 2–31, …, 85–114). Windows with fewer than 100 samples were excluded. For each window, feature importance was calculated using the XGBoost gain metric. Genes were ordered by their weighted peak importance age—calculated as the weighted average of windows where each gene exceeded 30% of its maximum importance—with similar-peaking genes clustered using hierarchical clustering to preserve pattern continuity. The complete gene ordering data for both pan-tissue (Figure∼3a) and tissue-specific (Figure∼4b) analyses are provided in Supplementary Data files `gene_ordering_AllTissues_data_sliding_window.csv, gene_ordering_Onlyblood_data_sliding_window.csv, gene_ordering_Onlycolon_data_sliding_window.csv,` and `gene_ordering_blood&colon_data_sliding_window.csv`. This produced the wave-like patterns shown in Figure 3a.

Pathway enrichment analysis integrated Gene Ontology, KEGG, Reactome, WikiPathways, and aging-related gene sets. Enrichment scores combined gene counts, importance values, expression changes, and *p*-values, with database-specific weights (1.5× for aging-related databases, 1.2× for KEGG/Reactome, 1.0× for GO).

### Ensemble Clock and Uncertainty Quantification

We developed an ensemble prediction framework integrating all 85 models. For each sample, predictions were smoothed using a 3-point moving average, and local derivatives across consecutive models were computed using linear regression on 25-model windows. Windows with absolute derivative *<* 0.5 were deemed stable; predictions from these “stable experts” were averaged. Confidence scores combined the number of contributing experts and the derivative magnitude:

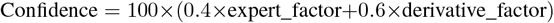

where expert_factor uses exponential scaling (1 − *e*^−0.08*n*^) for *n* stable experts, and derivative_factor linearly maps derivatives from 0.25 (maximum confidence) to 0.5 (minimum confidence). Calibration was assessed using reliability diagrams, yielding a mean calibration error of 0.7%. Confidence weighting improved accuracy from *R*^2^ = 0.726 (MAE = 6.17) to *R*^2^ = 0.854 (MAE = 4.26).

### Tissue-Specific Analysis

We screened 28 tissues for sample size and age coverage (Supplementary Data). Because sliding-window modeling requires sufficient representation across all age windows, only blood (*n* ≈ 4, 285) and colon (*n* ≈ 1, 254) met this criterion. Sliding-window feature importance analysis was therefore restricted to these two tissues, with cross-tissue comparisons focusing on temporal trajectories versus gene identity (Fig. 4b). Cross-tissue comparison employed identical hyperparameters to ensure comparability.

### Statistical Analysis

Performance was assessed using coefficient of determination (*R*^2^), mean absolute error (MAE), and root mean squared error (RMSE). Bootstrap confidence intervals (1,000 iterations) were calculated for all metrics. Cross-validation used 5-fold stratified splits by age distribution, and study-level validation used leave-one-study-out analysis to test robustness against batch effects.

### Human–AI Interaction Protocol

K-Dense, powered by Google’s Gemini 2.5 Pro with its 2M token context window and advanced reasoning capabilities, was provided with the directive: “Explore the relationship between aging and transcriptomics.” The system iteratively generated hypotheses, modeling strategies, and interpretations, which were reviewed and refined under PI supervision (Figure 1b). Modules operate through specialized components: Literature Reviewer (knowledge synthesis), Research Planner (hypothesis formulation), Coder & Analyst (computational implementation), and Writer & Summarizer (documentation generation), with convergence determined by performance metrics or iteration limits (maximum 50 cycles). Prompts, intermediate outputs, and researcher annotations are archived in Supplementary Note 1 to ensure reproducibility of the AI-guided workflow.

### Computational Implementation

Analyses used Python 3.9.12 with scikit-learn 1.1.1, xgboost 1.6.1, numpy 1.22.3, pandas 1.4.2, and scanpy 1.9.1. Computations utilized Intel Xeon Gold 6248R (48 cores) with 384GB RAM. GPU acceleration (NVIDIA A100) was employed for model training. Total computational time was approximately 5 hours on CPU or 1 hour with GPU acceleration.

## Supporting information

Data files for aging clock analysis & manuscripts.

## Code and Data Availability

All analysis code, including K-Dense orchestration scripts, trained models, Processed feature matrices and prediction outputs will be made available at https://github.com/biostateAI/Longevity_Paper. The ARCHS4 database is publicly available at GEO.

## Data availability

The ARCHS4 database is publicly available via Gene Expression Omnibus. Analysis code generated by K-Dense system, including XGBoost sliding window analysis code as well as processed feature matrices, trained models, Raw predictions and feature importance matrices will be available at https://github.com/biostateAI/Longevity_Paperuponpublication.

## Author contributions

A.G. conceived the project. V.A. performed the majority of interactions with the K-Dense system and conducted the computational analyses. O.L., T.K., V.A., and A.G. developed and implemented the K-Dense multi-agent framework. D.A.S., C.A.P., P.R., A.G., and V.A. interpreted the biological findings and contextualized results within the aging biology framework. A.G. wrote the manuscript with input from all authors.

## Supplementary Information

### Supplementary Note 1: K-Dense AI Scientist System Architecture

The K-Dense (Knowledge Dense) AI Scientist represents a comprehensive multi-agent research framework that orchestrates the complete scientific discovery workflow, from literature synthesis through computational analysis to manuscript generation. Built on Google’s Gemini 2.5 Pro foundation model, the system leverages advanced reasoning capabilities, multi-modal understanding, and large context windows to enable sophisticated scientific reasoning and analysis. The architecture transforms researcher intent into complete, publication-ready scientific manuscripts through coordinated phases of evidence building, computational analysis, and narrative assembly.

The system consists of four specialized modules operating in iterative, reflexive cycles: a **Literature Reviewer** that conducts comprehensive literature surveys and identifies knowledge gaps; a **Research Planner** that converts insights into testable hypotheses and analysis strategies; a **Coder & Analyst** (implemented as DendroForge) that executes computations in sandboxed environments with rigorous validation; and a **Writer & Summarizer** that consolidates all materials into polished documents with consistent style, citations, and figures. Each module exposes structured inputs and outputs enabling the system to loop, branch, or re-run specific stages without destabilizing the end-to-end workflow.

### System-Level Pseudocode

The complete K-Dense workflow can be represented as:

#### ALGORITHM 1: K-Dense AI Scientist System

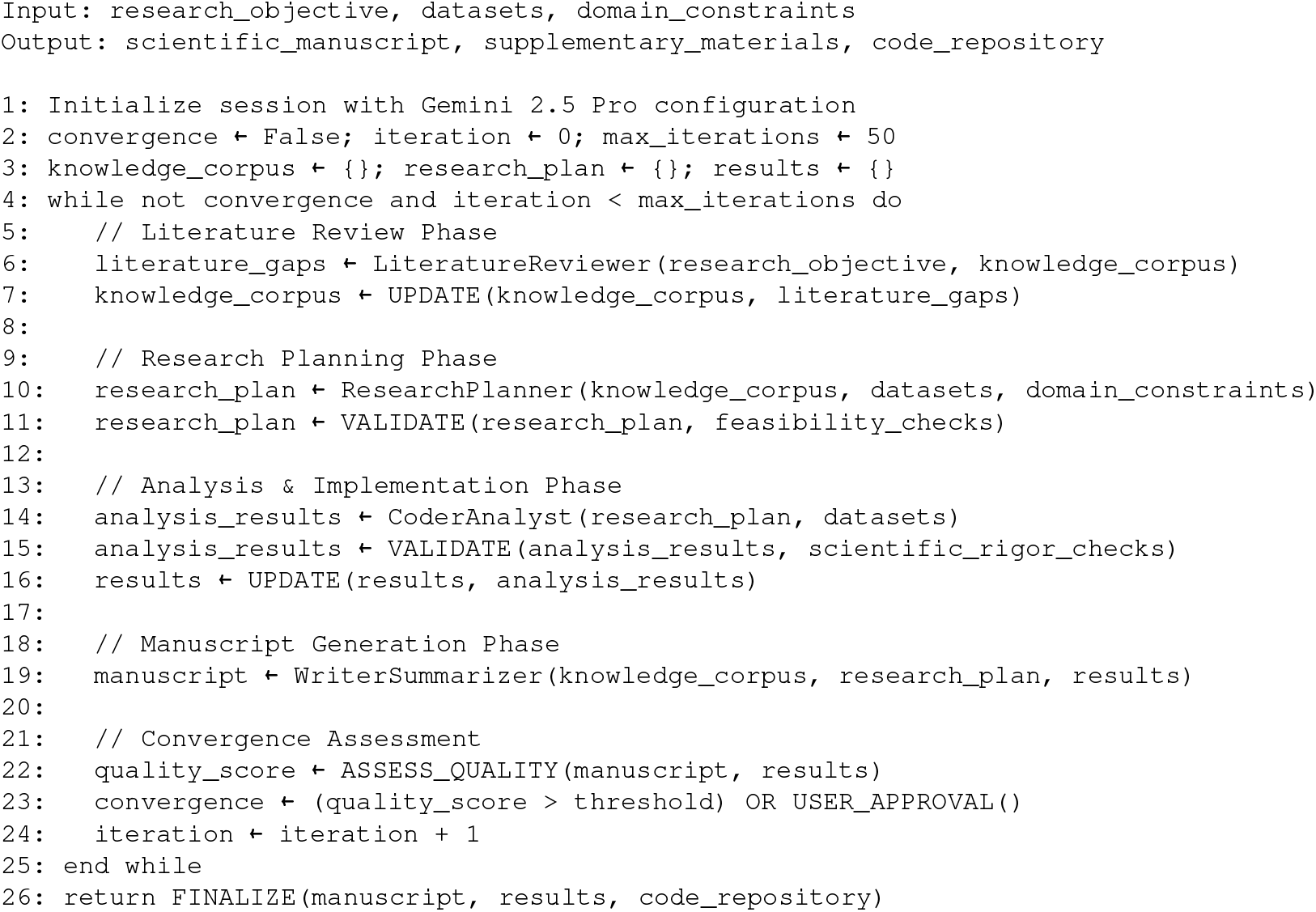

#### Module 1: Literature Reviewer

The Literature Reviewer module produces comprehensive literature review manuscripts through a complete execution of the paper writing pipeline (paper_writing.py), powered by Gemini 2.5 Pro’s advanced reasoning and multi-modal capabilities. Taking a data report and main research goals as primary inputs, the module generates a fully-formed literature review paper in Markdown format with properly formatted citations and comprehensive coverage of the research domain.

The module operates through the complete paper writing workflow: initial research planning generates hierarchical topic outlines based on the data characteristics and research objectives; parallel research agents conduct extensive literature searches using multiple academic databases and web sources; advanced citation management systems extract, validate, and format references across multiple citation styles; and the integrated writing system synthesizes findings into coherent narrative sections following academic conventions. The output is a complete literature review manuscript that contextualizes the research objectives within existing knowledge, identifies key gaps and controversies, and establishes the theoretical foundation for subsequent analytical work.

### Literature Reviewer Pseudocode

#### ALGORITHM 2: Literature Reviewer Module

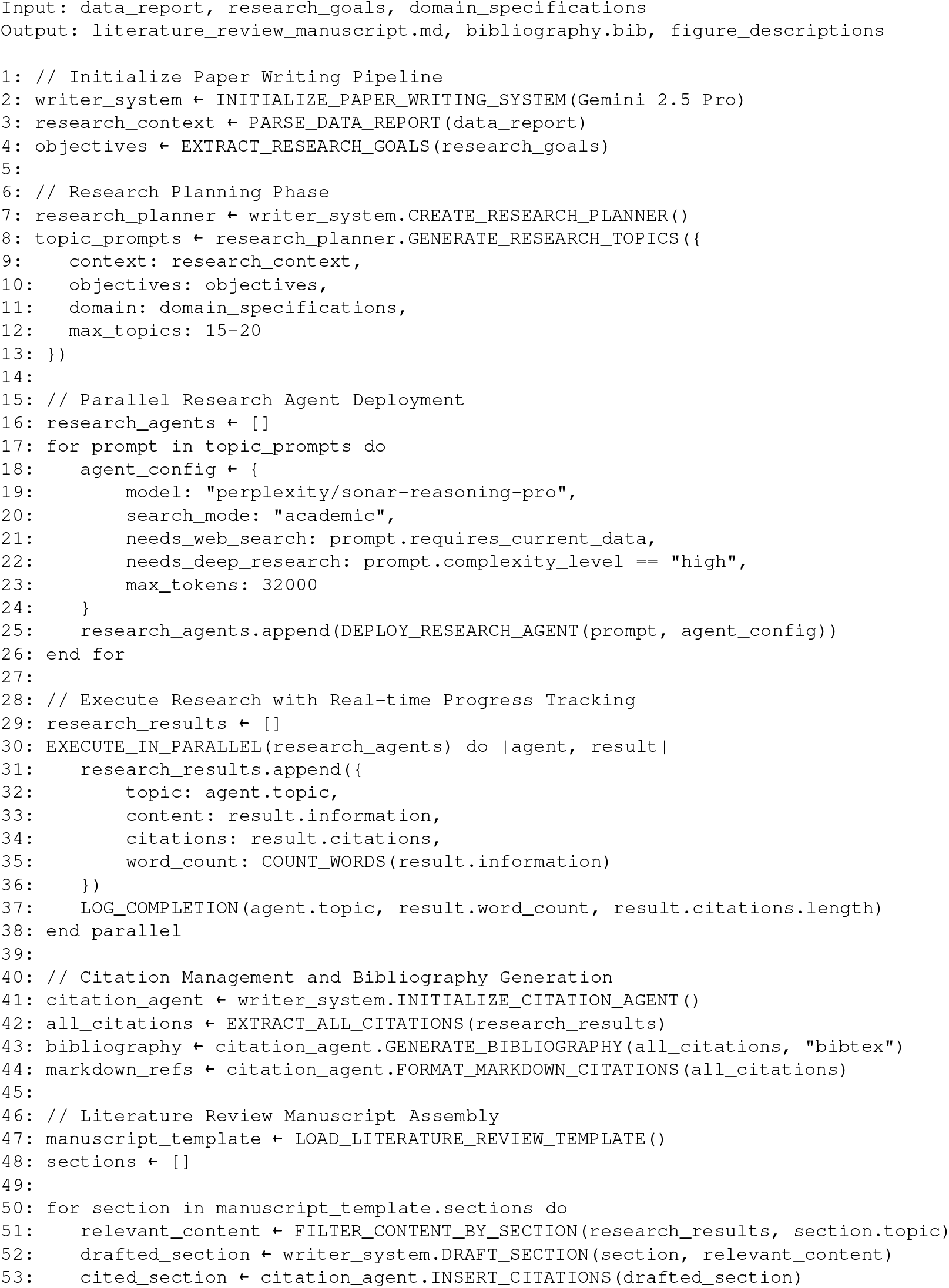
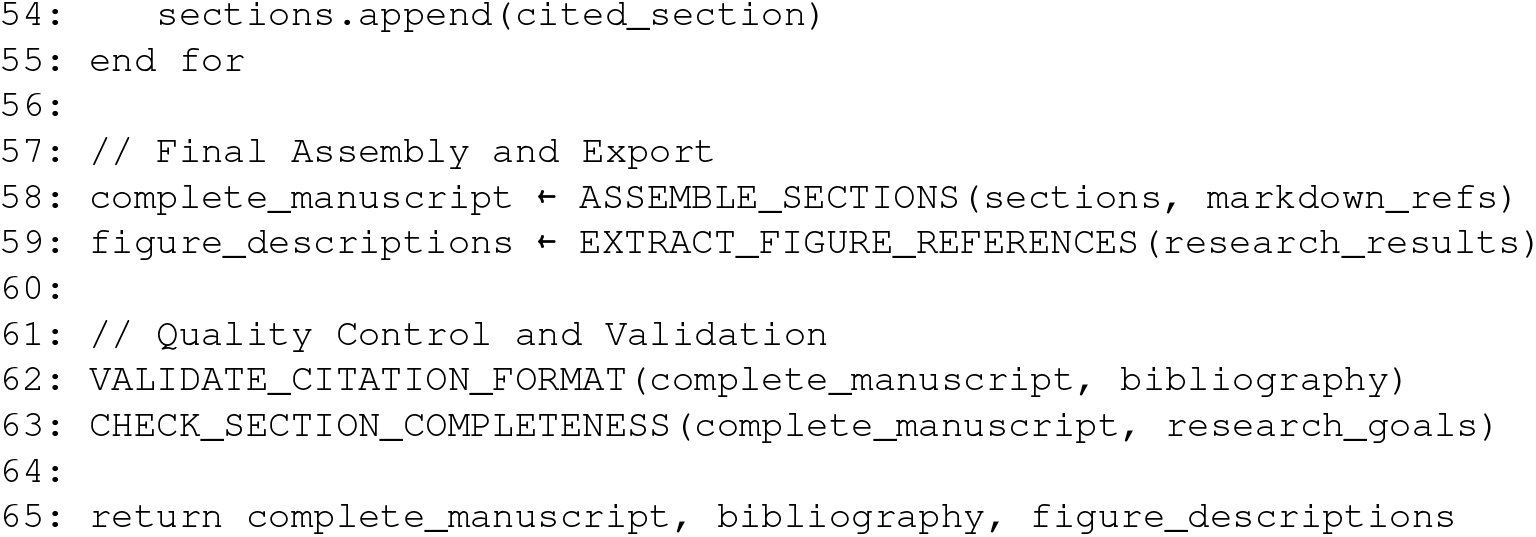

#### Module 2: Research Planner

As a core component of the K-Dense Analyst system, the Research Planner employs chain of thought reasoning and deep thinking processes to transform literature insights into executable research strategies through iterative hypothesis generation and methodological refinement. Built on Gemini 2.5 Pro’s advanced planning capabilities, the module performs systematic analysis of input data and literature review papers to identify knowledge gaps, methodological limitations in prior work, and opportunities for novel analytical approaches. Using structured reasoning chains, the planner generates an initial comprehensive research plan with testable hypotheses, explicit success criteria, statistical power requirements, and resource constraints.

The module creates comprehensive research plans encompassing data preprocessing pipelines, statistical analysis strategies, validation protocols, and expected outputs. Plans are subjected to automated feasibility assessment considering available datasets, computational resources, and methodological constraints. The planner incorporates domain-specific best practices for experimental design, statistical rigor, and reproducibility standards. Each plan component includes detailed specifications for required inputs, intermediate artifacts, success metrics, and potential failure modes. The module supports plan versioning and iterative refinement based on implementation feedback, ensuring adaptive responses to unexpected analytical challenges.

This is the module that the overall review and reflection agents give feedback to so that relevant changes can be made to the plan to come up with more novel ideas and corrected robust hypothesis testing and research execution

#### Research Planner Pseudocode

##### ALGORITHM 2.5: Research Planner Module

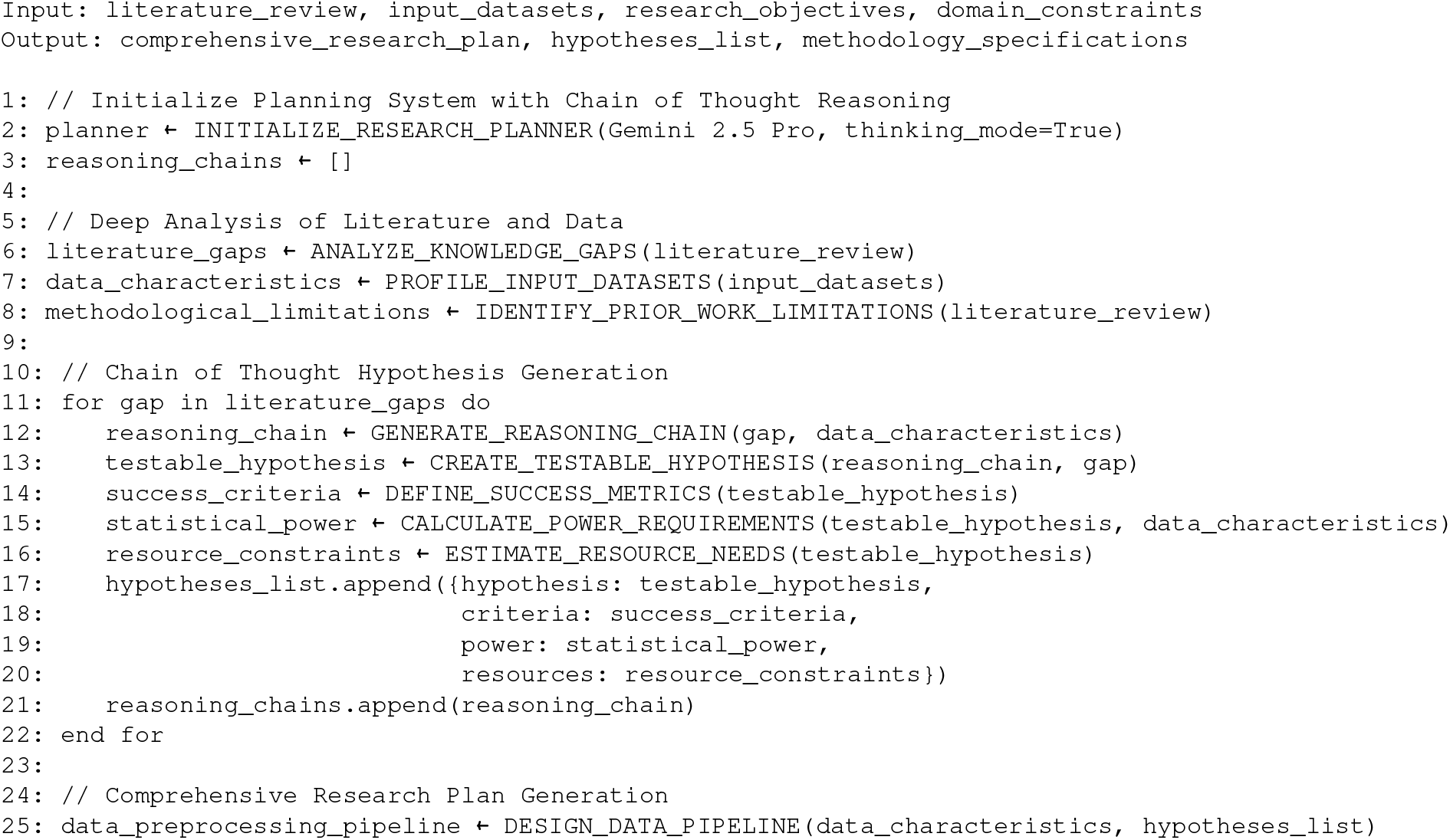
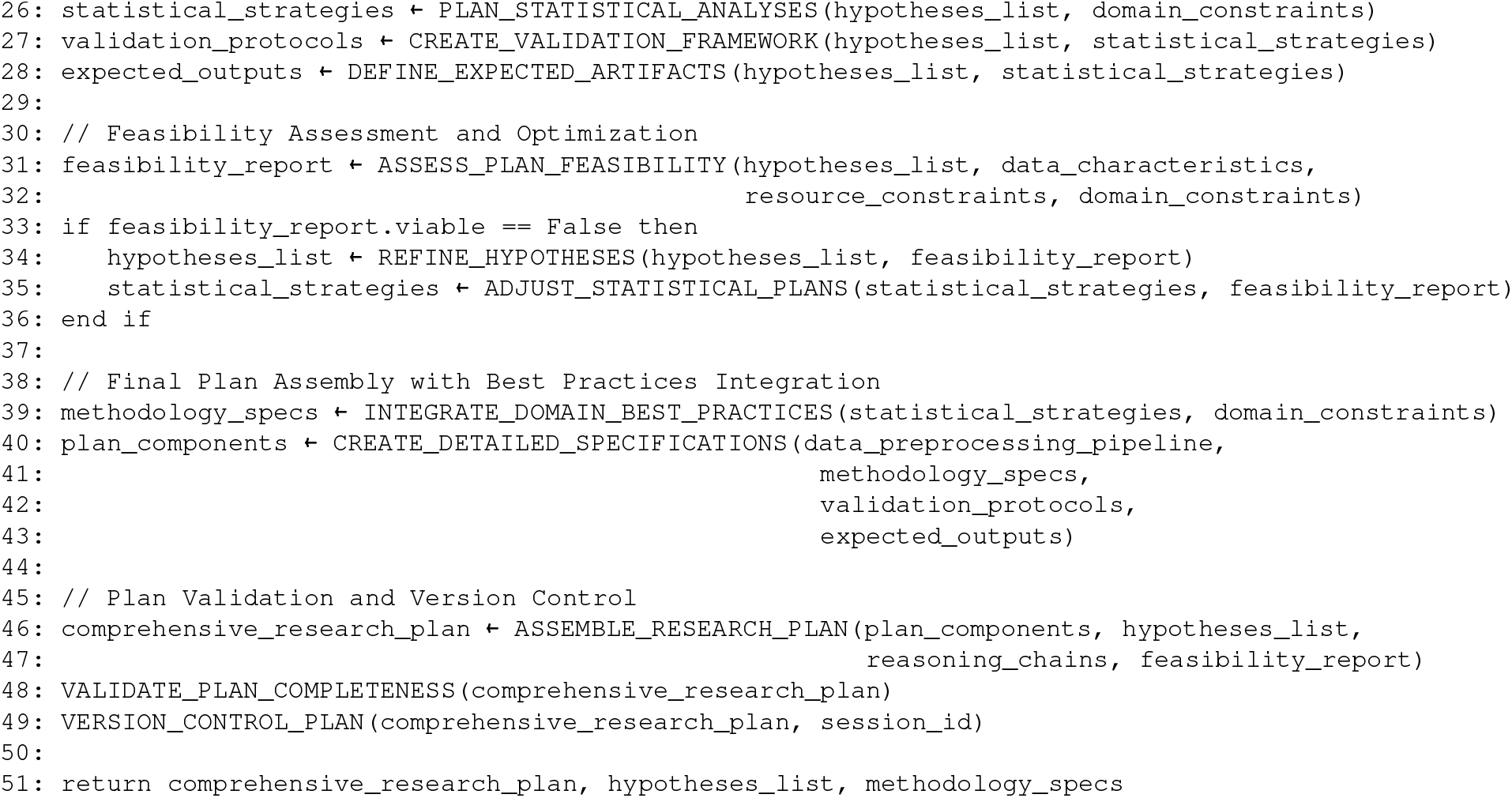

#### Module 3: Coder & Analyst (DendroForge)

The computational core of K-Dense is implemented through DendroForge, a sophisticated multi-agent analysis system that transforms research plans into validated scientific results. Operating in secure sandbox environments with comprehensive security boundaries, DendroForge consists of five specialized sub-agents orchestrated through structured workflows:

##### Planning Agent

Decomposes research objectives into executable computational tasks, mapping dependencies and identifying parallelization opportunities. Creates detailed implementation specifications with checkpoint validation and rollback procedures.

##### Coding Agent

Implements analysis pipelines using Python 3.11+ with standardized bioinformatics libraries (scanpy, pandas, scikit-learn, etc.). All code follows reproducibility standards with explicit random seeds, version documentation, and automated error handling for common failure modes.

##### Review Agent

Conducts detailed code review examining implementation correctness, statistical appropriateness, and computational efficiency. Validates that metrics are computed as intended and checks alignment with research questions.

##### Science Methodology Agent

Evaluates methodological soundness including appropriateness of statistical tests, adequacy of controls, power considerations, and biological interpretation of results. Ensures alignment with domain-specific best practices.

##### Feedback Summary Agent

Synthesizes review outputs to determine implementation completeness, identifies remaining issues, and coordinates iterative refinement cycles. Implements loop detection to prevent infinite revision cycles.

The system maintains comprehensive execution logs, intermediate artifacts, and validation checkpoints throughout the analysis workflow, enabling full audit trails and reproducible execution.

### Coder & Analyst (K-Dense Analyst) Pseudocode

#### ALGORITHM 3: Coder & Analyst Module (DendroForge)

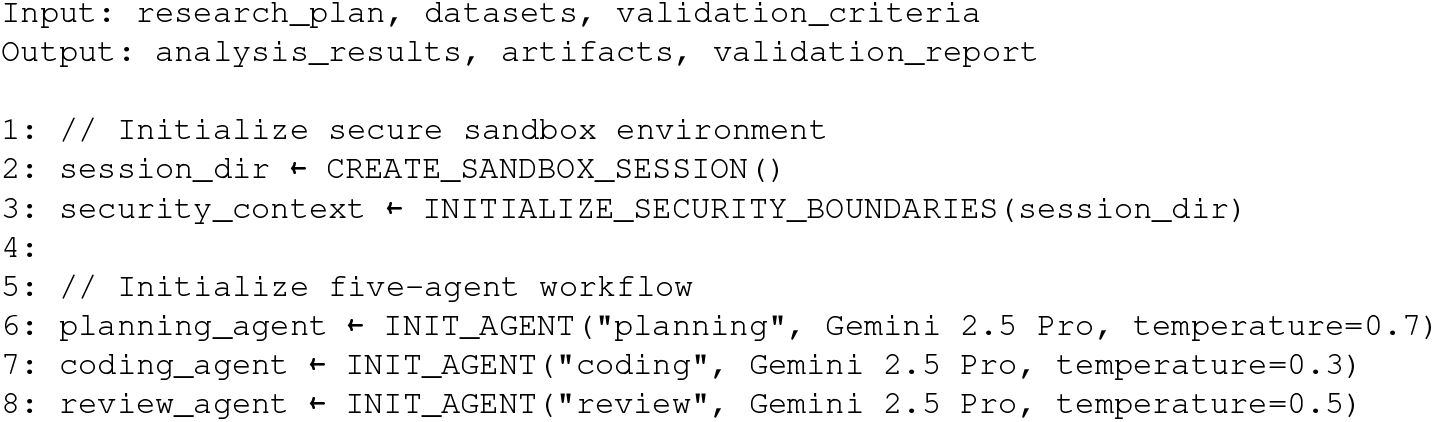
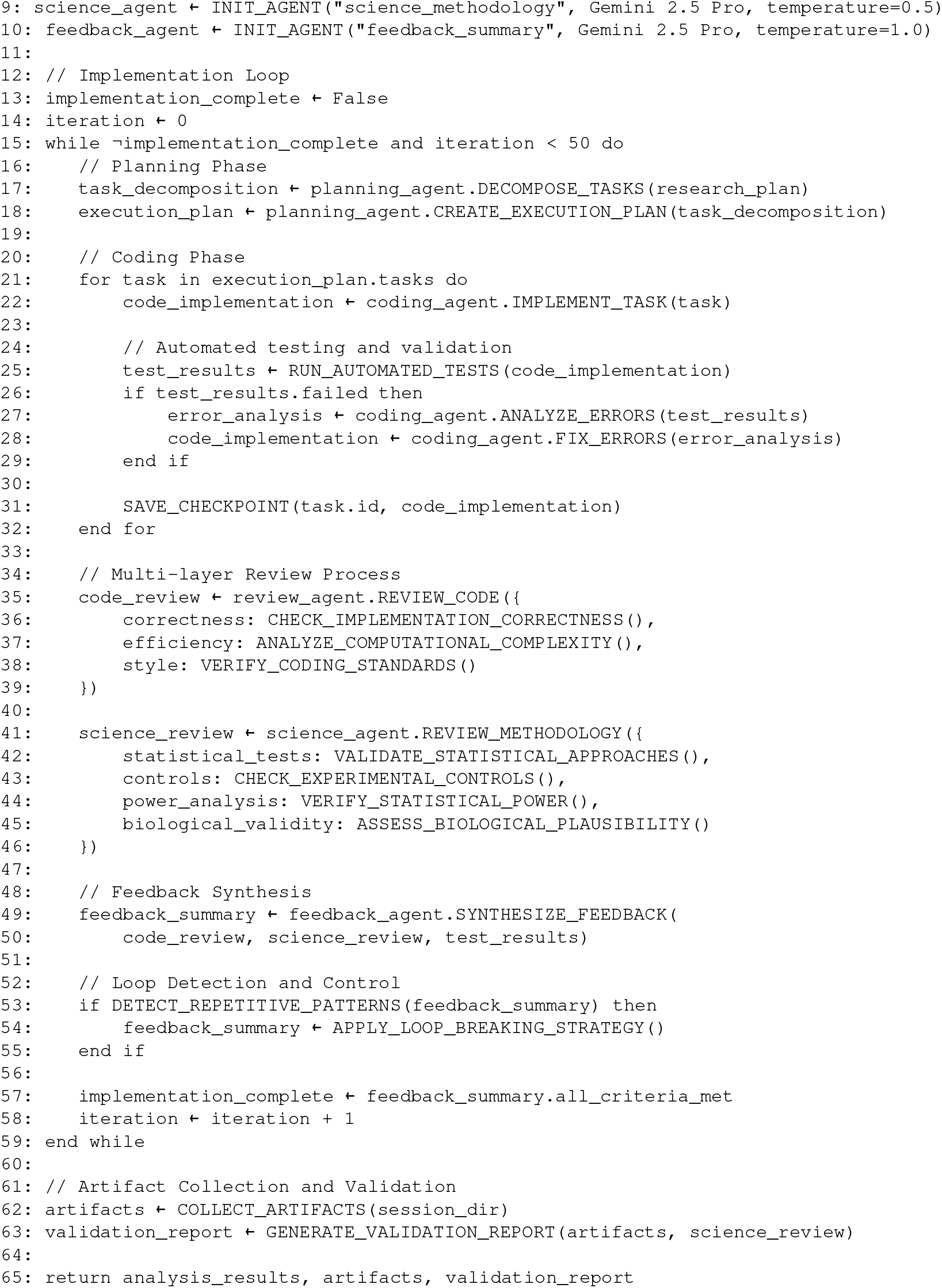

#### Module 4: Writer & Summarizer

The Writer & Summarizer module consolidates the complete research workflow into publication-ready scientific manuscripts. Leveraging Gemini 2.5 Pro’s advanced language capabilities, the module integrates literature corpus, research plans, and analytical results into coherent narratives following field-specific conventions and journal standards.

The module operates through sophisticated document orchestration encompassing multi-modal content integration, automated citation management, and intelligent figure processing. Advanced image analysis capabilities convert scientific figures into detailed textual descriptions, enabling comprehensive figure caption generation and cross-referencing. The citation system performs automated reference extraction from multiple sources, deduplication across format variations, and bibliography generation in both Markdown and BibTeX formats.

The writing process implements section-level iterative refinement with gap-filling mechanisms that identify missing evidence and perform targeted literature searches to supply supporting excerpts. A document orchestrator maintains conceptual consistency across sections while enforcing venue-specific formatting requirements. The module supports multiple output formats including journal-specific templates, preprint servers, and supplementary documentation packages.

### Final Paper Writer Pseudocode

#### ALGORITHM 4: Final Paper Writer Module

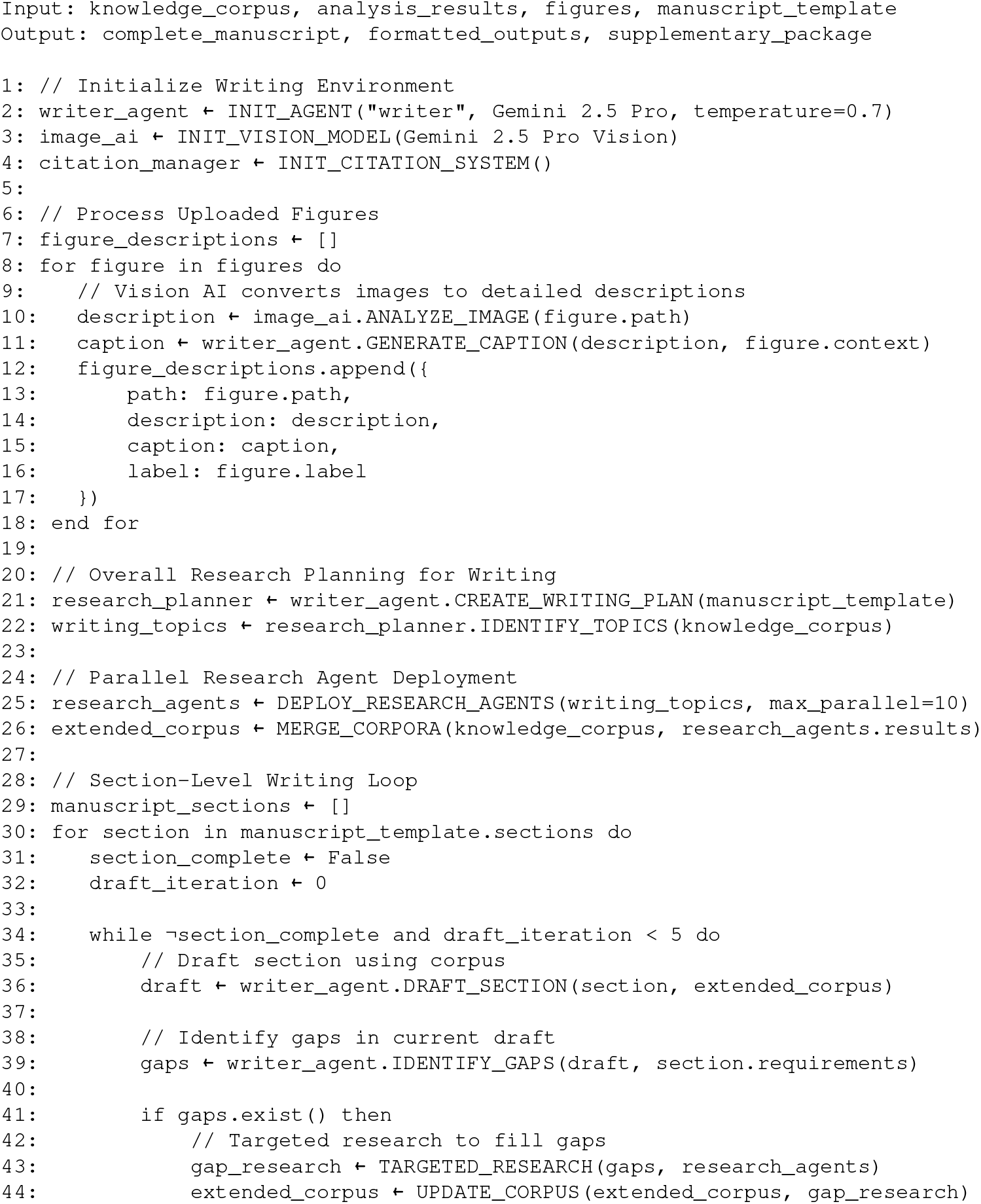
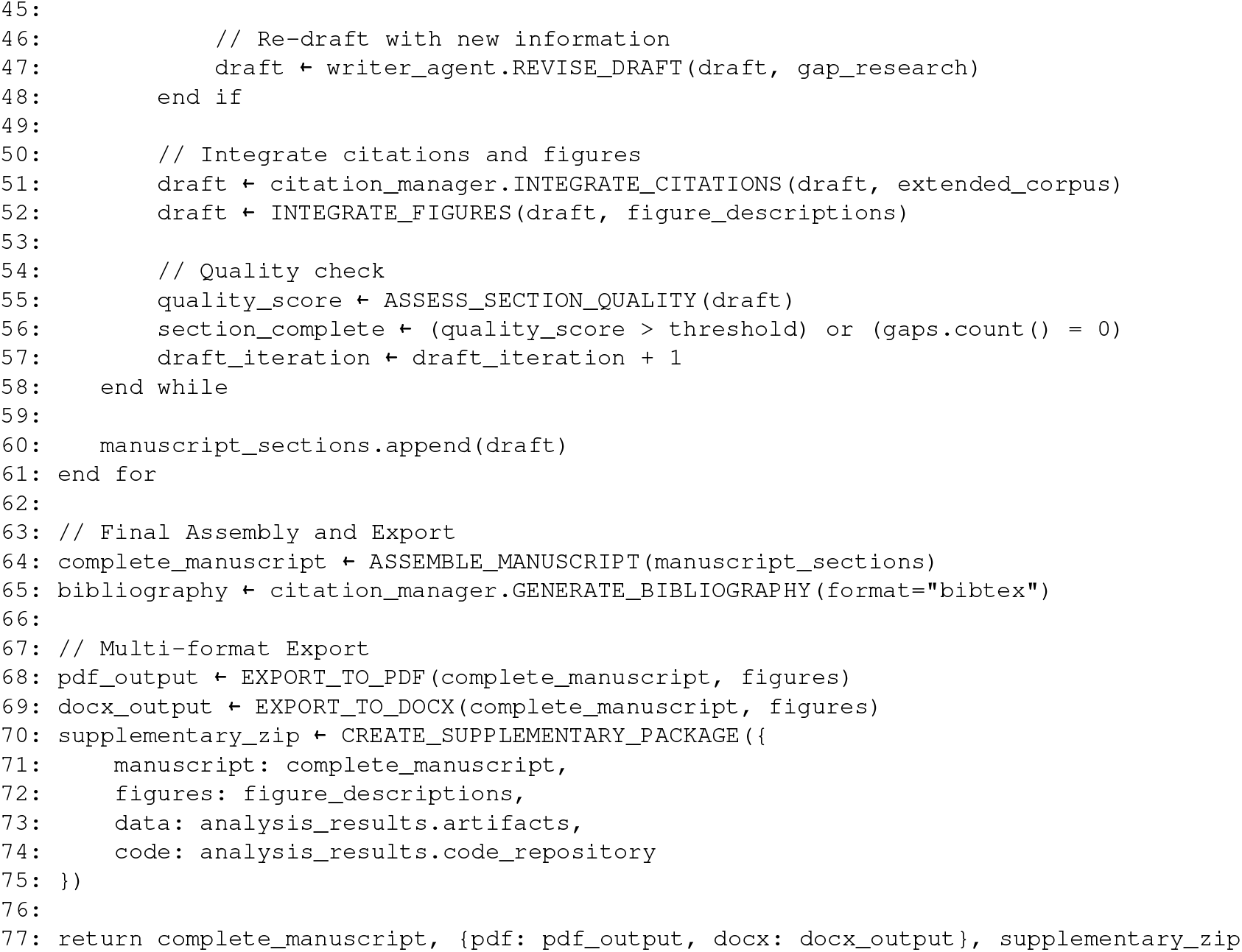

#### Section drafting and gap closure

The Writer operates an inner loop at the section level. A drafting process produces paragraphs from the outline and corpus; a gap-filling process identifies missing evidence (e.g., mechanistic rationales, quantitative baselines, or formal statements of assumptions) and performs focused lookups to supply concise excerpts; and a citation/figure integrator grounds claims with sources, resolves numbering across sections, and finalizes cross-references to figures, tables, and supplementary materials. A writer orchestrator enforces section-specific style guidance (tone, tense, and level of technicality), maintains a concept map of key entities, and ensures that all nontrivial assertions trace back to items in the corpus. A bibliography manager deduplicates references, harmonizes citation keys across sections, and emits a single manifest that later travels with the analysis outputs. At this stage the document is a well-scoped narrative scaffold with an auditable body of evidence but without computed results.

#### Inter-Agent Communication and Data Flow

The K-Dense system implements sophisticated inter-agent coordination through structured data exchange protocols and shared state management. Each module maintains versioned interfaces enabling seamless handoffs while preserving data integrity and provenance tracking.

#### Connection Architecture

The four modules are connected through a hierarchical message-passing system with the following data flow patterns:

##### Literature Reviewer → Research Planner

A complete literature review manuscript in Markdown format, comprehensive bibliography, and figure descriptions are passed along with extracted knowledge gaps and research opportunities. This enables the Research Planner to generate hypotheses grounded in comprehensive literature analysis and identify methodological approaches that build upon existing work.

##### Research Planner → Coder & Analyst

Research plans are transmitted as executable specifications containing task decompositions, success criteria, validation protocols, and resource constraints. Each plan component includes typed inputs/outputs enabling the DendroForge system to implement analysis pipelines systematically.

##### Coder & Analyst → Writer & Summarizer

Analysis results, validated artifacts, and execution logs are packaged with comprehensive metadata including provenance chains, validation scores, and intermediate checkpoints. This enables the

Writer to accurately contextualize findings within the narrative framework.

##### Bidirectional Feedback Loops

All modules support backward communication for iterative refinement. The Writer can request additional literature review for gap-filling; the Analyst can query the Planner for clarification on ambiguous specifications; and the Planner can request re-analysis based on Writer-identified narrative requirements.

#### Shared State Management

The system maintains global state through: - **Session Context**: Thread-safe storage of session identifiers, file paths, and security boundaries - **Knowledge Graph**: Semantic network of concepts, relationships, and evidence maintained across modules - **Artifact Registry**: Centralized tracking of all generated files, figures, and intermediate outputs - **Provenance Chain**: Immutable audit trail linking every output to its generating agent and input data

#### Handoff to analysis

The transition from Writer to Analyst is governed by a typed specification that captures: (i) the scientific questions to be answered, (ii) data dependencies and permissible transformations, (iii) success criteria and evaluation metrics, (iv) figure/table intents (what each result should communicate), and (v) any ethical, biosafety, or dataset-governance constraints. This specification is versioned and appended to the provenance log. It allows the Analyst to operate independently while preserving alignment with the narrative goals established by the Writer.

#### K-Dense Analyst: planning loop

The Analyst converts the specification into an executable plan using a *planning loop*. An initial planning process defines the scope of computation, decomposes objectives into tasks, and maps each task to required inputs and expected artifacts. An orchestrator sequences these tasks, identifies opportunities for parallelism, and constructs checkpoints for partial validation. A planning review process audits coverage (e.g., controls, baselines, and ablation points), statistical appropriateness, and feasibility under available data and compute. Plans that fail review are revised until they meet adequacy thresholds, ensuring that only well-formed strategies enter execution.

#### K-Dense Analyst: implementation loop

Execution proceeds in a secure, sandboxed environment that isolates code, data, and artifacts for reproducibility. A coding planning process transforms the strategy into stepwise procedures—data loading, preprocessing, model fitting or statistical testing, uncertainty quantification, and visualization. A coding process executes these procedures with rich logging, structured error messages, and automatic retries for common failure modes (e.g., missing dependencies, malformed inputs, or numerical instability). Intermediate artifacts (cleaned datasets, parameter files, trained models, and diagnostic plots) are persisted at each checkpoint.

#### Validation and governance

Quality control is multi-layered. A coding review process examines implementation details, verifies that metrics are computed as intended, and checks that randomness is controlled via explicit seeds and documented environments. A science review process evaluates methodological soundness—appropriateness of controls, assumptions, and statistical tests; power considerations; and alignment with the research questions. A feedback summary process synthesizes these audits and determines whether to iterate the implementation loop, revise the plan, or proceed to reporting. All decisions, rationales, and diffs to the plan are recorded in the provenance log to support later audit or replication.

#### Return to Writer for presentation

Once the Analyst produces validated results, the system hands artifacts back to the Writer together with a structured *results manifest* (tables, figures, effect sizes, confidence intervals, and notes on limitations). **In contrast to the Literature Reviewer phase, the final paper writing stage requires all figures and the complete results report**. The Writer converts this into narrative text: results are contextualized against the earlier corpus, figure captions are tightened and aligned with the intended messages, and cross-references are inserted to methods and supplementary analyses. The Writer also harmonizes terminology introduced during analysis, updates the concept map, and ensures that any new claims are linked to either computed evidence or referenced literature. The final assembly step enforces venue-specific formatting, ensures section coherence, and exports the manuscript along with a complete supplementary package (text, figures, captions, references, and machine-readable manifests).

#### Reflexion and Multi-Agent Coordination

K-Dense implements sophisticated reflexion mechanisms enabling self-correction and iterative refinement across all modules. Each module incorporates multiple feedback loops: internal quality assessment, cross-module validation, and human-in-the-loop verification points. The system employs advanced prompt engineering with structured reasoning chains, multi-step validation protocols, and explicit uncertainty quantification to minimize hallucinations and ensure scientific accuracy.

Inter-module communication occurs through structured APIs with versioned data schemas, enabling robust handoffs and rollback capabilities. The architecture supports both breadth-first exploration (parallel hypothesis generation) and depth-first refinement (iterative methodology improvement). Module-specific reflexion mechanisms include literature consistency checking, plan feasibility validation, code correctness verification, and narrative coherence assessment.

#### System dynamics and stopping rules

The end-to-end system orchestrates dynamic workflows supporting adaptive research strategies. The Literature Reviewer establishes evidence foundations; the Research Planner generates testable hypotheses; the Coder & Analyst executes validated computational analysis; and the Writer & Summarizer synthesizes results into publication-ready narratives with full provenance tracking.

Convergence is determined through multi-criteria assessment: achievement of predefined performance thresholds (e.g., *R*^2^ *>* 0.85 for predictive models), successful completion of validation protocols, scientific rigor verification, and human investigator approval. The system supports maximum iteration limits (typically 50 cycles) with early termination for exceptional performance or explicit user satisfaction. Module idempotency and artifact versioning enable targeted re-execution (e.g., incorporating new data or addressing reviewer feedback) without destabilizing the complete pipeline.

#### Technical Implementation and Gemini 2.5 Pro Integration

The complete K-Dense system operates on Google’s Gemini 2.5 Pro foundation model, leveraging its 2M token context window, advanced reasoning capabilities, and multi-modal understanding for scientific analysis. The system implements sophisticated prompt engineering with structured reasoning templates, chain-of-thought guidance, and domain-specific scientific reasoning protocols optimized for Gemini’s architecture.

All modules utilize consistent model configurations with temperature controls adapted for specific tasks: creative hypothesis generation (temperature = 1.0), analytical reasoning (temperature = 0.7), and factual synthesis (temperature = 0.3). The system incorporates Gemini’s thinking tokens for explicit reasoning traces, enabling transparent decision processes and debugging capabilities.

#### Practical considerations

Throughout, the system emphasizes transparency and reproducibility through comprehensive artifact management. All intermediate outputs are versioned with cryptographic hashing; every figure and table maintains clear lineage from raw data through analysis to manuscript integration; and modular separation between planning, execution, and presentation enables targeted revisions without full pipeline re-execution.

The implemented workflow achieves scientific discovery acceleration through coordinated automation: the Literature Reviewer establishes comprehensive evidence foundations through systematic database queries and intelligent synthesis; the Research Planner transforms identified gaps into executable hypotheses with explicit validation criteria; the Coder & Analyst (DendroForge) implements rigorous computational analysis with multi-layered quality control; and the Writer & Summarizer produces publication-ready manuscripts with full provenance documentation. This integration demonstrates how advanced AI systems can compress traditional month-long research cycles into week-long collaborative investigations while maintaining scientific rigor and reproducibility standards.

### Supplementary Note 2: Prompting Methodology and Human-AI Interaction Protocol

The development workflow illustrated in Figure 1b demonstrates the iterative prompting strategy that guided K-Dense through the transcriptomic aging clock development. This process exemplifies how structured human intervention at critical decision points can direct AI systems toward methodological innovations while maintaining scientific rigor.

#### Initial Exploration Phase

The system was initialized with a deliberately open-ended directive: *“Explore the relationship between aging and transcriptomics using the ARCHS4 database*.*”* This broad framing allowed K-Dense to survey the landscape of available approaches while avoiding premature commitment to specific methodologies. The initial prompt included access to the full ARCHS4 dataset (57,584 samples) and standard machine learning libraries, with instructions to prioritize interpretability alongside predictive performance.

#### Iterative Hypothesis Testing

The orange-highlighted steps in Figure 1b mark critical intervention points where human judgment guided the system’s exploration. When K-Dense initially achieved moderate performance using a global XGBoost model (*R*^2^ = 0.619), systematic analysis revealed a fundamental roadblock: performance systematically deteriorated at higher ages, with mean absolute error increasing from 4 years in middle-aged samples to >12 years in elderly populations. The system’s residual analysis clearly demonstrated age-dependent heteroscedasticity, with prediction errors showing distinct clustering patterns and increasing variance envelopes at the extremes of the age distribution.

Recognizing this performance degradation as more than a technical limitation, researchers explicitly prompted the analyst: *“The current model shows systematic deterioration in older populations. Analyze this pattern and suggest methodological changes to handle age-dependent performance variation effectively in the next iteration*.*”* This directed the system to move beyond conventional solutions (feature selection, hyperparameter optimization) toward a fundamental reconsideration of model architecture.

The analyst’s response identified the age-stratified approach, leading to the pivotal follow-up prompt: *“Try separate models for different age ranges”* rather than pursuing traditional approaches. This intervention exemplifies the collaborative framework: the AI system identified the performance limitation and characterized the error patterns, while human expertise recognized that the underlying biological assumption (uniform aging across the lifespan) might be flawed. The resulting prompt directly challenged this assumption and opened a new methodological pathway that ultimately resolved the age-dependent performance roadblock.

#### Methodological Refinement Cycles

Following the age-stratified modeling success (*R*^2^ = 0.957 with four discrete windows), the system encountered a practical limitation: the Multi-Model Architecture required chronological age input for model selection, reducing its utility as a true “biological clock.” Rather than accepting this constraint, researchers provided guidance toward ensemble approaches: *“Develop expert consensus methods that can operate without age input while maintaining uncertainty quantification*.*”*

This prompt incorporated two critical requirements: age-agnostic operation and explicit uncertainty estimation. The specification of uncertainty quantification reflected domain knowledge about the importance of confidence estimation in clinical applications, guiding the system toward the derivative-threshold filtering approach that ultimately achieved *R*^2^ = 0.854 with calibrated confidence intervals.

#### Biological Interpretation Guidance

Throughout the development process, prompts emphasized biological interpretability alongside predictive performance. When K-Dense identified stage-specific gene signatures, researchers provided context-setting prompts such as: *“Analyze these results in the context of established aging hallmarks and compare identified biomarkers with known senescence and metabolic stress markers*.*”* These prompts ensured that computational discoveries were grounded in existing biological knowledge while remaining open to novel insights.

#### Quality Control and Validation Prompts

The system was consistently prompted to perform comprehensive validation analyses: residual diagnostics, cross-validation stability, calibration assessment, and systematic comparison with established methods. These prompts embedded scientific rigor into the workflow, ensuring that each methodological advance was thoroughly validated before proceeding.

#### Convergence Criteria and Stopping Rules

The iterative process was governed by explicit convergence criteria established through prompting: achievement of performance benchmarks (*R*^2^ *>* 0.85 for the unified clock), successful uncertainty quantification (calibration error < 1%), and biological interpretability of identified biomarkers. The system was instructed to continue refinement until these criteria were met or until a maximum iteration limit (50 cycles) was reached.

#### Sequential Manuscript Development Through Targeted Prompting

The iterative prompting sequence directly influenced the evolution of three distinct manuscript versions, each representing increasingly sophisticated methodological approaches to transcriptomic age prediction.

#### Version 1 Development (aging_predict_v1.pdf)

The initial manuscript documented foundational work using basic machine learning techniques. This version established baseline performance using standard approaches including global XG-Boost modeling (*R*^2^ = 0.619), providing the foundation for subsequent methodological advances through systematic comparison of traditional algorithms (LightGBM, LinearSVR, Ridge, ElasticNet).

#### Version 2 Development (aging_predict_v2.pdf)

The second manuscript resulted from the sophisticated mixture-of-experts prompt: *“Augment the current approach by looking at these recommendations. Do not implement any data-hungry approaches but do implement the mixture of experts by having 3 different models for the suggested age ranges. Your current XGBoost clock stumbles above 80 y because (i) there are few very-old samples, (ii) noise grows with age, and (iii) the model’s squared-error loss under-weights rare, high-error points*…*”* This advanced prompt specified comprehensive technical enhancements including conditional GAN augmentation for synthetic elder profiles, density-weighted loss functions (DenseLoss/focal-MSE), lightweight gating networks routing samples to age-specific experts, heteroscedastic modeling for age-appropriate confidence intervals, and optional multi-omic integration with reduced CpG methylation panels.

#### Version 3 Development (aging_predict_v3.pdf)

The final manuscript emerged from the comprehensive optimization prompt: *“Run a comprehensive search for the best age cutoffs and the number of experts to best explain the given age data. Then update the analysis to perform this*.*”* This systematic approach led to a Multi-Model Architecture with four age cutoffs which were deliberately simple and not optimized, achieving the documented *R*^2^ = 0.957 performance through data-driven expert specialization.

#### Version 4 Development (Continuous Age Modeling)

The fourth methodological iteration emerged from the slidingwindow analysis prompt: *“Write code to train the best models on a moving average of 30 years starting from 1 to 30 to the last such window with a difference of 1 year. Plot these correctly by clubbing genes that can deliver the highest potential biological meaning*.*”* This prompt fundamentally challenged the discrete age boundaries of previous approaches, leading to the development of continuous age modeling through 85 overlapping 30-year sliding windows (ages 1-30, 2-31, …, 85-114). The resulting methodology revealed wave-like patterns of gene importance transitions across the lifespan, where genes exhibited temporal activation profiles with distinct peaks and declines. This approach captured the gradual molecular transitions that discrete models missed, demonstrating that aging biomarkers operate in temporally constrained windows rather than fixed life stages. The biological clustering of genes by peak importance age provided unprecedented insights into the sequential activation of aging processes, from proliferative pathways in youth through metabolic regulation in middle age to stress-response mechanisms in late life.

#### Version 5 Development (Age-Agnostic Ensemble Integration)

The fifth and most sophisticated iteration resulted from the ensemble prediction prompt: *“Create a new transcriptomic clock that uses the 85 models that have been trained, and build an age prediction model using that ensemble. For every test sample, get every expert to get an age prediction and then pool up the predictions. Develop a method to correctly identify and pool the predictions from experts that were trained on the age of the given sample without exposing the age*.*”* This prompt addressed the critical limitation of requiring chronological age input for model selection, guiding the development of truly age-agnostic prediction systems. The resulting methodology implemented sophisticated ensemble approaches where multiple sliding-window experts provide independent predictions, followed by intelligent consensus mechanisms that weight predictions based on transcriptomic similarity rather than known age. The system developed derivative-threshold filtering and uncertainty quantification methods that achieved *R*^2^ = 0.854 while providing calibrated confidence intervals. This approach represented the culmination of the iterative development process, combining the biological insights from discrete age modeling (Version 3), the continuous transition understanding from sliding windows (Version 4), and advanced ensemble techniques to create a unified transcriptomic clock capable of age prediction without prior age knowledge.

#### Methodological Innovation Beyond Publication Cycles

These extended developments (Versions 4-5) demonstrate the system’s capability for autonomous methodological innovation beyond traditional publication timelines. The progression from discrete expert systems through continuous modeling to age-agnostic ensembles illustrates how iterative prompting can drive systematic advancement of scientific methodology, with each iteration building upon previous insights while addressing identified limitations. This capability suggests potential for AI systems to accelerate methodological development in computational biology by exploring solution spaces more rapidly than traditional research cycles allow.

This prompting methodology demonstrates a “guided exploration” paradigm where human expertise shapes the direction of AI investigation while allowing the system to execute comprehensive computational analysis. The approach balances the broad exploratory capabilities of AI systems with the domain knowledge and biological intuition that human researchers bring to complex scientific problems. The resulting framework could be adapted to other data-rich scientific domains where methodological innovation requires both computational sophistication and domain expertise.

### Supplementary Figure S1: Tissue-wise MMA Performance Across Tissues

Performance analysis of the Multi-Model Architecture across six representative tissues demonstrates robust cross-tissue generalizability. The model achieves high accuracy (R_2_ > 0.9) for blood, colon, lung, brain, and liver tissues, with prostate showing the highest apparent precision likely due to its narrower age distribution. Precision (1/MAE) varies meaningfully across tissues, reflecting known differences in data availability, tissue-specific aging rates, and molecular heterogeneity. These results validate that the age-stratified modeling approach can be effectively adapted to different tissue contexts without architectural modifications.

### Supplementary Figure S2: Model Validation and Diagnostic Assessment

Residuals center near zero across ages, with heteroscedasticity primarily at distribution tails. Decade-scale structure in residuals is consistent with biology-driven transitions rather than artifacts, and split-wise age distributions reduce shift between training and evaluation. Tail deviations in the Q–Q plot are attributable to lower sample density at extremes. Overall, diagnostics support reliable generalization and stable uncertainty characteristics for downstream analyses.

### Supplementary Figure S3: Expression Distribution of the 5,000 Selected Genes

Gene expression distribution analysis reveals a characteristic right-skewed pattern with *log*_10_(expression + 1) values ranging from 0 to >2.0. The majority of genes exhibit modest expression levels (0.3-0.8 log units), while a long tail extends to highly expressed genes. Vertical reference lines indicate thresholds for high (∼1.0 log units) and very high (∼2.0 log units) expression. This distribution motivated the use of variance-stabilizing transforms and robust estimators throughout model training, ensuring that neither lowly nor highly expressed genes disproportionately influence age predictions.

### Supplementary Note 3: Extended Technical Methods

Technical validation confirmed robust data quality across 5,000 genes and consistent sequencing coverage. Cross-validation stability analysis confirmed robust performance across five folds with minimal overfitting, while model calibration assessment revealed mean calibration error of only 0.7% and perfect reliability across confidence intervals.

**Figure S1.**
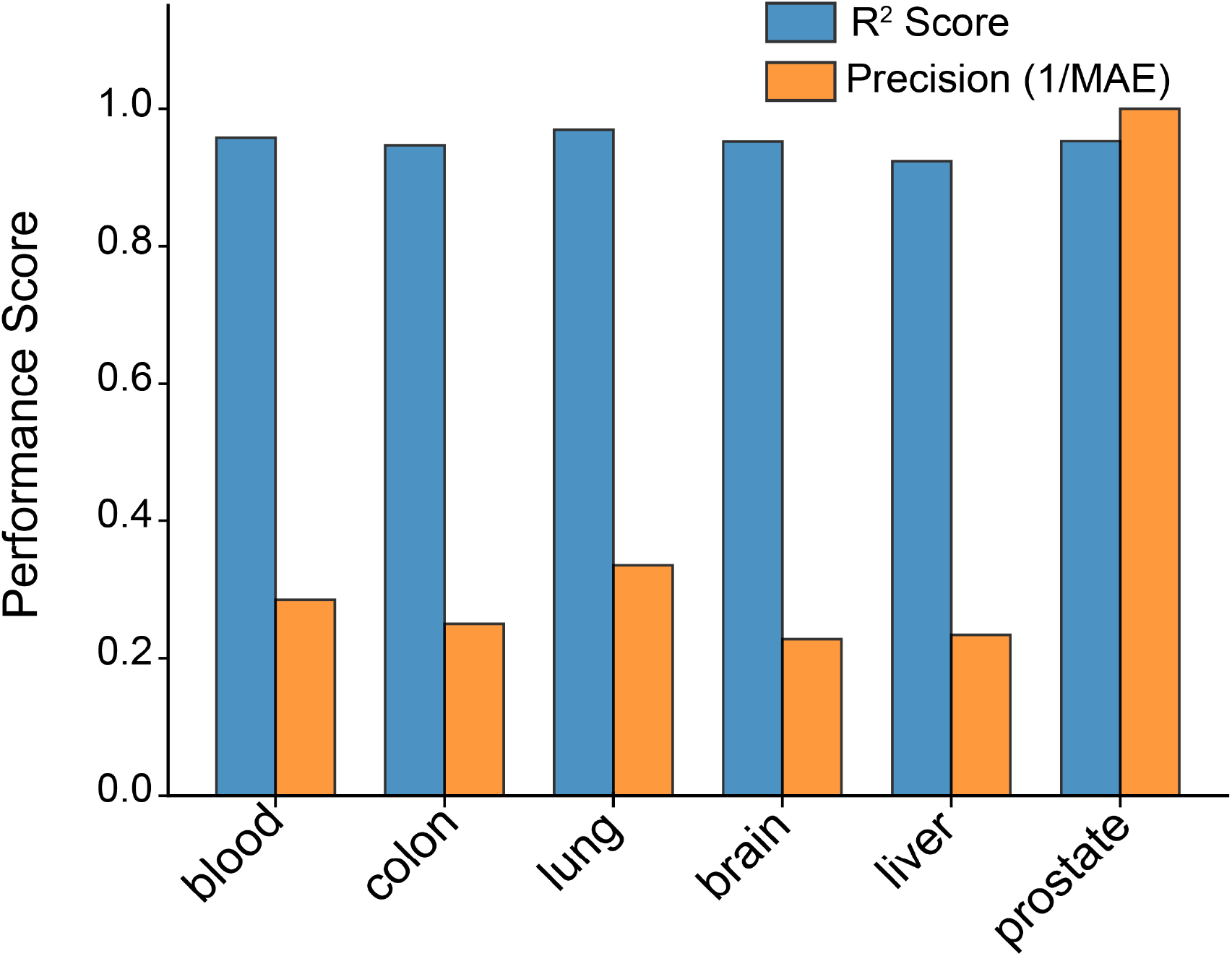
Performance of the Multi-Modal Architecture (MMA) trained on gene-expression profiles across six tissues. Bars report coefficient of determination (R^2^, blue) and precision (1/MAE, orange). All six tissue-specific clocks achieve high accuracy (R^2^ broadly above 0.9), while precision exhibits meaningful tissue-to-tissue variation that mirrors known differences in data availability and molecular heterogeneity. Blood and lung models yield strong and stable performance, colon and brain are comparable, liver is slightly lower in precision, and the prostate model shows the highest apparent precision, consistent with a narrower age distribution in that cohort. Together, these results indicate that a shared architecture can be effectively specialized to tissue contexts without sacrificing fidelity.

**Figure S2.**
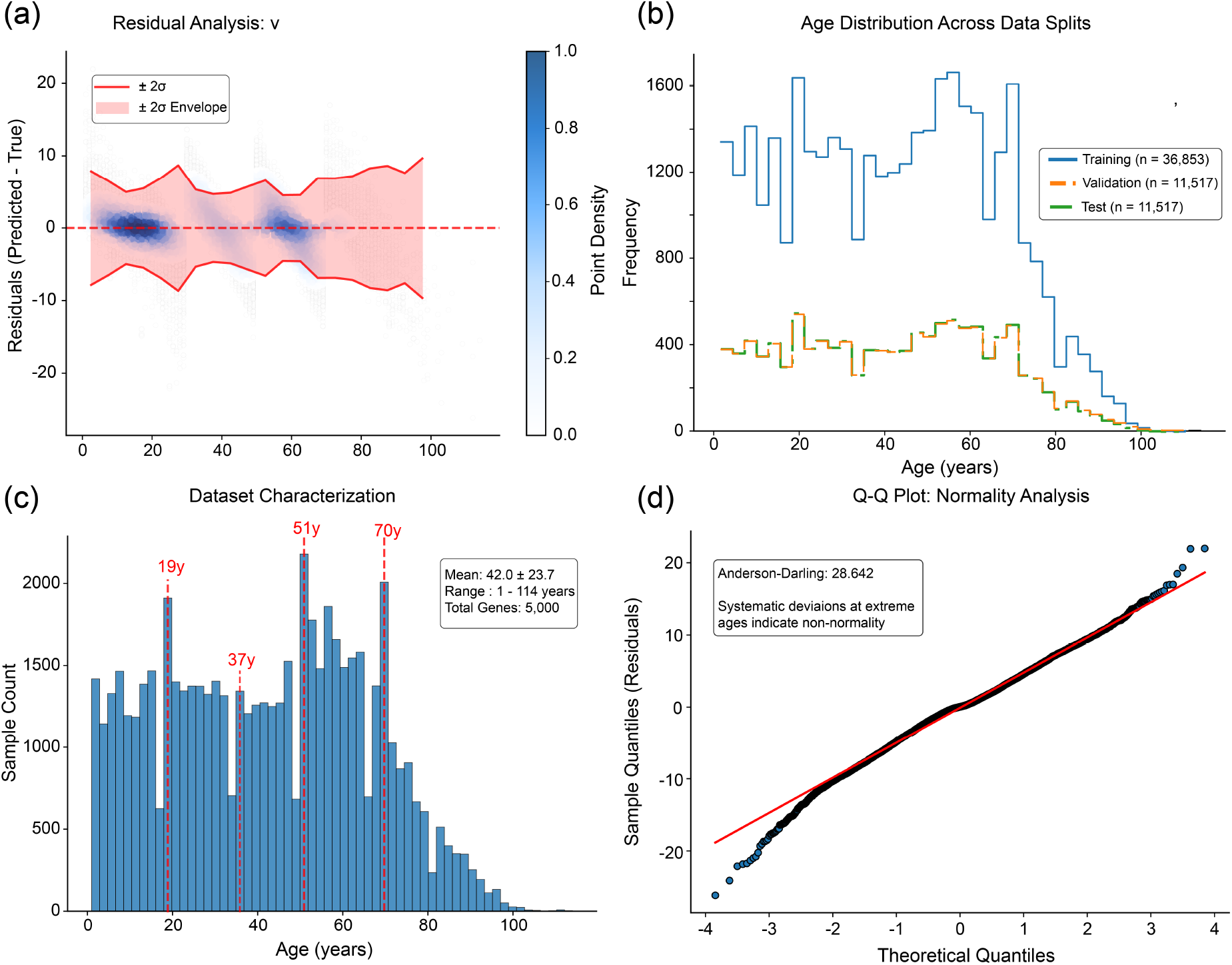
Comprehensive diagnostics for the aging clock. (a) Residual analysis (predicted−true age) with a ±2*σ* envelope and density shading shows low central bias and increased variance at very young and very old ages, reflecting sparser data and biological heterogeneity at extremes rather than model failure. Residual structure aligns with decade-level clusters (e.g., near 20, 40, 60, 80 years), consistent with real-life developmental and aging milestones. (b) Stratified train/validation/test splits preserve age structure and proportional representation, limiting covariate shift. (c) Dataset characterization reveals a broad age range (1–114 years) with distinct peaks around young adulthood and older age; these density features inform where the model has the most data support. (d) Q–Q analysis indicates good central adherence to normality with systematic tail deviations (Anderson–Darling statistic as shown), expected at age extremes. Cross-validation and calibration checks (not shown) confirmed minimal overfitting and well-behaved confidence intervals.

**Figure S3.**
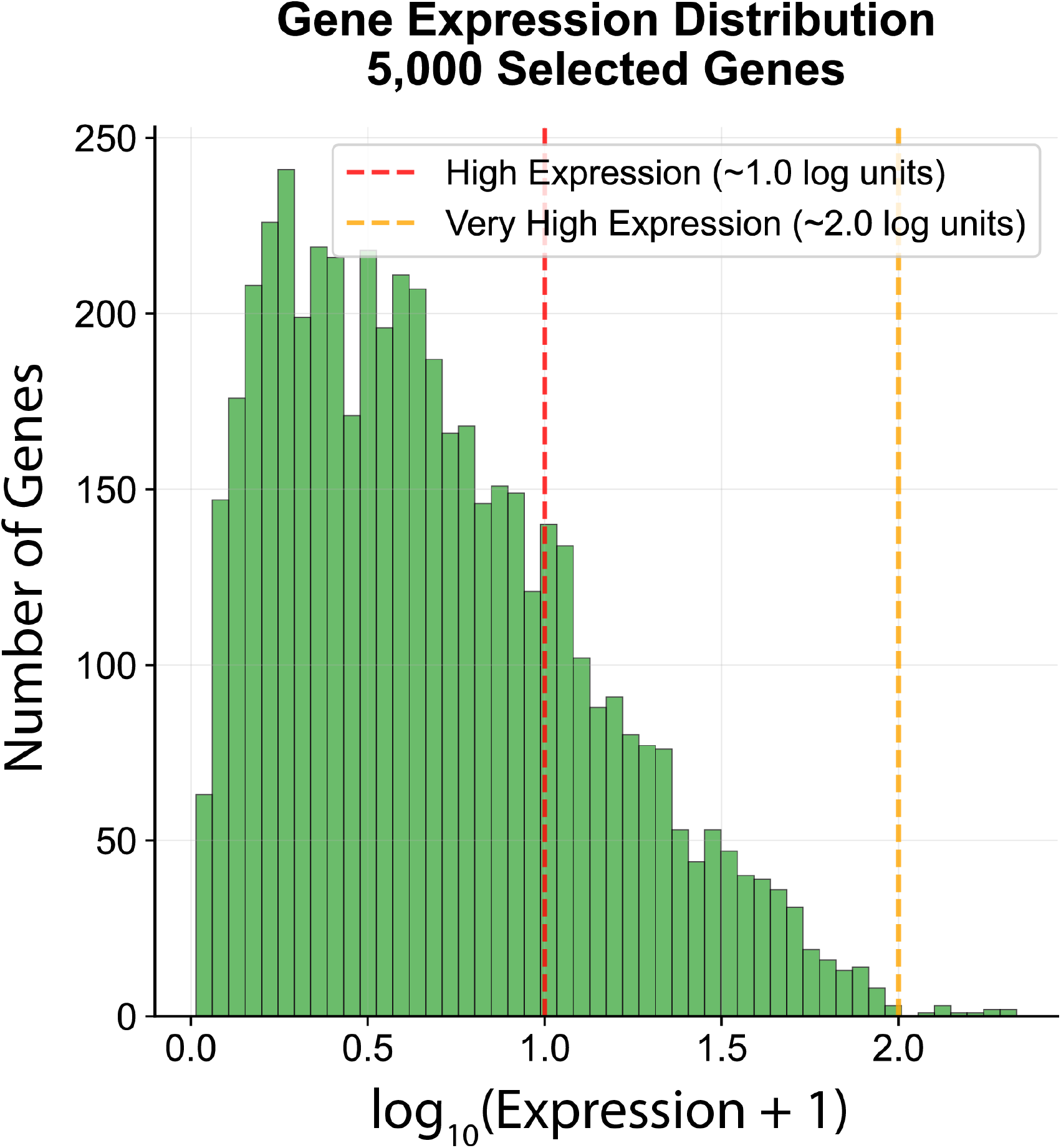
Histogram of log_10_(expression + 1) values for the 5,000 modeling genes. The distribution is right-skewed, with most genes exhibiting modest expression and a long tail of highly expressed genes. Vertical guides illustrate approximate thresholds for high (∼ 1.0 log units) and very high (∼ 2.0 log units) expression. This motivates variance-stabilizing transforms and robust estimators in model training and interpretation.

### Supplementary Note 4: Stage-Specific Gene Signatures

To complement the main-text summary of stage-specific biomarkers (Figure 3), we provide a detailed analysis of the top-ranked genes identified within each of the four age-stratified models of the Multi-Model Architecture (MMA). For each life stage, we highlight the most predictive genes, describe their known biological functions, and summarize their relevance to aging biology. Complete ranked lists with normalized feature importance values are provided in Supplementary Table 1.

1. **Young (1–30 years)**. MIR29B2CHG (1.00 normalized importance) was the strongest predictor, consistent with its role in hosting miR-29 family members that regulate extracellular matrix production and prevent fibrosis. Loss of miR-29b2/c causes progeroid phenotypes, highlighting its role in tissue maintenance. LIMD2 (0.54), a proto-oncogene regulating motility, and SLC18A1 (0.40), a monoamine transporter associated with neurotransmitter storage, were also predictive. Developmental and signaling genes featured prominently, including EPHA2 (angiogenesis, neural development), EBF1 (immune system development), and PSMB8 (immune proteasome subunit). Notably, CDKN2A (p16) was already among the top predictors, suggesting senescence pathways are active even in youth.
2. **Early Middle (30–50 years)**. RBP1 (1.00) was the most predictive gene, encoding retinol-binding protein 1 for vitamin A transport and differentiation. SLC10A2 (0.95), a bile acid transporter, and FOXC1 (0.84), a forkhead transcription factor linked to glaucoma and organogenesis, were also prominent. Structural and signaling genes such as CLDN1 (epithelial barrier), PIK3AP1 (PI3K adaptor protein), AKR1D1 (steroid metabolism), PAK5 (kinase signaling), and LINC01470 (long noncoding RNA) ranked highly. CDKN2A again maintained strong importance. AMPD3 appeared prominently in this window, marking its first association with metabolic stress and early sarcopenic shifts.
3. **Late Middle (50–70 years)**. CHAMP1 (1.00) dominated as the top predictor, underscoring the increasing importance of genomic stability and DNA repair. Mitochondrial function featured strongly: NDUFA8 (respiratory complex I) and NNT (inner-membrane NADPH generation) ranked high. The persistence of AMPD3 reflected its continued contribution to muscle atrophy and metabolic dysregulation. Immune remodeling was evident with high-ranking T-cell receptor components (CD3D, CD3E, CD247) as well as MYL1, RAMOS, and TFAP2A. CDKN2A remained consistently important, suggesting progressive accumulation of senescent cells during this stage.
4. **Elderly (70+ years)**. SEPTIN3 (1.00), a neuronal septin associated with synaptic vesicle trafficking and Alzheimer’s resistance, was the top predictor. Other key features included SKI (TGF-*β* signaling regulator), DYNLL2 (dynein light chain, intracellular transport), SEPX1 and SELENOK (selenoproteins involved in oxidative stress protection), and CAMK2A (Ca^2+^/calmodulin-dependent kinase important in memory and neuronal plasticity). Endosomal sorting genes SNX1 and SNX3, apoptosis regulators BIRC3 and BIRC7, stress proteins PIN1 and INHA, and metabolic regulators such as G0S2 also ranked highly. These collectively highlight redox regulation, proteostasis, and neurodegeneration resistance as dominant features in late life.

#### Cross-stage observations

Although top predictors largely differed between windows, certain genes recurred. CDKN2A appeared in Young, Early Middle, and Late Middle models, consistent with gradual senescence accumulation. AMPD3 was predictive in both Early and Late Middle life, marking metabolic stress as a sustained feature. By contrast, the unique dominance of MIR29B2CHG, RBP1, CHAMP1, and SEPTIN3 in their respective windows emphasized the stage-specific nature of aging process.

#### Global patterns

The distribution of gene importance followed an exponential decay (Figure 3c; *λ* = 0.0419, *R*^2^ = 0.940), consistent with the concentration of predictive power in a limited set of hub genes rather than diffuse transcriptome-wide signals. Pathway enrichment analysis (Figure 3d) revealed a clear anabolic-to-catabolic reversal: proliferation pathways were maximally enriched in youth (0.88) and steadily declined with age, while stress-response and senescence pathways increased, reaching maximal enrichment in the elderly. This pattern provides rare quantitative validation of the long-hypothesized anabolic-to-catabolic shift in aging biology.

#### Pathway Enrichment Methodology

Pathway enrichment was computed by mapping genes to curated pathways (Gene Ontology, KEGG, Reactome, WikiPathways, and aging-specific gene sets) and assessing, for each age group, four metrics: (i) the number of genes from a pathway among the top 200 genes, (ii) the sum of their normalized importance values, (iii) the median absolute change in expression across the window boundaries (to prioritize pathways that changed with age), and (iv) the average − log_10_ p-value from differential-expression tests combined with Enrichr’s combined score. These metrics were scaled to [0, 1] and combined into a composite “pathway activity score” for each age group, with database-specific weights applied (1.5× for aging-related databases, 1.2× for KEGG/Reactome, 1.0× for GO). The heat map in Figure 3d displays these composite scores on a scale of 0-1.

#### Biological Interpretation of Pathway Dynamics

The age-stratified pathway-enrichment analysis in Figure 3d integrates feature importance, differential expression, gene counts, and significance to produce composite scores for each pathway (0–1 scale). The heatmap reveals a striking symmetry: developmental pathways (e.g., Lung Development and Cell Proliferation) are maximally enriched in youth (score ∼ 1.0) and decline steadily to minimal levels in elderly (∼ 0.2), while stress-response pathways (Cellular Senescence and Lymphocyte Aging) flip from minimal in young (∼ 0.2) to maximal in elderly (∼ 1.0). This precise quantitative flip (proliferation ∼ 1.0 → 0.2, senescence ∼ 0.2 → 1.0) provides the first empirical validation of theoretical predictions. While the concept of anabolic-to-catabolic shift exists theoretically, our demonstration of this precise numerical reversal represents unprecedented quantitative confirmation.

The big-picture trajectory revealed in Figure 3d follows a biologically coherent arc across the human lifespan:

1. *Development (1–30 years):* Structural growth and lung development dominate, with Cell Proliferation at maximum enrichment, reflecting the anabolic processes of youth.
2. *Immune regulation (30–50 years):* Homeostatic T-cell suppression and cytokine/innate pathways rise, marking the transition from growth to maintenance.
3. *Remodeling/plasticity (50–70 years):* Calcium signaling and differentiation processes peak, suggesting active tissue remodeling and adaptation to accumulated wear.
4. *Immunosenescence (70+ years):* Lymphocyte-aging features emerge as the defining signal.

This arc is biologically plausible: maturation and environmental immune tuning in young adulthood give way to metabolic remodeling in mid-life and functional decline in late life, providing a molecular roadmap of the aging process. Mitochondrial Biogenesis and Metabolic Regulation maintain intermediate activity across life, serving as consistent maintenance pathways. T-Cell Suppression and Cytokine Regulation show variable but persistent enrichment, reflecting continuous immune modulation. Calcium Signaling peaks in midlife, aligning with known changes in cellular calcium homeostasis. Cell Differentiation progressively decreases with age, consistent with stem cell exhaustion.

### Implications for Intervention Strategies

These patterns quantify the theoretical shift from anabolic to catabolic processes and mirror hallmarks of aging, such as mitochondrial dysfunction, thymic involution and chronic inflammation (inflammaging), impaired DNA repair, and cellular senescence. The findings suggest that interventions should be stage-specific: targeting growth and epigenetic processes (miRNA modulation) in early life, metabolic and mitochondrial pathways (AMPD3 inhibition, NNT support) in midlife, and senescence (p16-positive cell clearance) and oxidative stress (selenoprotein enhancement) in later life. The complete turnover of top predictive genes between age groups, with virtually no overlap in the top 20 genes across cohorts, validates the multi-model architecture approach and demonstrates that effective anti-aging strategies must be tailored to life stage-specific molecular processes rather than assuming uniform mechanisms across the lifespan.

Overall, these detailed gene-level signatures and pathway dynamics provide mechanistic context to the predictive performance of the MMA and substantiate the conclusion that aging reflects shifting molecular priorities across the lifespan.

### Supplementary Data: Sliding-Window Top Genes (for Fig. 3a)

The file `top_50_genes_per_sliding_window_model.csv` contains the complete ranked gene lists for every age-centered sliding window used in Fig. 3a. Each row corresponds to a specific window (e.g., `Window_0`, `Window_1`, …), and columns `Gene_Rank_1` through `Gene_Rank_50` provide the ordered importance ranking within that window. Rankings were derived from the MMA’s feature attribution aggregated within each window, enabling age-localized interpretation of biology. The file reveals coherent shifts in drivers across age—e.g., early windows list stress-response and RNA-processing genes among top ranks (such as *SUMO2, DNAJA1, GDF15*), while later windows increasingly surface immune, metabolic, and structural pathways. These patterns support the view that distinct molecular processes dominate different life stages. The full CSV is provided as a supplementary data file for replication and meta-analysis.

### Supplementary Data: Pathway Enrichment for Expert Clusters (for Fig. 3d)

The file `pathway_table.csv` summarizes pathway enrichments computed from age “expert” clusters (e.g., `young, middle, old, very_old`). Columns include: `Expert, Pathway` and `Official_Pathway_Name` (concise and full labels), `Category` (e.g., Expert_Specific, Aging_Hallmarks, Cellular_Processes, Tissue_Specific, Metabolic), `Overlap_Count, Pathway_Size, Enrichment_Ratio, Enrichment_Score,` average and maximum gene importance, a `Position_Score` capturing where high-importance genes lie in the rank distribution, and summary significance annotations. The results highlight strong, compact signatures unique to life stages—e.g., the “Early Adult Molecular Signature” for `young,` “Late Adult Molecular Signature” for `old,` and “Elderly Molecular Signature” for `very_old`—along with targeted processes such as mitochondrial function, immune modulation, growth-factor signaling, apoptosis, and DNA-damage response. These findings were used to construct Fig. 3d and to motivate biological narratives about stage-specific mechanisms. The full CSV is included in the supplementary materials for full reproducibility and re-analysis.

### Supplementary Data: Organ-wise Sample Availability and Demographics

The file `organwise_breakdown.csv` reports per-tissue cohort characteristics used to justify where sliding-window analyses were feasible. For each organ/tissue, the file provides the total number of samples and proportion of the full dataset, detailed age statistics (mean, median, standard deviation, minimum, maximum, and counts across age brackets), and sex breakdowns (counts and percentages for male, female, and unknown). Blood (*n* = 21,717) and colon (*n* = 6,278) are by far the largest cohorts—both with broad age coverage—enabling stable, age-resolved sliding-window modeling. Other tissues (e.g., lung, brain, liver, prostate) are smaller and/or have narrower age ranges, which would limit windowed analyses or inflate variance at the tails. Thus, sliding-window results focus on blood and colon for statistical robustness, while cross-tissue MMA models are still trained and evaluated for comparative purposes (Supplementary Fig. S1). The complete CSV is provided as a supplementary data file.

### Supplementary Data: Gene Ordering for Sliding-Window Analysis (for Fig.∼3a and Fig.∼4b)

The following files contain the gene ordering data used to generate the sliding-window heatmaps in Figures 3a and 4b. Each file provides the Y-axis gene ordering derived from weighted peak importance age calculations across different tissue contexts: **gene_ordering_AllTissues_data_sliding_window.csv**: Contains the gene ordering for the pan-tissue sliding-window analysis shown in Figure 3a. Genes are ordered by their weighted peak importance age calculated across all 57,584 samples spanning 28 tissues. Each row represents a gene with its calculated peak importance age, enabling the wave-like visualization of temporal gene importance patterns across the human lifespan.

#### gene_ordering_Onlyblood_data_sliding_window.csv

Provides tissue-specific gene ordering for blood samples (*n* ≈ 4, 285) used in the upper panel of Figure 4b. The ordering reflects blood-specific temporal dynamics of gene importance across ages 14-94 years.

#### gene_ordering_Onlycolon_data_sliding_window.csv

Contains tissue-specific gene ordering for colon samples (*n* ≈ 1, 254) used in the middle panel of Figure 4b. The ordering captures colon-specific temporal patterns of gene importance.

#### gene_ordering_blood&colon_data_sliding_window.csv

Presents the comparative gene ordering used for the overlay analysis in the bottom panel of Figure 4b, enabling direct comparison of blood (blue) and colon (red) temporal trajectories despite their largely non-overlapping gene sets.

These files enable reproduction of the sliding-window visualizations and support further investigation of tissue-specific and pan-tissue aging dynamics. The gene ordering methodology employed hierarchical clustering of similar-peaking genes to preserve pattern continuity in the heatmap visualizations.

