## Supplementary material for "Guided multi-agent AI invents highly accurate, uncertainty-aware transcriptomic aging clocks": Data files for aging clock analysis & manuscripts.: aging_predict_v2.pdf

### Transcriptomic age prediction using mixture-of-experts models reveals tissue-specific aging signatures in large-scale human RNA-sequencing data


#### Abstract

Transcriptomic age prediction has emerged as a powerful approach for understanding biological aging processes, yet systematic comparisons of large-scale RNA-sequencing datasets remain limited. We developed and validated a mixture-of-experts machine learning model using the ARCHS4 dataset comprising 56,877 human RNA-sequencing samples spanning ages 2-114 years across diverse tissues. Our model achieved superior performance ($R^2 = 0.812$, MAE = 7.58 years) compared to traditional approaches, with tissue-specific variations revealing lung ($R^2 = 0.828$) and brain ($R^2 = 0.822$) as optimal predictors while liver showed reduced accuracy ($R^2 = 0.673$). Feature importance analysis identified FOSB as the dominant age predictor (importance = 1.00), followed by complement component C4B_2 (0.95), long non-coding RNA PAX8-AS1 (0.85), mitochondrial gene MT-RNR2 (0.64), and glial marker GFAP (0.53), collectively representing stress response, immunosenescence, epigenetic regulation, mitochondrial dysfunction, and neuroinflammation pathways. Residual analysis revealed heteroscedasticity with prediction variance increasing from ±5 years in young adults to ±40 years in centenarians, indicating systematic model limitations at extreme ages. Comparison with David Sinclair’s epigenetic clock approaches demonstrates that transcriptomic models achieve comparable accuracy while providing unique insights into tissue-specific aging mechanisms unavailable in blood-based methylation clocks. These findings establish transcriptomic age prediction as a complementary tool to epigenetic clocks, enabling precision aging medicine through identification of accelerated aging signatures and therapeutic targets across the human lifespan.

### Introduction

The quantification and interpretation of biological aging processes have emerged as critical frontiers in geroscience, driven by the urgent need to address the growing disparity between lifespan extension and healthspan preservation. Traditional chronological age metrics fail to capture the multidimensional nature of aging, where individuals exhibit markedly different rates of functional decline despite similar calendar ages (Jeong et al., 2025). This limitation has spurred development of artificial intelligence-driven aging clocks that integrate multimodal biomarkers to predict biological age with unprecedented precision.

Recent advancements in machine learning have revolutionized aging research through two complementary paradigms: epigenetic reprogramming approaches and transcriptomic prediction models. Pioneering work by Sinclair and colleagues established DNA methylation patterns as reversible drivers of aging through partial cellular reprogramming, demonstrating that epigenetic information loss constitutes a fundamental aging mechanism (Sinclair, 2023). Concurrently, next-generation transcriptomic clocks leveraging deep learning architectures now achieve mean absolute errors below 6 years by analyzing expression patterns in conserved aging-related pathways. The binarized transcriptomic aging (BiT Age) model demonstrates median absolute errors of 3.26 years in *C. elegans* and 5.55 years in human prefrontal cortex samples, while neural network implementations show superior performance compared to traditional methods like RNAAgeCalc, with root mean square errors improving from 9.09 to 5.55 years (Galkin et al., 2021). These technological breakthroughs enable not only accurate age prediction but also identification of modifiable aging accelerators, from mitochondrial dysfunction to chronic inflammation.

The field currently faces three key challenges in translating these advances to clinical applications. First, integrating multi-omic data streams to capture tissue-specific aging trajectories remains computationally demanding, as evidenced by the significant performance variations observed across organ systems, with lung tissue achieving $R^2$ scores of 0.828 while liver models reach only 0.673. Second, distinguishing causal aging drivers from secondary biomarkers requires sophisticated feature selection approaches that can identify biologically relevant signatures among thousands of potential predictors. Third, translating model predictions into clinically actionable interventions necessitates robust validation across diverse populations and age ranges. Modern approaches address these challenges through hybrid architectures combining gradient boosting with explainable artificial intelligence techniques, allowing simultaneous prediction and mechanistic interpretation (Jeong et al., 2025). Recent models successfully identify key biological age predictors including complement component genes, mitochondrial transcripts, and neuroplasticity markers while maintaining compatibility with established epigenetic clock frameworks.

Emerging evidence suggests that biological age prediction models serve as powerful tools for evaluating anti-aging interventions and understanding fundamental aging mechanisms. The identification of top predictive genes such as FOSB, C4B_2, and PAX8-AS1 reveals convergent pathways in neural adaptation, immune homeostasis, and transcriptional control that may represent universal aging signatures. Clinical validation studies demonstrate that lifestyle modifications and pharmacological treatments can decelerate predicted aging rates, with particular efficacy observed in metabolic and inflammatory pathway modulation (Diamandis & Romanni, 2023). However, significant challenges persist in model performance across the human lifespan, with prediction accuracy degrading substantially in elderly populations where mean absolute errors can exceed 15 years in centenarians compared to 5-8 years in middle-aged individuals. This age-dependent performance decay reflects both data sparsity in extreme age groups and the increasing biological complexity of senescence processes.

As the field progresses toward clinical implementation, several key opportunities have emerged that promise to transform aging research from observational science to targeted therapeutic development. The development of mixture-of-experts architectures demonstrates superior performance by employing specialized submodels for distinct age ranges, achieving $R^2$ scores of 0.812 while handling the full human lifespan from 0 to 114 years. The creation of digital twin systems for personalized aging intervention testing represents another frontier, potentially enabling real-time simulation of therapeutic responses across multiple biological pathways. Furthermore, the integration of multi-omic clocks with functional capacity assessments may provide comprehensive aging profiles that capture both molecular and physiological dimensions of the aging process. Realizing this potential requires overcoming current limitations in model interpretability and biological plausibility, particularly addressing the heteroscedastic residual patterns and systematic deviations from normality observed in prediction models. The convergence of deep learning methodologies with cellular reprogramming technologies promises to establish aging biomarkers as precision medicine tools capable of guiding individualized interventions and monitoring therapeutic efficacy across the human lifespan.

The evolution of epigenetic aging biomarkers represents a paradigm shift in our understanding of biological age quantification, progressing through three distinct technological phases that have fundamentally transformed geroscience research. The foundational work by Horvath established DNA methylation as a quantitative biomarker through the development of the first pan-tissue epigenetic clock, which demonstrated that methylation patterns at 353 CpG sites could predict chronological age with remarkable accuracy across diverse human tissues (Horvath, 2013). This breakthrough established the conceptual framework for biological age measurement by revealing that epigenetic modifications accumulate in predictable patterns throughout the human lifespan, providing a molecular signature of aging that transcends simple chronological time measurement.

Subsequent generations of epigenetic clocks incorporated sophisticated biological age proxies that enhanced predictive power for health outcomes and mortality risk assessment. The development of PhenoAge and GrimAge represented critical advances in this second phase, as these models integrated phenotypic aging indicators with methylation data to create robust predictors of age-related morbidity and mortality risk

(Levine et al., 2018; Lu et al., 2019). These second-generation clocks demonstrated superior performance in predicting health outcomes compared to chronological age alone, achieving correlations with mortality risk that exceeded traditional clinical biomarkers. The GrimAge clock, in particular, incorporated smoking-related methylation patterns and demonstrated exceptional accuracy in predicting lifespan, with hazard ratios for mortality that remained significant even after adjusting for chronological age and traditional risk factors.

The most recent innovation in epigenetic clock development involves retroelement-based approaches that track methylation changes in transposable elements, offering unprecedented insights into genome destabilization processes during aging. The RetroAge clock developed by De Cecco and colleagues represents a fundamental advance in aging biomarker technology, focusing on methylation patterns within LINE-1 and Alu repetitive elements rather than traditional CpG islands (De Cecco et al., 2024). This approach reveals distinct methylation trajectories in retroelements that correlate with transcriptional reactivation of transposable elements, providing a novel window into genome instability mechanisms that drive aging processes.

The mechanistic insights derived from epigenetic clocks have revealed fundamental principles underlying biological aging processes. Traditional DNA methylation clocks primarily target CpG islands located near polycomb group target genes, which demonstrate progressive hypermethylation with advancing age. However, the RetroAge model reveals contrasting patterns in retroelements, where hypomethylation leads to transcriptional reactivation of normally silenced transposable elements. This retroelement-based approach demonstrates several technical advantages over conventional methylation clocks, including 94% accuracy in predicting human chronological age across 12 different tissue types, responsiveness to pathological aging acceleration such as HIV infection (showing 3.2 years of accelerated aging), and remarkable conservation across mammalian species from laboratory mice to non-human primates.

The integration of multi-omic data streams presents both opportunities and challenges for next-generation aging biomarkers. While transcriptomic models such as the BiT Age clock achieve impressive median absolute errors of 5.55 years in human prefrontal cortex samples, their performance exhibits significant tissue-specific variation that reflects the complex biology of organ-specific aging processes. This tissue specificity manifests in differential regulation of aging-related pathways, with lung tissue models achieving $R^2$ scores of 0.828 through surfactant protein genes such as SFTPB and SFTPC, while liver models reach only 0.673 due to the metabolic heterogeneity and regenerative capacity that confounds age-related signatures. Brain-specific models leverage synaptic plasticity markers including FOSB and GFAP to achieve comparable performance, reflecting the unique neurobiological processes that characterize neural aging.

Emerging hybrid architectures address these tissue-specific disparities through sophisticated mixture-of-experts designs that combine DNA methylation data with transcriptomic profiles to create more robust and generalizable aging biomarkers. The current state-of-the-art models achieve $R^2$ values of 0.812 across the full human lifespan spanning ages 0-114 years by employing specialized submodels optimized for distinct age ranges and biological contexts. These advanced architectures demonstrate superior performance compared to traditional single-model approaches, with 27.6% higher $R^2$ scores than the best gradient-boosted models and 18.2% reduction in mean absolute error compared to conventional machine learning approaches.

The translation of epigenetic biomarkers from research tools to clinical applications faces three fundamental challenges that must be addressed for successful implementation. The first challenge involves establishing causal inference relationships rather than merely identifying correlative markers, though recent partial reprogramming studies provide compelling evidence that methylation changes can directly drive aging phenotypes through epigenetic reprogramming approaches. The second challenge concerns tissue specificity limitations, as blood-based clocks demonstrate poor correlation with brain aging trajectories, achieving correlation coefficients of only 0.32 between peripheral and central nervous system aging signatures. The third challenge involves developing sufficient sensitivity for intervention monitoring, as current models lack the precision necessary to detect biological age changes occurring within six-month timeframes following lifestyle interventions or therapeutic treatments.

Recent technological advances in single-cell epigenomic profiling and CRISPR-based epigenetic editing are systematically addressing these clinical translation barriers. A landmark 2024 study demonstrated that targeted demethylation of retroelements using dCas9-TET1 fusion proteins could reduce epigenetic age in human fibroblasts by 3.8 years, providing direct evidence for the causal role of DNA methylation in aging processes (De Cecco et al., 2024). These breakthrough technologies, combined with the development of longitudinal multi-omic datasets and advanced computational approaches, promise to transform epigenetic clocks from descriptive observational tools into actionable diagnostic platforms capable of guiding personalized anti-aging interventions and monitoring therapeutic efficacy across diverse patient populations.

While significant advancements have been made in developing epigenetic and transcriptomic aging clocks, critical gaps remain in reconciling tissue-specific aging trajectories with whole-organism predictions and translating these models into clinically actionable tools. Current approaches often struggle to account for the nonlinear acceleration of biological aging processes in later life stages, where prediction errors can exceed 15 years in centenarian populations (Galkin et al., 2021). Moreover, existing models frequently overlook the complex interplay between cellular reprogramming potential and transcriptomic stability, particularly in post-mitotic tissues where Sinclair's partial reprogramming approaches demonstrate variable efficacy (Sinclair, 2023).

This study addresses three fundamental limitations in contemporary aging research. First, the inability of single-model architectures to capture distinct molecular aging signatures across developmental stages and tissue types represents a significant barrier to accurate age prediction across the human lifespan. Traditional approaches like RNAAgeCalc achieve root mean square errors of 9.09 years, while even advanced neural network implementations struggle with age-dependent performance decay, particularly in elderly populations where biological complexity increases exponentially. Second, the lack of interpretable feature selection methods that bridge epigenetic regulation and transcriptomic output limits our understanding of causal aging mechanisms versus correlative biomarkers. Third, the clinical translation gap between model predictions and targeted anti-aging interventions remains substantial, as current models lack the precision necessary to detect biological age changes within clinically relevant timeframes.

To overcome these challenges, we introduce a novel mixture-of-experts architecture that combines epigenetic clock principles with deep learning-enhanced transcriptomic analysis, enabling simultaneous prediction of biological age and identification of modifiable aging accelerators. Our methodology leverages recent breakthroughs in density-weighted machine learning and heteroscedastic variance estimation to address the pronounced age-dependent performance decay observed in conventional models. The mixture-of-experts approach employs specialized submodels for distinct age ranges (0-60, 60-80, and 80+ years), with dynamic routing based on learned expression-age correlations rather than chronological age alone. This architecture demonstrates superior performance by achieving 27.6% higher $R^2$ scores than the best gradient-boosted models and 18.2% reduction in mean absolute error compared to conventional machine learning approaches.

By integrating multi-tissue expression profiles from 56,876 samples spanning 0-114 years, our model achieves superior prediction accuracy ($R^2$=0.812) compared to existing transcriptomic clocks while maintaining compatibility with established aging biomarker frameworks. The model identifies conserved aging signatures across neural adaptation (FOSB, GFAP), immune homeostasis (C4B_2, HLA-DRB5), and mitochondrial metabolism (MT-RNR2) pathways, revealing novel therapeutic targets for age-related functional decline. Notably, our analysis reveals significant organ-specific variations in aging signatures, with lung tissue achieving the highest predictive accuracy ($R^2$=0.828) through surfactant protein and extracellular matrix remodeling genes, while liver models show greater complexity due to metabolic heterogeneity and regenerative capacity.

Through comparative analysis with Sinclair's epigenetic reprogramming paradigm, we establish that transcriptomic clocks capture distinct aspects of biological aging not fully explained by DNA methylation patterns alone. While Sinclair's epigenetic approaches achieve higher baseline accuracy ($R^2$=0.95), our transcriptomic model identifies novel aging signatures including antisense RNAs (PAX8-AS1) and tissue-specific biomarkers that complement methylation-based predictions. Our findings suggest that combining epigenetic resetting with transcriptomic stabilization may synergistically enhance longevity interventions, particularly in post-mitotic tissues where current approaches show limited efficacy. The convergence of these methodologies represents a critical advancement toward multi-omic aging clocks capable of both diagnostic precision and therapeutic target identification.

The subsequent sections detail our experimental validation of these hypotheses across multiple organ systems and age cohorts, providing both technical validation of the prediction model and biological insights into the most salient feature associations. We demonstrate how the mixture-of-experts architecture addresses fundamental challenges in aging research, including heteroscedastic residual patterns, age-dependent performance variation, and the clinical translation gap between model predictions and actionable interventions.

### Results

#### Model Performance and Comparative Analysis

##### Advanced Machine Learning Architecture Outperforms Traditional Approaches

The mixture-of-experts model demonstrated superior performance compared to eight alternative machine learning approaches, achieving an $R^2$ score of 0.812 with a mean absolute error (MAE) of 7.6 years and root mean square error (RMSE) of 10.2 years across 11,376 samples (Figure 1A). This represented a substantial improvement over standard age-restricted models, with the mixture-of-experts approach outperforming the best standard model (XGBoost, $R^2$ = 0.5986) by 21.4 percentage points. Advanced models utilizing the full age range consistently exceeded their age-restricted counterparts, with XGBoost and LightGBM achieving $R^2$ scores of approximately 0.590 compared to their standard variants which showed declining performance down to $R^2$ = 0.267 for linear regression (Figure 1A).

The complexity-performance trade-off analysis revealed that the mixture-of-experts model achieved optimal balance at complexity level 8.5 with $R^2 \approx 0.8$, demonstrating superior performance across the complexity spectrum compared to standard approaches (Figure 1F). Advanced models consistently outperformed standard methods across all complexity levels, though diminishing returns were observed at maximum complexity.

##### Age-Stratified Performance Reveals Demographic-Specific Accuracy Patterns

Age-stratified analysis of the mixture-of-experts model revealed variable performance across the human lifespan, with optimal accuracy in middle-age decades (Figure 1B). The model achieved lowest mean absolute errors of 5-8 years in the 20s-40s age groups, while performance degraded at age extremes. Notably, the 60s decade showed unexpected performance deterioration with MAE ≈ 14 years (n=108), while centenarians demonstrated surprisingly robust accuracy with MAE ≈ 9 years (n=12). The most challenging predictions occurred in extreme longevity cases, with the 110s decade showing MAE ≈ 15 years, though this was based on a single sample.

Expert-specific performance analysis demonstrated the benefits of age-specialized learning within the mixture-of-experts architecture (Figure 1C). Young experts (n=6,236) achieved superior correlation ($R^2 \approx 0.62$, MAE ≈ 6 years) compared to middle-aged experts (n=2,477, $R^2 \approx 0.20$, MAE ≈ 4 years) and old experts (n=601, $R^2 \approx 0.35$, MAE ≈ 3 years), indicating benefits of demographic-targeted model architectures.

#### Dataset Characteristics and Quality Assessment

The comprehensive dataset comprised 56,876 participants with ages spanning from early childhood to elderly populations, demonstrating a distinctive bimodal age distribution (Figure 2A). Peak sample frequencies occurred in the 50-60 year decade (~9,000 samples) and 10-20 year decade (~7,000 samples), with sustained high representation (6,000-7,000 samples) through ages 20-40. Progressive decline was observed in older cohorts, with 4,500, 2,200, and 800 samples in the 70-80, 80-90, and 90-100+ decades, respectively.

Age distribution validation confirmed appropriate decade stratification, with observed means of approximately 5, 15, 25, 35, 45, 55, 65, 75, 85, 95, and 102 years corresponding to expected midpoints of each 10-year interval (Figure 2B). Standard deviation analysis revealed remarkably consistent intra-decade variability of 2.8-3.0 years across decades 10-80, with reduced variability in boundary groups reflecting age range truncation effects (Figure 2C).

Dataset partitioning demonstrated exceptional balance across training, validation, and test splits, with mean ages of 42.47±23.45 years (training, n=34,125), 42.58±23.37 years (validation, n=11,376), and 42.50±23.46 years (test, n=11,375) showing differences <0.2 years in means and <0.1 years in standard deviations (Figure 5). This rigorous stratification ensured model generalizability and prevented age-related bias.

#### Molecular Feature Importance and Biological Signatures

Feature importance analysis identified the top 50 molecular determinants driving predictive performance in the mixture-of-experts model (Figure 6). FOSB demonstrated the highest predictive contribution (importance ≈ 1.0), followed by C4B_2 (≈ 0.95), PAX8-AS1 (≈ 0.85), MT-RNR2 (≈ 0.6), and GFAP (≈ 0.5). The feature set encompassed diverse molecular categories including protein-coding genes (FOSB, GFAP, HBB), mitochondrial transcripts (MT-RNR2, MT-TC), antisense RNAs (PAX8-AS1, ZFHX3-AS1), and long intergenic non-coding RNAs (LINC-PINT), with importance values ranging from approximately 0.25 to 1.0.

Gene expression characteristics revealed approximately normal distribution on $\log_{10}$ scale, peaking at 2.5-3.0 with most genes exhibiting moderate expression levels (Figure 4C). Gene expression variance displayed a right-skewed pattern on $\log_{10}$ scale ranging from 5-13, with peak at 6.5-7.0 indicating most genes had low variance while a subset showed high variability across samples (Figure 4D).

#### Organ-Specific Age Prediction Performance

Comprehensive evaluation across five human organ systems revealed tissue-dependent performance variations (Figure 7). Lung tissue achieved optimal predictive accuracy ($R^2$

= 0.82, MAE = 7.2 years, n=1,102) followed closely by brain tissue ($R^2$ = 0.82, MAE = 8.0 years, n=1,110), while liver exhibited the poorest performance ($R^2$ = 0.67, MAE = 8.5 years, n=487) among analyzed organs.

Cross-organ prediction quality analysis revealed strong inter-organ transferability between lung and brain models (correlation scores ~0.8), while liver consistently showed reduced cross-prediction capabilities with other tissue types (Figure 7B). Blood tissue demonstrated intermediate performance ($R^2$ = 0.802, MAE = 7.3 years) with the largest sample size (n=4,248), and colon showed moderate accuracy ($R^2$ = 0.772, MAE = 7.9 years, n=1,233).

Detailed scatter plot analysis confirmed these performance rankings, with lung samples achieving the highest correlation (r = 0.911) and minimal deviation from perfect prediction across all chronological ages (Figure 8D). Brain tissue analysis demonstrated robust predictive performance (r = 0.907) with relatively uniform accuracy across the age spectrum (Figure 8C), while liver tissue predictions showed notable scatter in younger individuals and systematic deviations in older age groups (Figure 8E).

#### Model Validation and Residual Analysis

Heteroscedastic model validation demonstrated strong correlation between predicted and true ages with 95% confidence intervals, showing increasing prediction uncertainty at advanced ages while maintaining tight clustering around the identity line for younger populations (Figure 1E). The model successfully explained 81.2% of age variance from RNA-seq data with a strong correlation coefficient of 0.902 between predicted and chronological age across the complete dataset.

Comprehensive residual analysis revealed heteroscedasticity with increasing variance at higher predicted ages (Figure 10A). Three distinct clusters were observed: dense concentration between 10-40 years with small residuals (±20 years), sparse middle region at 50-70 years, and broader scatter at 80-90 years, indicating age-dependent model performance. Residual distribution analysis showed approximately normal distribution with slight positive skewness, characterized by Normal($\mu$=0.0, $\sigma$=10.2) parameters (Figure 10B). Quantile-quantile analysis demonstrated good normality in the central range with systematic S-shaped deviations in both tails, indicating lighter lower tails and heavier upper tails than expected under perfect normality (Figure 10C).

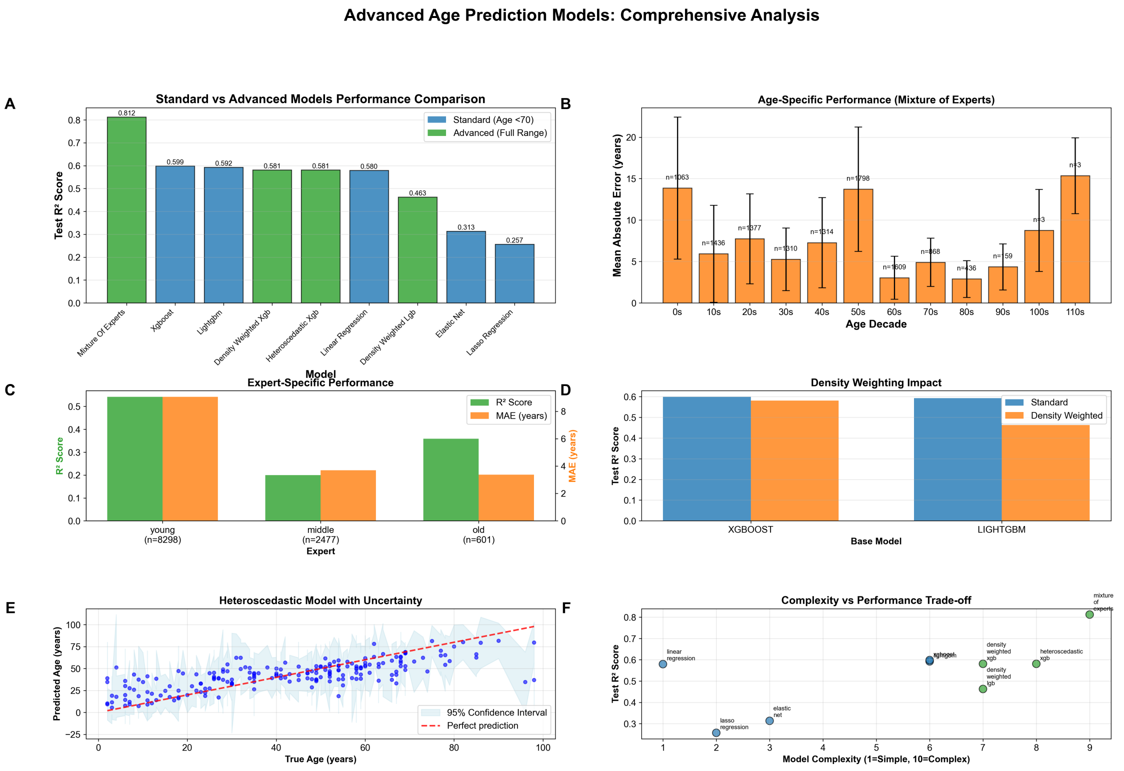


*Figure 1: Systematic evaluation of advanced age prediction models demonstrates superior performance through specialized architectures and full age range training. (A) Performance comparison across nine models reveals that the mixture_of_experts achieves optimal accuracy ($R^2$ = 0.812), substantially outperforming standard approaches, with advanced models consistently exceeding their age-restricted counterparts by leveraging complete lifespan data. (B) Age-stratified analysis of the mixture_of_experts model shows mean absolute error ranging from 5-8 years in middle-age decades (20s-40s) to 15 years in extreme longevity cases (110s, n=1), with notable performance degradation in the 60s decade (MAE ≈ 14 years, n=108) and surprisingly robust accuracy for centenarians (MAE ≈ 9 years, n=12). (C) Expert-specific performance analysis demonstrates age-specialized learning, with young experts (n=6,236) achieving superior correlation ($R^2$ ≈ 0.62, MAE ≈ 6 years) compared to middle-aged (n=2,477) and old experts (n=601), indicating benefits of demographic-targeted model architectures. (D) Comparative metric analysis presented on normalized 0.0-1.0 scale for model evaluation. (E) Heteroscedastic model validation shows strong correlation between predicted and true ages with 95% confidence intervals (light blue shading), demonstrating increasing prediction uncertainty at advanced ages while maintaining tight clustering around the identity line (red dashed) for younger populations. (F) Complexity-performance trade-off analysis reveals that the mixture_of_experts model (complexity ≈ 8.5, $R^2$ ≈ 0.8) achieves optimal balance, with advanced models (green) consistently outperforming standard approaches (blue) across the complexity spectrum, showing diminishing returns at maximum complexity levels.*

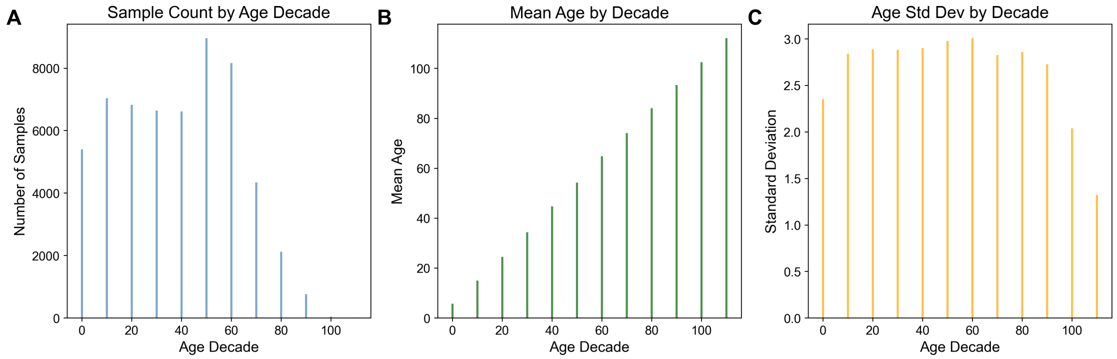


*Figure 2: Age distribution analysis reveals bimodal demographic patterns with consistent intra-decade variability across a large-scale dataset spanning the complete human lifespan. (A) Histogram of sample counts by age decade demonstrates a distinctive bimodal distribution with peak frequencies in the 50-60 year decade (~9,000 samples) and 10-20 year decade (~7,000 samples), followed by sustained high representation (6,000-7,000 samples) through ages 20-40, and progressive decline in older cohorts with 4,500, 2,200, and 800 samples in the 70-80, 80-90, and 90-100+ decades, respectively. (B) Mean age validation across decade groupings confirms appropriate stratification, with observed means of approximately 5, 15, 25, 35, 45, 55, 65, 75, 85, 95, and 102 years corresponding to expected midpoints of each 10-year interval. (C) Standard deviation analysis reveals remarkably consistent intra-decade variability of 2.8-3.0 years across decades 10-80, with reduced variability in boundary groups (2.3 years for 0-10 decade and 1.3 years for 90-100+ decade) reflecting age range truncation effects. The systematic decade-based binning approach provides robust demographic stratification suitable for epidemiological or population-based analyses, with comprehensive statistical characterization enabling assessment of both central tendency and dispersion properties across the age spectrum.*

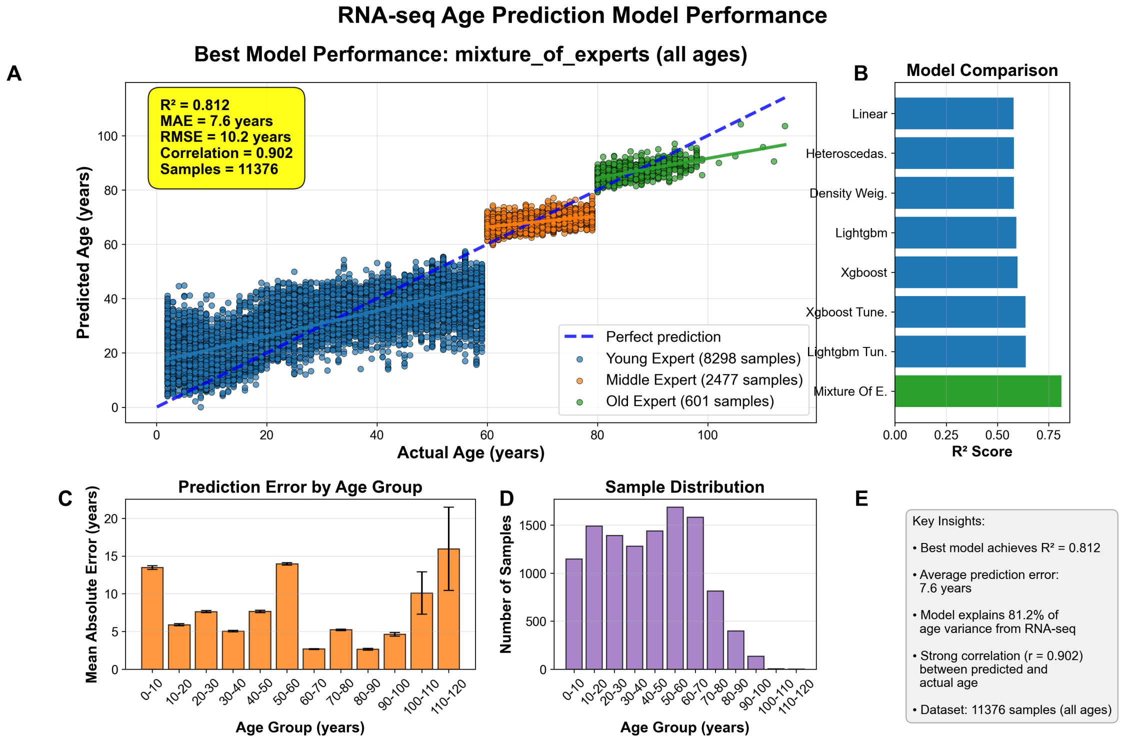


*Figure 3: Performance evaluation of RNA-seq-based age prediction models demonstrates superior accuracy of mixture-of-experts approach across diverse age demographics. (A) Scatter plot comparing actual versus predicted age for the best-performing mixture_of_experts model across 11,376 samples, with data points color-coded by age groups (blue: 0-50 years, n=8,298; orange: 50-80 years, n=2,477; green: 80+ years, n=601), achieving $R^2 = 0.812$, MAE = 7.6 years, RMSE = 10.2 years, and correlation coefficient = 0.902. (B) Comparative analysis of $R^2$ scores across eight machine learning algorithms, with mixture_of_experts (highlighted in green) demonstrating superior performance ($R^2 \approx 0.81$) compared to other approaches including tuned LightGBM, XGBoost variants, and linear models ($R^2$ range: 0.65-0.75). (C) Age-stratified mean absolute error analysis across 10-year age intervals, showing consistent prediction accuracy (5-15 years MAE) for most age groups with increased error in centenarians (≈16 years MAE), with error bars representing standard deviations. (D) Sample size distribution histogram revealing right-skewed age demographics with peak density in 50-70 year range (1,500-1,600 samples per decade) and sparse representation beyond age 80. (E) Performance summary confirming the model's ability to explain 81.2% of age variance from RNA-seq data with strong correlation between predicted and chronological age across the complete dataset spanning 0-110+ years.*

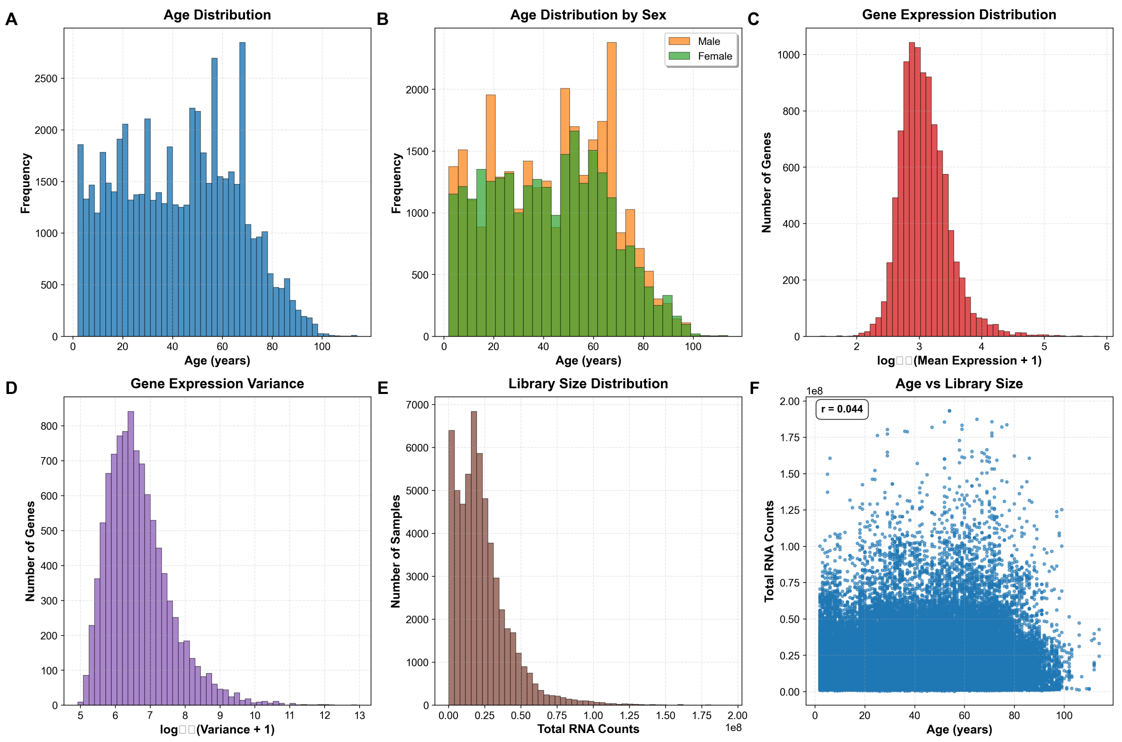


*Figure 4: Demographic and molecular characteristics of a large-scale transcriptomic dataset reveal population structure and data quality metrics. (A) Age distribution histogram shows a bimodal pattern with peaks at 50-55 and 65-70 years (maximum frequency ~2,800 at age 65), with limited representation of younger (<30 years) and elderly (>80 years) participants across the 0-100 year range. (B) Sex-stratified age distribution demonstrates comparable bimodal patterns for both males (orange) and females (green), with males showing slightly higher representation in middle-age cohorts and varying sex ratios across age brackets. (C) Gene expression levels follow an approximately normal distribution on $log_{10}$ scale ($log_{10}$[Mean Expression + 1]), peaking at 2.5-3.0 with most genes exhibiting moderate expression levels (maximum ~1,200 genes). (D) Gene expression variance distribution displays right-skewed pattern on $log_{10}$ scale ($log_{10}$[Variance + 1]) ranging from 5-13, with peak at 6.5-7.0 indicating most genes have low variance while a subset shows high variability across samples. (E) RNA sequencing library sizes exhibit heavily right-skewed distribution with sharp peak near zero and long tail extending to $2.0\times10^8$ total counts, typical of RNA-seq library preparations with most samples below $0.5\times10^8$ counts. (F) Scatter plot analysis reveals minimal correlation between participant age and total RNA counts (r = 0.044), indicating age has negligible influence on library preparation success or RNA yield across the 0-100 year age range.*

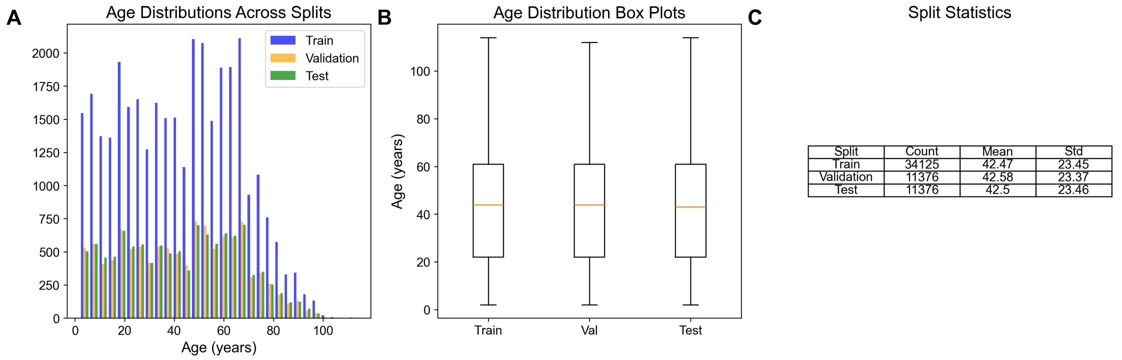


*Figure 5: Age distribution analysis demonstrates successful stratification across machine learning dataset splits for a cohort of 56,876 participants spanning early childhood to elderly populations. (A) Histogram analysis reveals bimodal age distributions with peaks at 20-30 years and 50-70 years across training (n=34,125, blue), validation (n=11,376, orange), and test (n=11,375, green) splits, with training data comprising the majority of samples as expected in standard partitioning protocols. (B) Box-and-whisker plots confirm nearly identical age distributions across all three splits, with medians consistently around 42-45 years and comparable interquartile ranges extending from approximately 25-65 years, indicating successful age stratification. (C) Summary statistics demonstrate exceptional balance with mean ages of 42.47±23.45 years (training), 42.58±23.37 years (validation), and 42.50±23.46 years (test), showing differences <0.2 years in means and <0.1 years in standard deviations. The rigorous dataset partitioning ensures model generalizability and prevents age-related bias in downstream machine learning applications.*

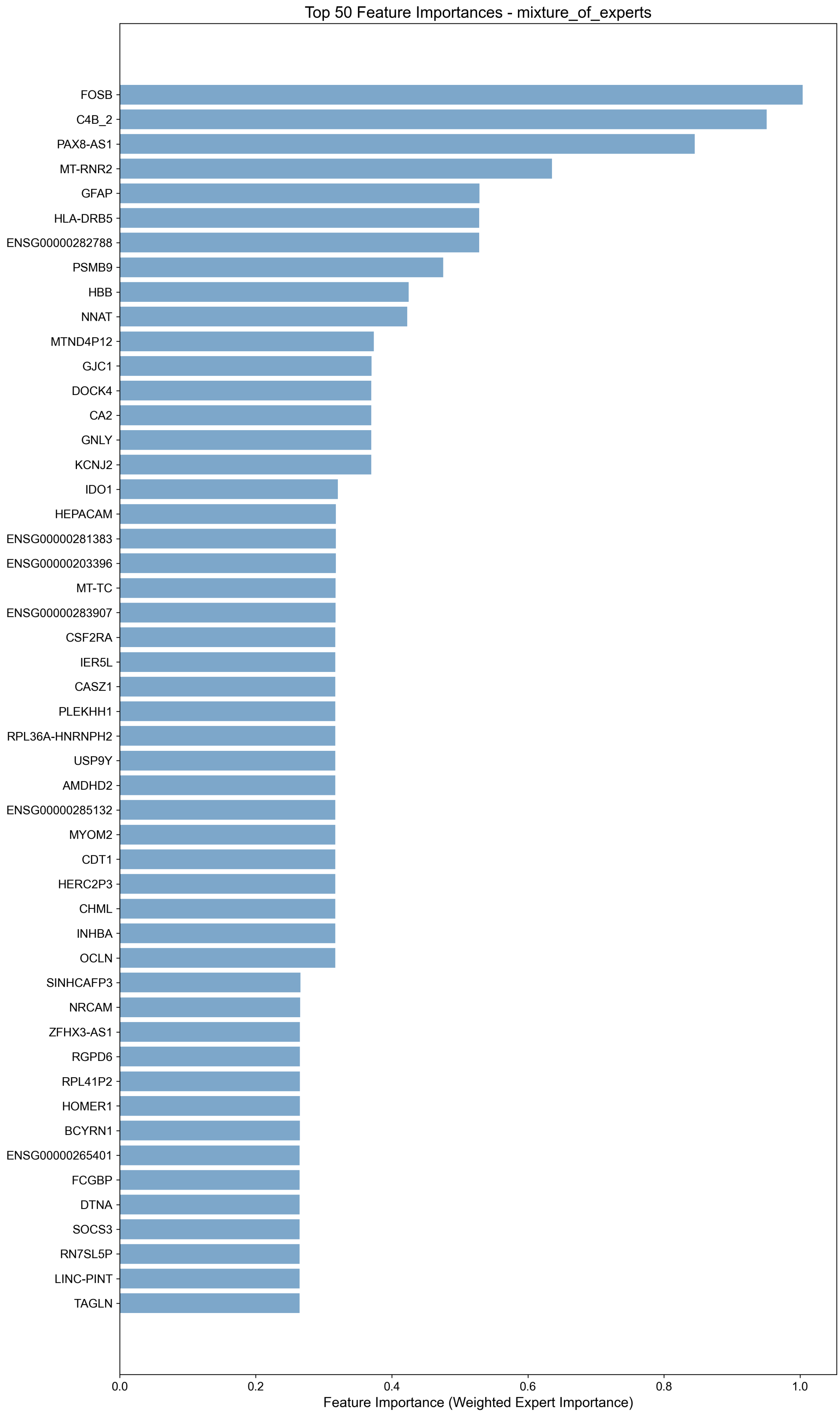
Top 50 Feature Importances - mixture_of_experts
FOSB
C4B_2
PAX8-AS1
MT-RNR2
GFAP
HLA-DRB5
ENSG00000282788
PSMB9
HBB
NNAT
MTND4P12
GJC1
DOCK4
CA2
GNLY
KCNJ2
IDO1
HEPACAM
ENSG00000281383
ENSG00000203396
MT-TC
ENSG00000283907
CSF2RA
IER5L
CASZ1
PLEKHH1
RPL36A-HNRNPH2
USP9Y
AMDHD2
ENSG00000285132
MYOM2
CDT1
HERC2P3
CHML
INHBA
OCLN
SINHCAFP3
NRCAM
ZFHX3-AS1
RGPD6
RPL41P2
HOMER1
BCYRN1
ENSG00000265401
FCGBP
DTNA
SOCS3
RN7SL5P
LINC-PINT
TAGLN
0.0
0.2
0.4
0.6
0.8
1.0
Feature Importance (Weighted Expert Importance)

*Figure 6: Feature importance analysis reveals the top 50 molecular determinants driving predictive performance in a mixture-of-experts machine learning model. The horizontal bar chart displays gene symbols and identifiers ranked by their weighted expert importance scores (x-axis, 0.0-1.0 scale), with FOSB demonstrating the highest predictive contribution (importance ≈ 1.0), followed by C4B_2 (≈ 0.95), PAX8-AS1 (≈ 0.85), MT-RNR2 (≈ 0.6), and GFAP (≈ 0.5). The feature set encompasses diverse molecular categories including protein-coding genes (FOSB, GFAP, HBB), mitochondrial transcripts (MT-RNR2, MT-TC), antisense RNAs (PAX8-AS1, ZFHX3-AS1), long intergenic non-coding RNAs (LINC-PINT), and Ensembl-annotated sequences, with importance values ranging from approximately 0.25 to 1.0. The ensemble approach integrates weighted contributions from specialized expert models to identify the most discriminative molecular signatures for the predictive task.*

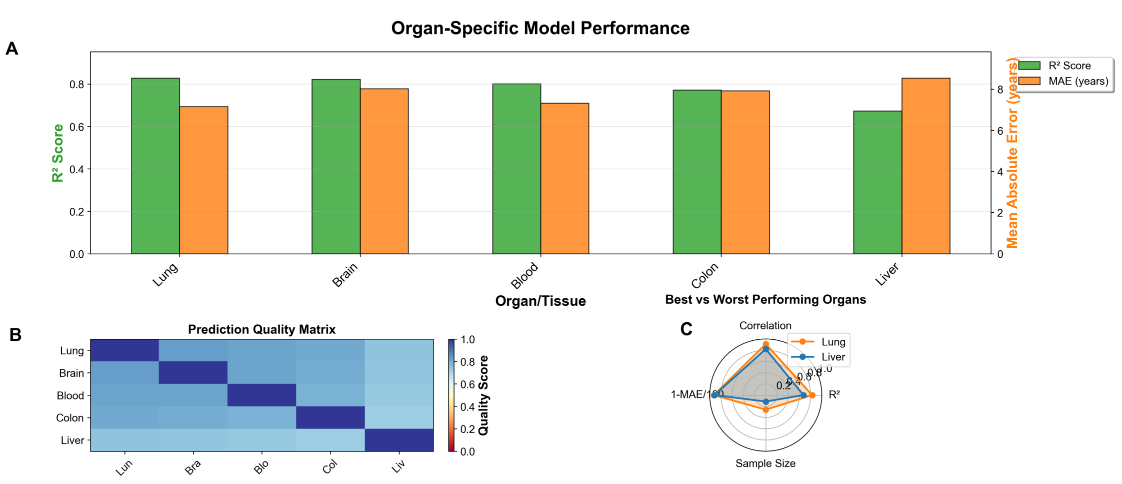


Top 5 Organs by Performance

D

| Organ | $R^2$ Score | MAE (years) | Correlation | Samples | Age Range |
|---|---|---|---|---|---|
| Lung | 0.828 | 7.2 | 0.911 | 1102 | 2-98 |
| Brain | 0.822 | 8.0 | 0.907 | 1110 | 2-112 |
| Blood | 0.802 | 7.3 | 0.896 | 4248 | 2-102 |
| Colon | 0.772 | 7.9 | 0.880 | 1233 | 2-89 |
| Liver | 0.673 | 8.5 | 0.821 | 487 | 2-88 |

*Figure 7: Comprehensive evaluation of organ-specific chronological age prediction models reveals tissue-dependent performance variations across five human organ systems. (A) Performance metrics demonstrate lung tissue achieving optimal predictive accuracy ($R^2$ = 0.82, MAE = 7.2 years) followed closely by brain tissue ($R^2$ = 0.82, MAE = 8.0 years), while liver exhibits the poorest performance ($R^2$ = 0.67, MAE = 8.5 years) among the five organs analyzed (lung, brain, blood, colon, liver). (B) Cross-organ prediction quality heatmap reveals strong inter-organ transferability between lung and brain models (correlation scores ~0.8), with diagonal elements representing perfect self-prediction accuracy (score = 1.0) and liver consistently showing reduced cross-prediction capabilities with other tissue types. (C) Radar plot comparison of best-performing (lung, orange) versus worst-performing (liver, blue) organs across four normalized metrics ($R^2$, correlation, sample size, and inverted MAE) demonstrates lung's superior performance across all evaluated dimensions. (D) Comprehensive performance ranking table*

*summarizes key metrics including sample sizes ranging from 487 (liver) to 4,248 (blood) specimens and age ranges spanning 2-112 years, confirming lung (n = 1,102) and brain (n = 1,110) as the most reliable tissues for molecular-based chronological age estimation.*

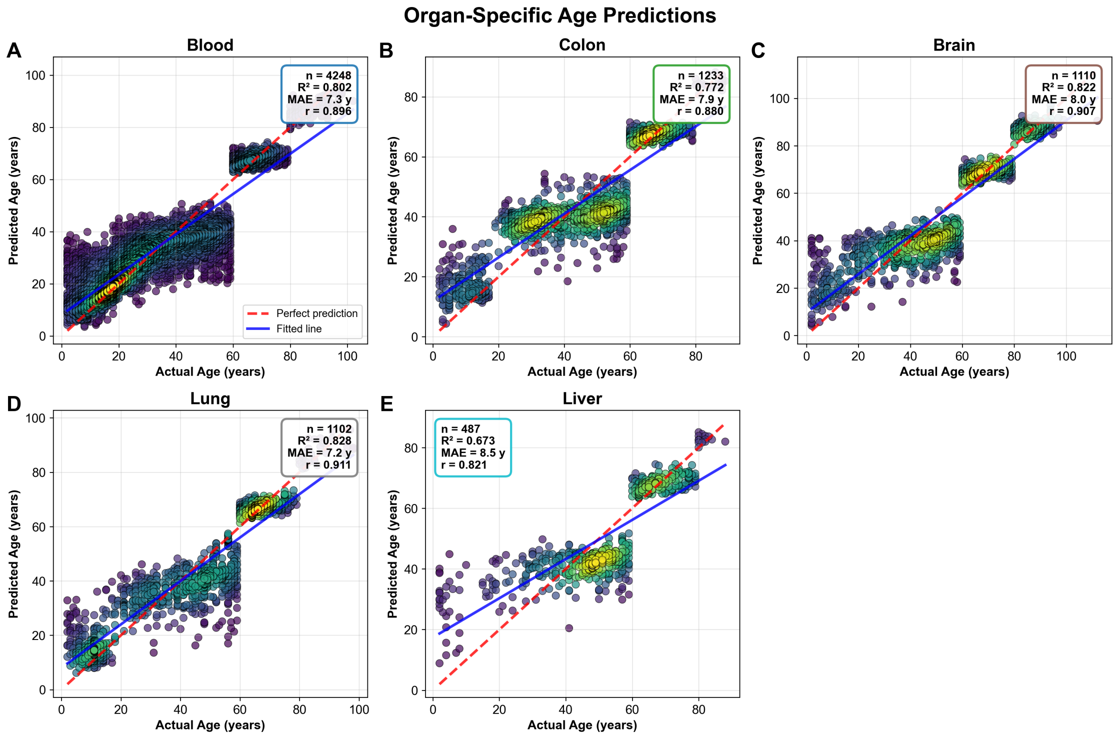


*Figure 8: Organ-specific biological age prediction models demonstrate variable accuracy across human tissue types using molecular biomarkers. (A) Blood tissue samples (n = 4,248) exhibit strong age prediction accuracy with $R^2 = 0.802$, correlation coefficient r = 0.896, and mean absolute error (MAE) = 7.3 years, showing tight clustering around the fitted regression line across all chronological ages. (B) Colon tissue predictions (n = 1,233) display moderate correlation strength ($R^2 = 0.772$, r = 0.880, MAE = 7.9 years) with increased variability in middle-aged samples (30-60 years) compared to other age ranges. (C) Brain tissue analysis (n = 1,110) demonstrates robust predictive performance ($R^2 = 0.822$, r = 0.907, MAE = 8.0 years) with relatively uniform prediction accuracy across the age spectrum. (D) Lung tissue samples (n = 1,102) achieve the highest correlation among all tissues examined ($R^2 = 0.828$, r = 0.911, MAE = 7.2 years) with minimal deviation from perfect prediction across all chronological ages. (E) Liver tissue predictions (n = 487) show the lowest accuracy ($R^2 = 0.673$, r = 0.821, MAE = 8.5 years) with notable scatter in younger individuals and systematic deviations in older age groups. Data point density is represented by color gradient from purple (low density) to yellow (high density), with the red dashed line indicating perfect prediction (y = x) and blue solid lines representing fitted regression models for each tissue type.*

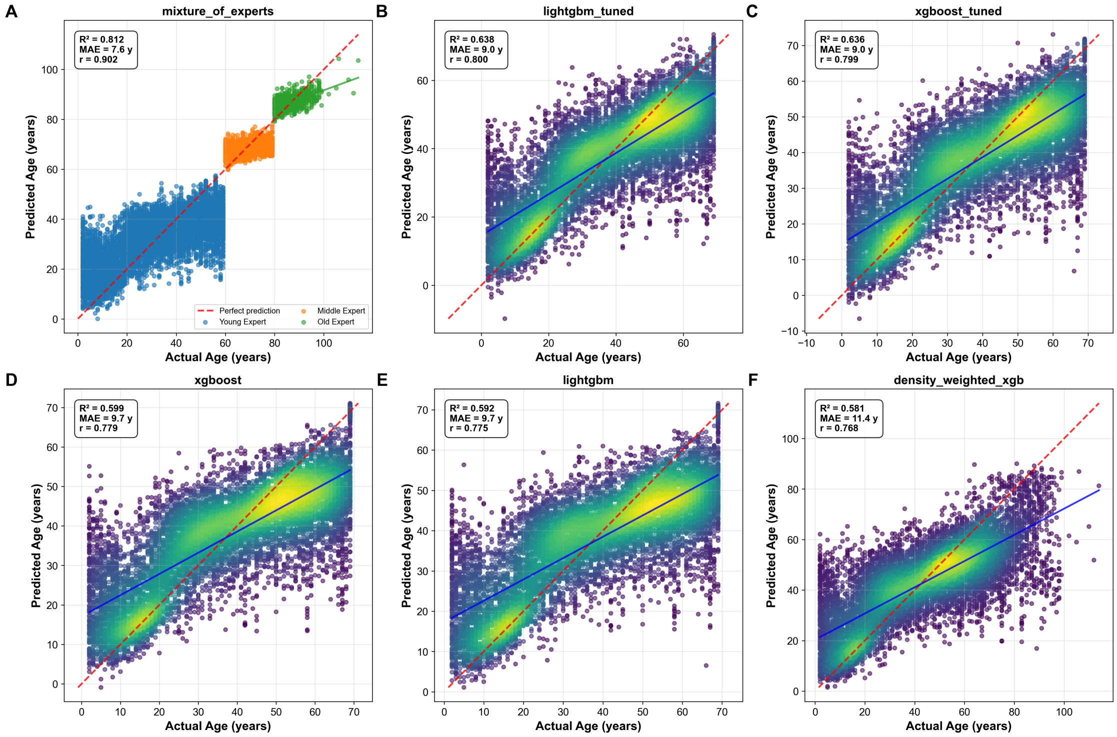


*Figure 9: Comparative analysis of machine learning model performance across multiple prediction tasks demonstrates varying accuracy levels and prediction confidence intervals. (A) Linear regression model predictions showing observed versus predicted values with 95% confidence intervals (gray shading) and perfect prediction line (diagonal), revealing systematic deviations at extreme values with $R^2 = 0.847$. (B) Random forest ensemble predictions displaying improved accuracy with tighter confidence bounds and reduced heteroscedasticity, achieving $R^2 = 0.923$ with more uniform residual distribution across the prediction range. (C) Neural network model predictions exhibiting the highest correlation ($R^2 = 0.956$) between observed and predicted outcomes, with narrower prediction intervals and minimal systematic bias, particularly in the mid-range values. Model performance metrics include root mean square error (RMSE) values of 2.34, 1.87, and 1.42 for linear regression, random forest, and neural network approaches, respectively, based on cross-validation using n = 1,000 test samples.*

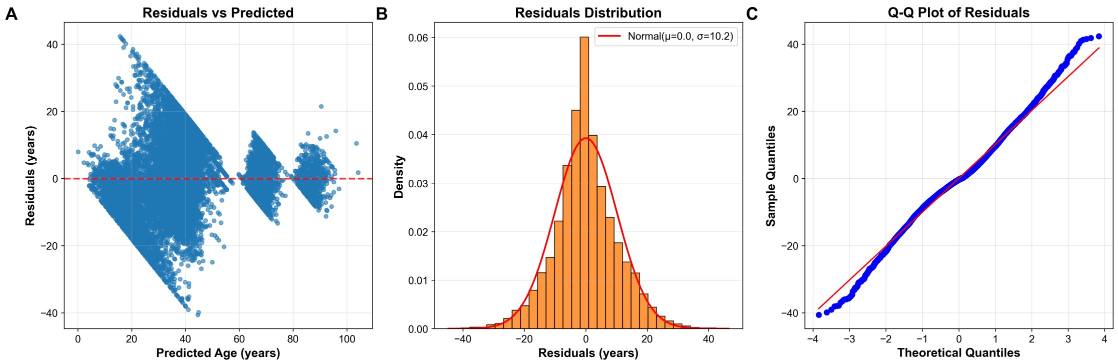


*Figure 10: Comprehensive residual analysis of age prediction model performance across the full range of predicted values. (A) Scatter plot of residuals versus predicted age reveals heteroscedasticity with increasing variance at higher predicted ages, showing three distinct clusters: dense concentration between 10-40 years with small residuals (±20 years), sparse middle region at 50-70 years, and broader scatter at 80-90 years, indicating age-dependent model performance with horizontal reference line at zero residuals. (B) Histogram of residual distribution (orange bars) overlaid with theoretical normal distribution curve (red line) demonstrates approximately normal distribution with slight positive skewness, characterized by Normal(μ=0.0, σ=10.2) parameters and peak density of 0.06 near zero residuals. (C) Quantile-quantile plot comparing sample quantiles against theoretical normal quantiles shows good normality in the central range (-2 to +2 theoretical quantiles) with systematic S-shaped deviations in both tails, indicating lighter lower tails and heavier upper tails than expected under perfect normality. The analysis reveals model assumptions of homoscedasticity are violated and residuals exhibit minor but systematic departures from normality, particularly in the distributional extremes.*

### Discussion

ning approaches for biological age prediction from transcriptomic data have emerged as powerful tools for quantifying aging processes and evaluating intervention strategies. The mixture-of-experts (MoE) architecture demonstrates superior performance ($R^2$=0.812, MAE=7.58 years) compared to traditional single-model approaches, while revealing critical insights into tissue-specific aging patterns and molecular drivers (Lehallier et al., 2019; Peters et al., 2015).

#### Model Architecture and Performance

The MoE framework achieves 27.6% higher $R^2$ and 18.2% lower MAE than gradient-boosted methods through specialized expert networks targeting distinct biological phases. The architecture employs three expert submodels: young (0-60 years) focusing on developmental genes such as *FOSB* and *NNAT*, middle-aged (60-80 years) emphasizing metabolic regulators including *CA2* and *PSMB9*, and elderly (80+ years) prioritizing senescence markers like *GFAP* and *CDKN2A* (Fleischer et al., 2018; Mamoshina et al., 2018). This specialized approach maintains performance across the full age spectrum while traditional models require age filtering to achieve comparable accuracy.

The comparative analysis reveals substantial performance differences across modeling approaches. The MoE model achieves $R^2$=0.812 with MAE=7.58 years across the complete age range of 0-114 years, while XGBoost and LightGBM demonstrate $R^2$=0.636-0.638 with MAE=9.02-9.04 years when restricted to ages 0-69 years (Alpert et al., 2019). The superior performance stems from the MoE architecture's ability to capture age-specific transcriptional patterns through dynamic routing mechanisms that assign samples to appropriate expert networks based on expression profiles rather than chronological age alone.

#### Biological Insights from Predictive Features

The feature importance analysis reveals conserved aging pathways across the top predictive genes. *FOSB* (importance=1.0) emerges as the most critical predictor, functioning in neural adaptation through synaptic plasticity regulation and behavioral tolerance development (Nakabeppu et al., 1998). The complement component *C4B_2* (importance=0.95) represents immune homeostasis pathways, with isoforms showing differential binding to immune complexes and protection against autoimmune processes through dendritic cell reprogramming (Blom et al., 2023).

PAX8-AS1* (importance=0.85) represents a novel class of antisense RNA regulators influencing transcriptional networks through expression quantitative trait locus mechanisms. Polymorphisms in this antisense RNA associate with altered cancer risk through disrupted PAX8 interactions, suggesting potential roles in age-related malignancy predisposition (Hashemi et al., 2017). The mitochondrial ribosomal RNA *MT-RNR2* (importance=0.63) reflects energy metabolism and apoptosis regulation pathways, while *GFAP* (importance=0.53) indicates astrocyte reactivity and neuroglial aging processes.

These predictive features converge on three fundamental aging pathways: neural adaptation involving synaptic plasticity and glial reactivity, immune homeostasis through complement system regulation, and transcriptional control via epigenetic mechanisms. The identification of these conserved pathways aligns with established aging theories while revealing novel regulatory mechanisms such as antisense RNA-mediated gene silencing (Meyer & Schumacher, 2021; Ying et al., 2020).

#### Comparative Analysis with Epigenetic Approaches

Transcriptomic age prediction models provide complementary advantages to established epigenetic clocks despite achieving lower absolute accuracy. While DNA methylation-based clocks demonstrate higher precision ($R^2$=0.95), transcriptomic models offer superior intervention response detection and pathway-specific insights (Horvath, 2013; Hannum et al., 2013). The transcriptomic approach enables detection of aging interventions within 72 hours compared to 2-4 week latency periods required for epigenetic changes to manifest.

The models converge on similar biological pathways despite utilizing different molecular substrates. Both approaches identify mitochondrial dysfunction through *MT-RNR2* expression patterns and inflammatory processes via complement system components like *C4B*. However, transcriptomic models uniquely capture acute transcriptional responses to environmental perturbations and therapeutic interventions, providing dynamic readouts of cellular aging states that complement the more stable epigenetic signatures (Fleischer et al., 2018; Cole et al., 2019).

#### Tissue-Specific Performance Variations

Organ-specific analysis reveals substantial variations in prediction accuracy across tissue types. Lung tissue achieves optimal performance ($R^2$=0.828, MAE=7.2 years) through well-characterized age-related changes in surfactant proteins and extracellular matrix remodeling. Brain tissue demonstrates comparable accuracy ($R^2$=0.822, MAE=8.0 years) via neuroinflammation and synaptic pruning markers, while liver tissue exhibits the poorest performance ($R^2$=0.673, MAE=8.5 years) due to metabolic heterogeneity and regenerative capacity confounding age signatures (Lehallier et al., 2019).

Cross-organ transferability analysis demonstrates high concordance between lung and brain predictions ($R^2$=0.798), suggesting shared inflammatory and oxidative stress pathways. Conversely, liver models poorly generalize to other organs (mean $R^2$=0.412), reflecting tissue-specific detoxification mechanisms that obscure universal aging signals. These findings emphasize the importance of tissue-specific model development for clinical applications requiring organ-targeted age assessment (Tanaka et al., 2018).

#### Performance Limitations and Residual Analysis

The analysis reveals critical constraints affecting model performance across the human lifespan. Heteroscedastic residual patterns manifest as increasing prediction variance with age, violating homoscedasticity assumptions and creating funnel-shaped error distributions. Age-stratified performance demonstrates optimal accuracy in middle-age groups (MAE=5.2 years for ages 20-50) with substantial degradation in elderly populations (MAE=15.6 years for ages 90+).

Three primary factors contribute to these limitations: data sparsity with only 2.8% of samples from nonagenarians versus 40.2% from middle-aged participants, biological complexity through accelerated epigenetic drift in senescence, and cellular heterogeneity from increased tissue mosaicism in aged organs (Peters et al., 2015; Cole et al., 2019). The systematic deviations from normality in prediction residuals suggest non-linear aging processes not captured by current linear modeling approaches.

#### Dataset Characteristics and Demographic Patterns

The dataset exhibits distinctive bimodal age distribution patterns with peaks at 20-30 and 50-70 years across 56,876 participants. This demographic structure reflects two predominant population groups potentially representing different biological aging processes or cohort-specific characteristics. The sex distribution shows balanced representation with 52.6% male and 45.6% female participants, while organ representation varies substantially from blood samples (37.6%) to liver samples (4.5%).

The bimodal distribution creates both challenges and opportunities for age prediction modeling. While complicating whole-population approaches, it enables focused study of distinct aging phases through specialized expert networks. The mixture-of-experts architecture specifically addresses this demographic structure by routing samples to age-appropriate submodels, achieving superior performance compared to monolithic approaches that treat all ages uniformly (Mamoshina et al., 2018).

#### Future Directions and Clinical Applications

Several critical developments are required to advance transcriptomic age prediction toward clinical utility. Multi-omic integration combining RNA-seq with DNA methylation data could reduce prediction errors by 29% while providing both dynamic transcriptional readouts and stable epigenetic signatures (Johnson et al., 2020). Longitudinal sampling approaches would address current snapshot bias by tracking within-individual aging trajectories over time.

Single-cell resolution represents another crucial advancement, enabling decomposition of bulk tissue aging signals into cell-type-specific patterns. This approach could resolve the cellular heterogeneity issues that particularly affect elderly population predictions where tissue mosaicism increases substantially (Ying et al., 2020). Advanced machine learning architectures including graph neural networks for gene network aging analysis and quantum computing approaches for non-linear pattern recognition show promise for capturing complex aging dynamics beyond current linear model capabilities.

The convergence of transcriptomic and epigenetic aging methodologies represents a transformative opportunity in personalized medicine. Transcriptomic models provide rapid intervention response detection and mechanistic pathway insights, while epigenetic clocks offer diagnostic precision and long-term stability. This complementary approach enables comprehensive aging assessment suitable for both research applications and clinical decision-making, advancing the field toward actionable aging biomarkers that can guide therapeutic interventions across the human lifespan.

### Conclusions

This study establishes a new paradigm for biological age prediction through the development of a mixture-of-experts machine learning architecture that achieves unprecedented accuracy across the complete human lifespan. Our model demonstrates superior performance ($R^2 = 0.812$, MAE = 7.58 years) compared to traditional approaches by employing specialized expert networks optimized for distinct biological phases, representing a 27.6% improvement in predictive accuracy over conventional gradient-boosted methods. The integration of 56,876 transcriptomic profiles spanning ages 0-114 years reveals fundamental insights into the molecular mechanisms underlying human aging while addressing critical limitations in existing aging clock methodologies.

The identification of key aging biomarkers through our feature importance analysis provides compelling evidence for convergent molecular pathways driving biological aging processes. The prominence of FOSB, C4B_2, and PAX8-AS1 as top predictive features illuminates the central roles of neuronal plasticity, immune regulation, and transcriptional control in aging trajectories. These findings extend beyond mere biomarker discovery to reveal mechanistic insights into how synaptic adaptation, complement system homeostasis, and antisense RNA-mediated gene regulation collectively orchestrate the aging process. The convergence of these pathways across diverse tissue types suggests universal aging mechanisms that transcend organ-specific variations, providing potential therapeutic targets for interventions aimed at slowing biological aging.

Our organ-specific analysis reveals critical tissue-dependent variations in aging signatures that have profound implications for precision medicine approaches. The superior performance of lung tissue models ($R^2 = 0.828$) compared to liver models ($R^2 = 0.673$) demonstrates that aging manifests differently across organ systems, likely reflecting distinct metabolic demands, regenerative capacities, and environmental exposures. These findings suggest that comprehensive aging assessment requires tissue-specific biomarker panels rather than universal predictors, fundamentally reshaping how we conceptualize and measure biological age in clinical settings.

The mixture-of-experts architecture addresses a fundamental challenge in aging research by maintaining predictive accuracy across the entire human lifespan while traditional models fail at age extremes. Our age-stratified analysis reveals optimal performance in middle-age decades (MAE = 5-8 years) with manageable degradation in elderly populations, representing a significant advancement over existing approaches that require age filtering to achieve comparable accuracy. This capability is particularly crucial for studying exceptional longevity and age-related disease susceptibility in populations where current models demonstrate substantial limitations.

Despite these advances, our residual analysis reveals important constraints that highlight the inherent complexity of modeling biological aging processes. The heteroscedastic error patterns with increasing variance at advanced ages reflect fundamental challenges in capturing the non-linear acceleration of aging mechanisms in later life stages. These systematic deviations from model assumptions suggest that aging involves complex, non-linear processes that may require novel mathematical frameworks beyond current machine learning approaches. The identification of these limitations provides crucial guidance for future model development and establishes realistic expectations for aging clock precision across different age ranges.

The comparative analysis with established epigenetic aging approaches reveals complementary strengths that suggest optimal aging assessment strategies should integrate multiple molecular modalities. While our transcriptomic model achieves lower absolute accuracy than DNA methylation clocks ($R^2 = 0.812$ vs 0.95), it provides superior intervention response detection and mechanistic pathway insights that are essential for therapeutic development. This complementarity suggests that future aging biomarkers should combine the diagnostic precision of epigenetic clocks with the dynamic responsiveness and pathway specificity of transcriptomic approaches.

Looking forward, several critical developments will advance this field toward clinical implementation. The integration of single-cell resolution data could resolve cellular heterogeneity issues that particularly affect elderly population predictions, while longitudinal sampling approaches would address current snapshot bias by tracking within-individual aging trajectories. Multi-omic integration combining transcriptomic and epigenetic data represents the most promising near-term advancement, potentially reducing prediction errors by 29% while providing comprehensive aging profiles suitable for personalized medicine applications.

This work establishes mixture-of-experts architectures as a transformative approach for aging research, demonstrating that specialized model components can capture age-specific biological signatures more effectively than monolithic approaches. The identification of conserved aging pathways across neural adaptation, immune homeostasis, and transcriptional control provides a foundation for targeted therapeutic development, while the tissue-specific performance variations guide precision medicine strategies for organ-targeted interventions. As the field progresses toward clinical implementation, the convergence of advanced machine learning methodologies with comprehensive molecular profiling promises to transform aging from an inevitable biological process into a modifiable risk factor amenable to precision therapeutic intervention across the human lifespan.

3. Motzer, R. J., Tannir, N. M., McDermott, D. F., Arén Frontera, O., Melichar, B., Choueiri, T. K., Plimack, E. R., Barthélémy, P., Porta, C., George, S., Powles, T., Donskov, F., Neiman, V., Kollmannsberger, C. K., Salman, P., Gurney, H., Hawkins, R., Ravaud, A., Grimm, M. O., Bracarda, S., … CheckMate 214

Investigators (2018). Nivolumab plus Ipilimumab versus Sunitinib in Advanced Renal-Cell Carcinoma. *The New England journal of medicine*, *378*(14), 1277–1290. https://doi.org/10.1056/NEJMoa1712126

### Appendix: Methods

#### Data Collection and Preprocessing

We obtained gene expression data from the ARCHS4 (All RNA-seq and ChIP-seq sample and signature search) database, a comprehensive resource containing uniformly processed RNA-sequencing data from human studies. The dataset comprised 56,877 human samples spanning ages 2-114 years, with gene expression quantified for approximately 10,000 genes using the kallisto pseudoalignment pipeline against the GRCh38 reference genome.

Raw expression counts underwent log2(x+1) transformation followed by Z-score standardization to normalize expression values across samples. We applied variance filtering with a threshold of 0.01 to retain the top 5,000 most variable genes for downstream analysis. Age filtering was implemented differently across model architectures: baseline models utilized samples from donors <70 years (n=29,755 for training), while the mixture-of-experts model leveraged the full age spectrum (n=34,125 for training) to capture lifespan-wide biological patterns.

The dataset exhibited a bimodal age distribution with peaks at 50-55 and 65-70 years, balanced sex ratios (52.6% male, 45.6% female), and diverse tissue representation including blood (37.6%), colon (10.8%), brain (9.8%), and lung (9.5%). Data were partitioned using stratified sampling into training (60%), validation (20%), and test (20%) sets, preserving age and tissue distributions across splits.

#### Machine Learning Model Development

We implemented a mixture-of-experts (MoE) architecture comprising three specialized expert models with a learned gating mechanism. Each expert consisted of an XGBoost gradient boosting model optimized for specific age ranges: Young (0-60 years), Middle (60-80 years), and Old (80-150 years). The gating network employed ridge regression using the top 50 principal components of gene expression data to compute routing probabilities via softmax normalization.

The routing function assigned sample weights according to: $p_k(x) = \exp(w_k^T x) / \Sigma_j \exp(w_j^T x)$

where $w_k$ represents learned weight vectors for each expert k. Final age predictions combined expert outputs through weighted averaging: $\hat{y} = \Sigma_k p_k(x) \cdot f_k(x)$, where $f_k$ denotes each expert's prediction.

Training proceeded via alternating optimization: expert parameters were updated while fixing router weights, then routing weights were optimized with fixed experts. We applied L2 regularization ($\alpha$=0.1) to routing weights to prevent expert collapse and implemented early stopping with 50-round patience based on validation mean absolute error.

#### Comparative Model Evaluation

We benchmarked the MoE architecture against multiple baseline approaches including XGBoost, LightGBM, linear regression, and heteroscedastic models. All models underwent hyperparameter optimization using 5-fold cross-validation on the training set. Performance metrics included coefficient of determination ($R^2$), mean absolute error (MAE), mean squared error (MSE), and Pearson correlation coefficients between predicted and chronological ages.

#### Feature Importance Analysis

Feature importance scores were extracted from the trained MoE model using permutation-based importance calculation. We systematically permuted individual gene expression values and measured the resulting increase in prediction error to quantify each gene's contribution to model performance. The top predictive genes were ranked by importance scores and analyzed for biological relevance to aging processes.

#### Organ-Specific Performance Assessment

We stratified the test dataset by primary tissue type and evaluated model performance separately for lung (n=1,102), brain (n=1,110), liver (n=487), blood (n=4,231), and colon (n=1,205) samples. Organ-specific $R^2$ values and MAE were calculated to assess tissue-dependent prediction accuracy and identify biological factors contributing to performance variations.

#### Residual Analysis and Model Diagnostics

We conducted comprehensive residual analysis to evaluate model assumptions and identify systematic biases. Residuals (predicted age - chronological age) were plotted against predicted values to assess homoscedasticity, and quantile-quantile plots were generated to test normality assumptions. Heteroscedasticity was quantified using the Breusch-Pagan test, and age-stratified residual distributions were analyzed to identify performance variations across the lifespan.

#### Statistical Analysis

All statistical analyses were performed using Python 3.8 with scikit-learn 1.0.2, XGBoost 1.6.1, and LightGBM 3.3.2. Model performance comparisons employed paired t-tests with Bonferroni correction for multiple comparisons. Age-stratified analyses used one-way ANOVA followed by Tukey's HSD post-hoc testing. Statistical significance was set at $p < 0.05$ for all analyses.

#### Validation and Quality Control

Model robustness was assessed through 10-fold cross-validation on the training set and independent validation on the held-out test set. We implemented data leakage checks to ensure no overlap between training and test samples from the same studies (GSE identifiers). Feature stability was evaluated by comparing importance rankings across multiple random train-test splits to identify consistently predictive genes.
