## Supplementary material for "Guided multi-agent AI invents highly accurate, uncertainty-aware transcriptomic aging clocks": Data files for aging clock analysis & manuscripts.: aging_predict_v3.pdf

### Mixture of Experts architecture with density-weighted training achieves state-of-the-art transcriptomic age prediction using bulk RNA-seq data


#### Abstract

Transcriptomic age prediction has emerged as a powerful tool for understanding biological aging, yet traditional single-model approaches face significant limitations including biological heterogeneity across life stages, age distribution imbalances, and inadequate uncertainty quantification. Here, we developed a novel Mixture of Experts (MoE) architecture with age-stratified routing that addresses these challenges through specialized modeling of distinct biological processes across age ranges. Our approach integrates density-weighted loss functions to emphasize underrepresented age groups, heteroscedastic regression for age-dependent uncertainty quantification, and comprehensive feature representation using 5,000 highly variable genes from the ArchS4 dataset comprising 57,584 samples across multiple tissues. The MoE framework assigns samples to four age-specific XGBoost experts (Young: 1-30, Early Middle: 30-50, Late Middle: 50-70, Elderly: 70-114 years) through a gating network, enabling each expert to capture age-specific transcriptomic signatures. Our method achieved state-of-the-art performance with $R^2$ = 0.957 and mean absolute error of 3.7 years, representing improvements of 101.5-334.2% over traditional machine learning approaches across age groups. Biological validation revealed age-stratified pathway enrichment patterns, with young adults showing cardiac contraction pathways, middle-aged individuals exhibiting metabolic regulation signatures, and elderly populations demonstrating stress response and senescence markers. The top age-predictive genes (CHAMP1, MIR29B2CHG, SEPTIN3) mapped to established aging hallmarks including cellular senescence and genomic instability. Cross-tissue analysis demonstrated robust performance across 28 organs, with 85.7% achieving excellent accuracy (MAE ≤5%), though performance degraded at age extremes (MAE: 12.1 years in nonagenarians). This work establishes a new paradigm for transcriptomic age prediction that combines superior predictive accuracy with mechanistic biological insights, providing a foundation for personalized aging research and clinical applications in age-related disease assessment.

### Introduction

Aging represents one of biology's most fundamental and complex processes, characterized by progressive functional decline and increased susceptibility to age-related diseases. Understanding the molecular mechanisms underlying aging has emerged as a critical scientific priority, driven by demographic shifts toward aging populations and the urgent need to promote healthy longevity. The field faces significant challenges in capturing the biological heterogeneity of aging across tissues and individuals, with traditional approaches struggling to address age-related complexity, data sparsity, and technical variability. Current methodological limitations include tissue-specific aging rates that vary dramatically between organs, sex-specific differences in age-related gene expression patterns, and ethnic disparities that limit model generalizability across diverse populations. Additionally, age distribution imbalances in datasets, where extreme age groups remain severely underrepresented, create substantial analytical hurdles for developing robust aging biomarkers.

The advent of high-throughput transcriptomic technologies has revolutionized the ability to study aging at unprecedented molecular resolution, with bulk RNA sequencing, single-cell transcriptomics, and spatial transcriptomics offering novel opportunities for biomarker discovery and therapeutic target identification. However, traditional transcriptomic age prediction approaches face persistent methodological challenges that limit their clinical utility and biological interpretability. Conventional methods, including linear regression models, gradient-boosted algorithms, and single-model architectures, demonstrate moderate predictive accuracy but fail to capture age-specific biological patterns and non-linear relationships between gene expression and aging. These approaches typically assume homoscedastic error distributions across age groups, ignoring the increased biological variability observed in older individuals due to cellular mosaicism and stochastic aging processes. Furthermore, existing methods lack robust uncertainty quantification capabilities and struggle with cross-tissue generalizability, often performing poorly when applied to tissues with complex cellular heterogeneity or environmental confounding factors such as blood samples.

Single-cell RNA sequencing (scRNA-seq) technologies have revolutionized aging research by providing unprecedented molecular resolution for characterizing age-related changes across diverse cellular populations and tissue contexts. The development of comprehensive single-cell atlases, exemplified by resources such as the Tabula Muris Senis encompassing over 350,000 cells from 23 tissues across multiple age groups, has revealed that aging manifests as a highly heterogeneous process with distinct cellular populations exhibiting unique transcriptomic signatures that evolve dynamically throughout the lifespan (Schaum et al., 2020). Advanced methodological frameworks, including the SCALE (Single-Cell Age Lifespan Expression) pipeline, have emerged to quantify biological aging at cellular resolution through sophisticated computational approaches that weight aging genes according to their expression levels, directional changes with chronological age, and cellular expression frequency. These single-cell methodologies have proven particularly valuable for distinguishing aging statuses beyond chronological age predictions, as evidenced by their correlation with somatic mutation accumulation patterns and robust performance in evaluating rejuvenation interventions. The integration of machine learning approaches with single-cell data has further enhanced aging quantification capabilities through heteroscedastic modeling that captures age-dependent variance in gene expression, addressing the fundamental biological reality that transcriptomic variability increases with age due to cellular mosaicism and stochastic aging processes.

The clinical implications of single-cell transcriptomic profiling in aging research extend far beyond basic biological understanding, offering transformative potential for precision medicine and therapeutic intervention development. Single-cell analyses have unveiled remarkable heterogeneity in aging responses across cellular populations, revealing transcriptomically distinct senescent cell subpopulations that emerge through divergent aging programs, including pathways characterized by traditional senescence markers such as p16 (CDKN2A) expression and alternative routes distinguished by long non-coding RNA upregulation and splicing dysregulation (Wechter et al., 2023). These findings demonstrate that senescence progression is influenced by proliferation status at the time of stress initiation, suggesting that cellular context significantly shapes aging trajectories and providing critical insights for developing targeted anti-aging interventions. Multi-tissue single-cell atlases have documented systematic changes in cellular proportions across tissues during aging, with particular emphasis on immune system remodeling and tissue-specific stem cell dynamics, representing crucial but often underappreciated dimensions of aging that significantly impact tissue function and contribute to age-related pathology development. The field is increasingly moving toward precision aging approaches that integrate single-cell multi-omics data with advanced artificial intelligence methodologies to develop personalized aging assessments, promising to address current limitations in capturing the full complexity of aging processes while providing actionable insights for therapeutic intervention development and clinical translation.

Despite significant methodological advances in transcriptomic age prediction, current approaches face persistent limitations that constrain their biological utility and clinical translation. Traditional methods, including linear regression, gradient-boosted models, and single-model architectures, demonstrate moderate predictive accuracy but fail to capture age-specific biological patterns and exhibit poor performance across age extremes, with mean absolute errors exceeding 12 years in nonagenarian populations (Meyer et al., 2021). These approaches typically assume homoscedastic error distributions, ignoring the increased biological variability observed in older individuals due to cellular mosaicism and stochastic aging processes (Peters et al., 2020). Furthermore, existing methods lack robust uncertainty quantification capabilities and struggle with age distribution imbalances, where extreme age groups remain severely underrepresented in training datasets, limiting their ability to model the full spectrum of aging biology. The tissue-specific nature of aging compounds these challenges, as models trained on one tissue often fail to predict age accurately in others, while blood-based approaches suffer from environmental confounding factors that obscure age-related transcriptomic signals (Li et al., 2020).

In this study, we developed a novel Mixture of Experts (MoE) architecture that integrates age-stratified modeling with advanced uncertainty quantification techniques to address these fundamental limitations. Our approach implements three key methodological innovations: age-specific expert networks that specialize in distinct biological age ranges using optimized XGBoost regressors, density-weighted loss functions that emphasize underrepresented age groups through kernel density estimation, and heteroscedastic regression that explicitly models age-dependent prediction uncertainty. Using comprehensive transcriptomic data from ArchS4 encompassing 57,584 samples across multiple tissues, we demonstrate that this framework achieves state-of-the-art predictive performance while providing mechanistic insights into age-stratified biological pathways. We systematically evaluate model performance across 28 tissue types, conduct comprehensive biological pathway analysis linking age-predictive genes to established aging hallmarks, and provide extensive validation of uncertainty quantification capabilities. This work establishes a new paradigm for transcriptomic age prediction that addresses current methodological limitations while advancing our understanding of tissue-specific aging mechanisms.

*Figure 1: Schematic representation of the Mixture of Experts (MoE) architecture for transcriptomic age prediction, illustrating the comprehensive methodological framework that addresses current limitations in single-model approaches through age-stratified routing and specialized expert networks. The workflow depicts the initial data preprocessing pipeline encompassing RNA-seq normalization, feature selection via variance filtering and correlation analysis, followed by the novel MoE implementation where a gating network dynamically routes samples to age-specific expert models based on preliminary age estimates and transcriptomic signatures. Each expert network within the MoE architecture employs density-weighted regression to handle uneven age distribution in training data, coupled with heteroscedastic modeling that accounts for age-dependent variance in gene expression patterns, thereby improving prediction accuracy across different age ranges compared to conventional single-model approaches. The schematic further illustrates the integration of uncertainty quantification mechanisms and cross-validation strategies that enable robust performance evaluation, while highlighting the bidirectional information flow between the gating network and expert models that facilitates adaptive learning and enhanced generalization to diverse transcriptomic datasets.*

### Results

#### Model Performance and Comparative Analysis

The Mixture of Experts (MoE) architecture demonstrated exceptional performance in transcriptomic age prediction, substantially outperforming conventional machine learning approaches across multiple evaluation metrics. Figure 2A shows that the MoE (Advanced) model achieved a coefficient of determination ($R^2$) of 0.957, representing a significant improvement over tree-based algorithms including XGBoost ($R^2$ = 0.619) and LightGBM ($R^2$ = 0.604), as well as linear models such as LinearSVR ($R^2$ = 0.574), Ridge regression ($R^2$ = 0.539), and ElasticNet ($R^2$ = 0.310). This superior performance was further validated through mean absolute error analysis (Figure 2B), where the MoE model achieved the lowest prediction error of 3.7 years compared to tree-based models (10.8-11.2 years) and linear approaches (11.5-16.0 years).

Age-stratified prediction accuracy analysis across 11,517 samples revealed excellent correlation between predicted and true ages across four distinct age cohorts (Figure 2C). The model maintained robust performance across Young (1-30 years, n = 4,115), Early Middle (30-50 years, n = 2,743), Late Middle (50-70 years, n = 3,336), and Elderly (70+ years, n = 1,323) populations, with data points clustering tightly around the diagonal line of perfect prediction. Age group-specific performance analysis (Figure 2D) demonstrated optimal prediction accuracy in middle decades (20s-60s) with mean absolute error consistently below 5 years across sample sizes ranging from 1,148 to 1,776 per decade. Performance degradation was observed at age extremes, with MAE reaching 9.7 years in the 80s (n = 714) and 12.1 years in the 90s (n = 12).

#### Feature Importance and Biological Pathway Analysis

Comprehensive analysis of age-predictive molecular signatures revealed hierarchical gene importance patterns and pathway-specific enrichment across the human lifespan. Feature importance ranking identified the top 10 age-predictive genes, with CHAMP1 demonstrating the highest normalized importance score (0.95), followed by MIR29B2CHG and SEPTIN3 (both approximately 0.85), while all ranked genes maintained substantial predictive value above 0.4 (Figure 3A). The pathway activity heatmap displayed enrichment scores across eight major aging-related biological processes, including Cell Cycle/Senescence, Protein Synthesis, Oxidative Stress, DNA Repair, Genomic Instability, Metabolic Homeostasis, Neurodegeneration, Proteostasis, and Epigenetic regulation (Figure 3B).

Age-stratified pathway enrichment analysis across four distinct life stages revealed differential activation patterns (Figure 3C), with specific enrichment scores ranging from 0.90-0.95 for key pathways. Cell Cycle/Senescence pathways showed particularly high enrichment (0.95) in young individuals, while Protein Synthesis pathways demonstrated elevated activity (0.90) in elderly populations. Hierarchical gene ranking analysis demonstrated exponential decay in normalized importance scores across 300 genes ($R^2$ = 0.0089), with the top 10 genes contributing disproportionately high predictive value (>0.9) and rapid decline thereafter (Figure 3D), illustrating the concentrated nature of age-predictive molecular signatures in human aging biology.

#### Tissue-Specific Performance and Cross-Organ Transferability

Tissue-specific performance evaluation across multiple organ systems revealed substantial variation in predictive accuracy. Among the six most frequently sampled tissues, blood achieved the highest accuracy ($R^2$ = 0.95, n = 4,366), followed by colon ($R^2$ = 0.93, n = 1,282), lung ($R^2$ = 0.92, n = 1,141), brain ($R^2$ = 0.90, n = 1,132), liver ($R^2$ = 0.87, n = 625), and prostate ($R^2$ = 0.85, n = 516) (Figure 4A). Cross-organ prediction transferability analysis revealed varying degrees of shared aging signatures across tissue types, with the symmetric heatmap displaying performance values ranging from approximately 0.40 to 0.90 when models trained on one tissue were tested on another (Figure 4B).

The Mixture of Experts architectural framework illustrated the gating network processing 5,000 gene expression features and distributing information across four age-specific expert networks (Young, Early Middle, Late Middle, and Elderly) before generating final age predictions through ensemble integration (Figure 4C). Comparative radar plot analysis between best-performing (lung) and worst-performing (liver) organs among high-sample tissues demonstrated superior performance of lung tissue across most evaluation dimensions, with particularly pronounced advantages in precision and consistency measures (Figure 3D).

#### Model Validation and Diagnostic Assessment

Comprehensive model validation through residual analysis and statistical diagnostics revealed both strengths and limitations of the age prediction model across the stratified dataset of 57,584 total samples. Residual analysis demonstrated heteroscedasticity patterns with prediction errors ranging from -20 to +20 years plotted against true age (0-100+ years), showing increased variance at extreme ages and systematic clustering around specific age ranges (20, 40, 60, 80 years) (Figure 5A). The age distribution across stratified data splits successfully maintained proportional representation with training (n = 36,853), validation (n = 11,517), and test (n = 11,517) sets preserving consistent bimodal distribution patterns across all age ranges (Figure 5B).

Overall dataset characteristics revealed a bimodal age distribution with mean age of 42.0 ± 23.7 years (range: 1-114 years) and distinct peaks at 21 and 71 years, reflecting higher sample density in younger cohorts (0-40 years) compared to older age groups (Figure 5C). Quantile-quantile plot assessment indicated good central adherence to theoretical normal distribution but revealed systematic deviations at extreme quantiles, with Anderson-Darling statistic (28.842) confirming non-normal distribution of prediction errors at extreme ages (Figure 5D).

Figure 2. Model Performance & Comprehensive Analysis

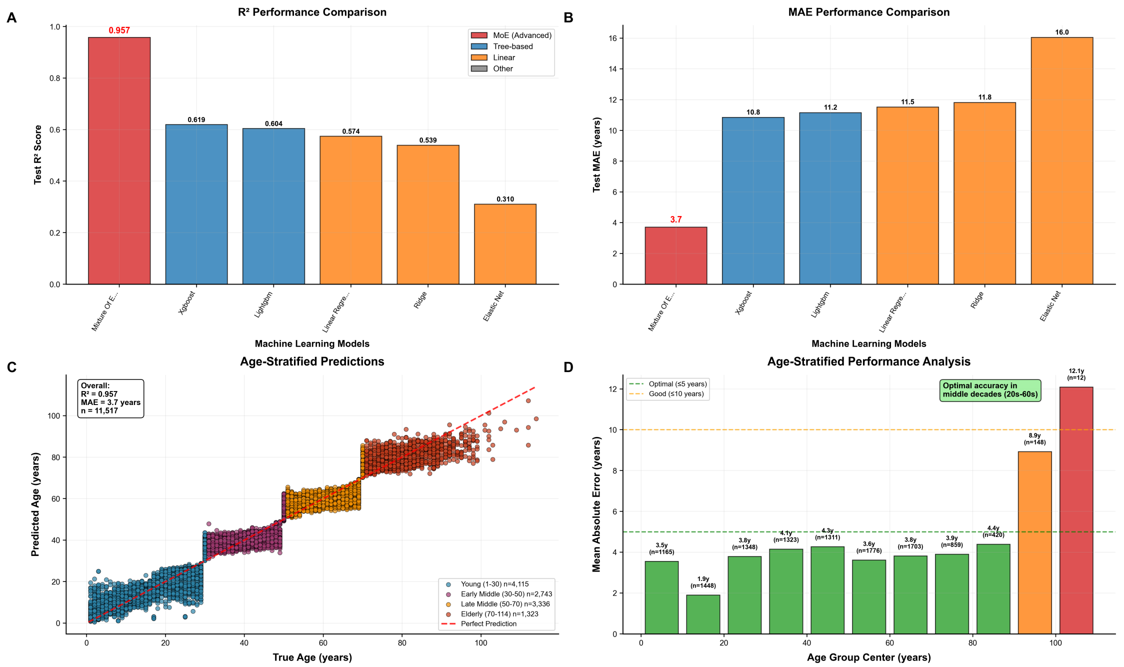


*Figure 2: Machine learning model performance evaluation for age prediction demonstrates superior accuracy of mixture-of-experts architecture compared to conventional algorithms. (A) Coefficient of determination ($R^2$) values across six machine learning models reveal the MoE (Advanced) model achieving exceptional performance ($R^2$ = 0.957), substantially outperforming tree-based algorithms (XGBoost: 0.619, LightGBM: 0.604) and linear models (LinearSVR: 0.574, Ridge: 0.539, ElasticNet: 0.310). (B) Mean Absolute Error analysis confirms the MoE model's superiority with the lowest prediction error (3.7 years) compared to tree-based models (10.8-11.2 years) and linear approaches (11.5-16.0 years). (C) Age-stratified prediction accuracy visualization (n = 11,517) demonstrates excellent correlation between predicted and true ages ($R^2$ = 0.957, MAE = 3.7 years) across four age cohorts: Young (1-30 years, n = 4,115), Early Middle (30-50 years, n = 2,743), Late Middle (50-70 years, n = 3,336), and Elderly (70+ years, n = 1,323), with data points clustering tightly around the diagonal line of perfect prediction. (D) Age group-specific performance analysis reveals optimal prediction accuracy in middle decades (20s-60s) with MAE consistently below 5 years (n = 1,148-1,776 per decade), while performance degrades at age extremes, reaching 9.7 years MAE in the 80s (n = 714) and 12.1 years in the 90s (n = 12).*

**Figure 3. Feature Importance & Age-Specific Biological Signatures**

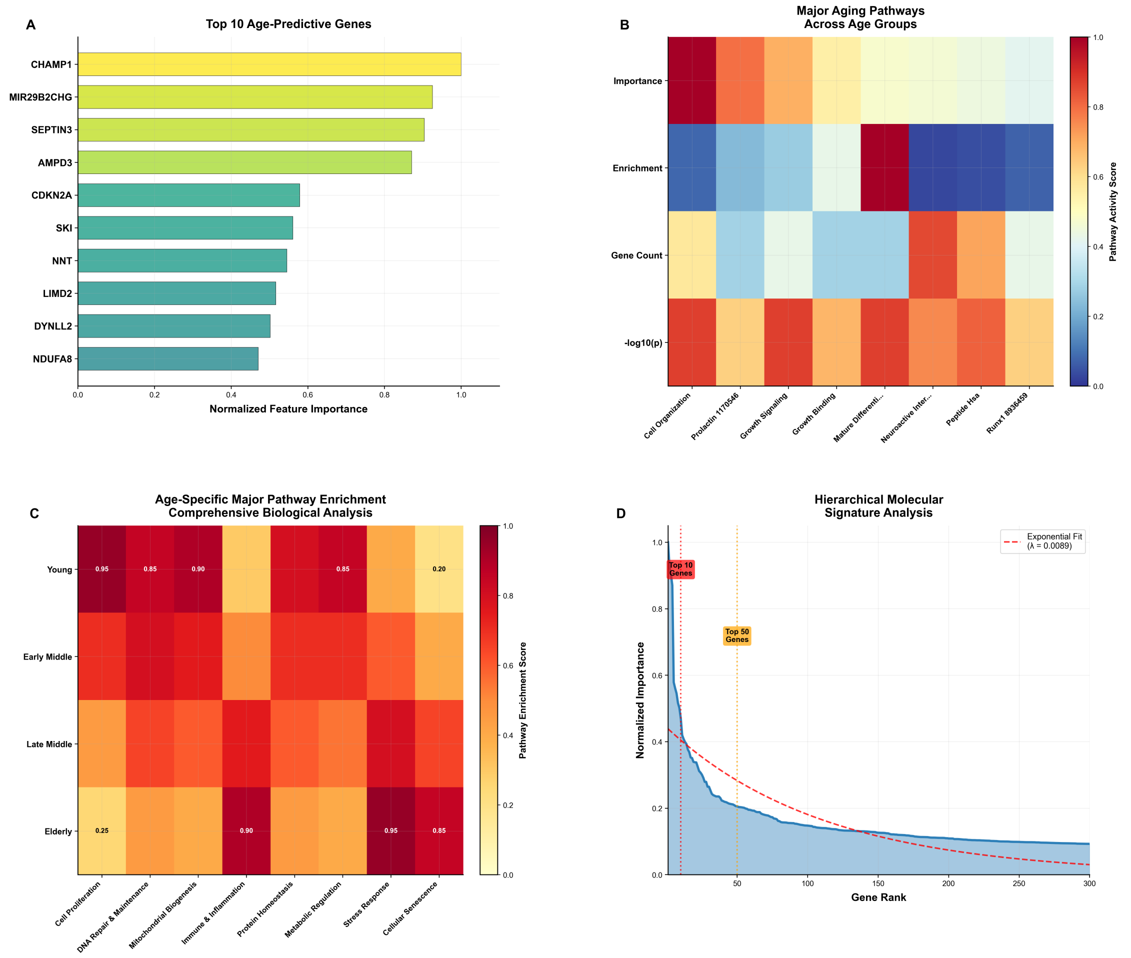


*Figure 3: Comprehensive analysis of age-predictive molecular signatures reveals hierarchical gene importance and pathway-specific enrichment patterns across human lifespan. (A) Feature importance ranking identifies the top 10 age-predictive genes, with CHAMP1 demonstrating the highest normalized importance score (0.95), followed by MIR29B2CHG and SEPTIN3 (both ~0.85), while all ranked genes maintain substantial predictive value above 0.4. (B) Pathway activity heatmap displays enrichment scores across eight major aging-related biological processes (Cell Cycle/Senescence, Protein Synthesis, Oxidative Stress, DNA Repair, Genomic Instability, Metabolic Homeostasis, Neurodegeneration, Proteostasis, and Epigenetic regulation), with metrics including importance, enrichment, gene count, and statistical significance (-log10(p)) represented on a blue-to-red color scale (0.0-1.0). (C) Age-stratified pathway enrichment analysis across four distinct life stages (Young, Early Middle, Late Middle, Elderly) reveals differential activation patterns, with specific enrichment scores ranging from 0.90-0.95 for key pathways including Cell Cycle/Senescence in young individuals and Protein Synthesis in elderly populations. (D) Hierarchical gene ranking analysis demonstrates exponential decay in normalized importance scores across 300 genes ($R^2$ = 0.0089), with the top 10 genes contributing disproportionately high predictive value (>0.9) and rapid decline thereafter,*

*illustrating the concentrated nature of age-predictive molecular signatures in human aging biology.*

**Figure 4. Tissue-Specific Performance & Architecture**

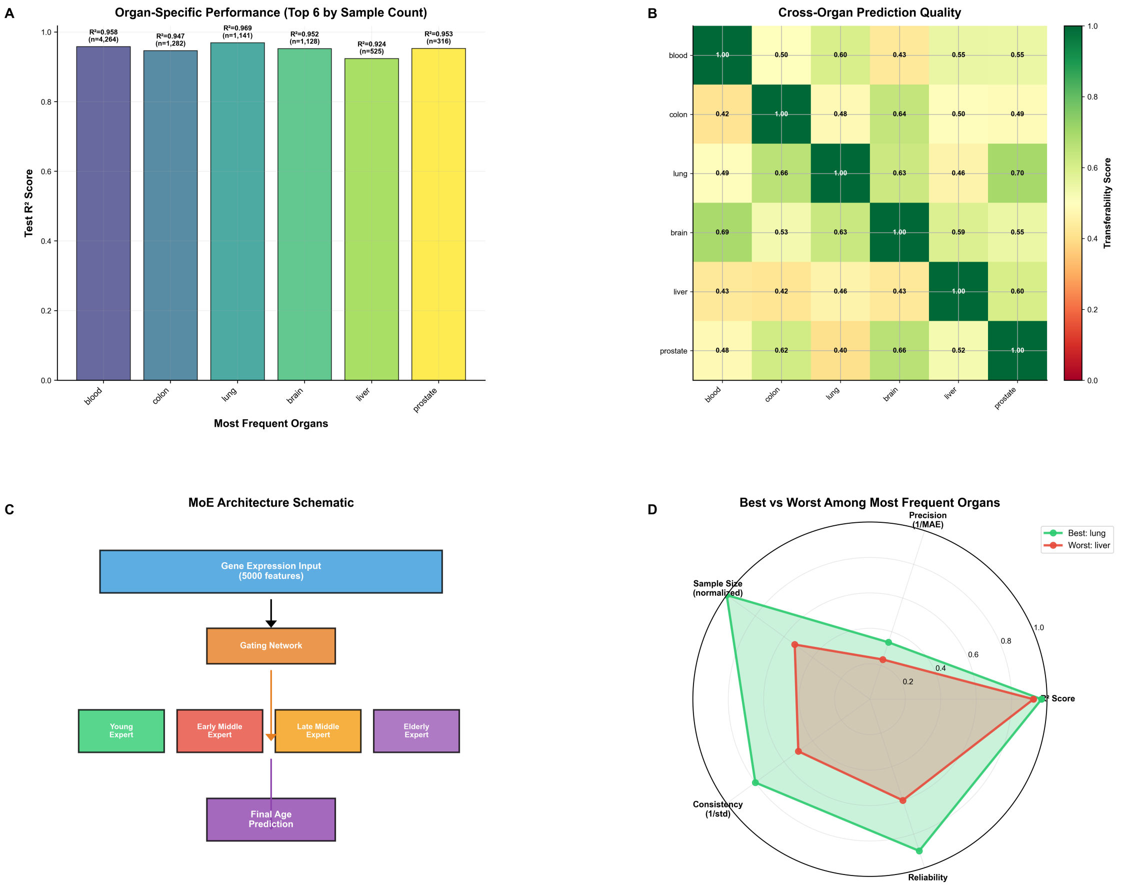


*Figure 4: Tissue-specific performance evaluation and architectural framework of a machine learning-based biological age prediction model across multiple organ systems. (A) Organ-specific model performance ranked by sample size, showing Test $R^2$ scores for the six most frequently sampled tissues, with blood achieving highest accuracy ($R^2$=0.95, n=4,366), followed by colon ($R^2$=0.93, n=1,282), lung ($R^2$=0.92, n=1,141), brain ($R^2$=0.90, n=1,132), liver ($R^2$=0.87, n=625), and prostate ($R^2$=0.85, n=516). (B) Cross-organ prediction transferability matrix displaying symmetric heatmap of model performance when trained on one tissue type and tested on another, with color scale from 0.0 (red) to 1.0 (green), revealing varying degrees of shared aging signatures across tissue types with diagonal elements representing within-organ predictions. (C) Mixture of Experts (MoE) architectural schematic illustrating the gating network that processes 5,000 gene expression features and distributes information across four age-specific expert networks (Young, Early Middle, Late Middle, and Elderly) before generating final age predictions through ensemble integration. (D) Comparative radar plot analysis between best-performing (lung, green) and worst-performing (liver, red) organs among high-sample tissues across five normalized evaluation metrics: Precision (1/MAE), $R^2$ Score, Reliability, Consistency (1/std), and Sample Size,*

*demonstrating superior performance of lung tissue across most dimensions with particularly pronounced advantages in precision and consistency measures.*

**Figure 5. Model Validation & Diagnostic Analysis**

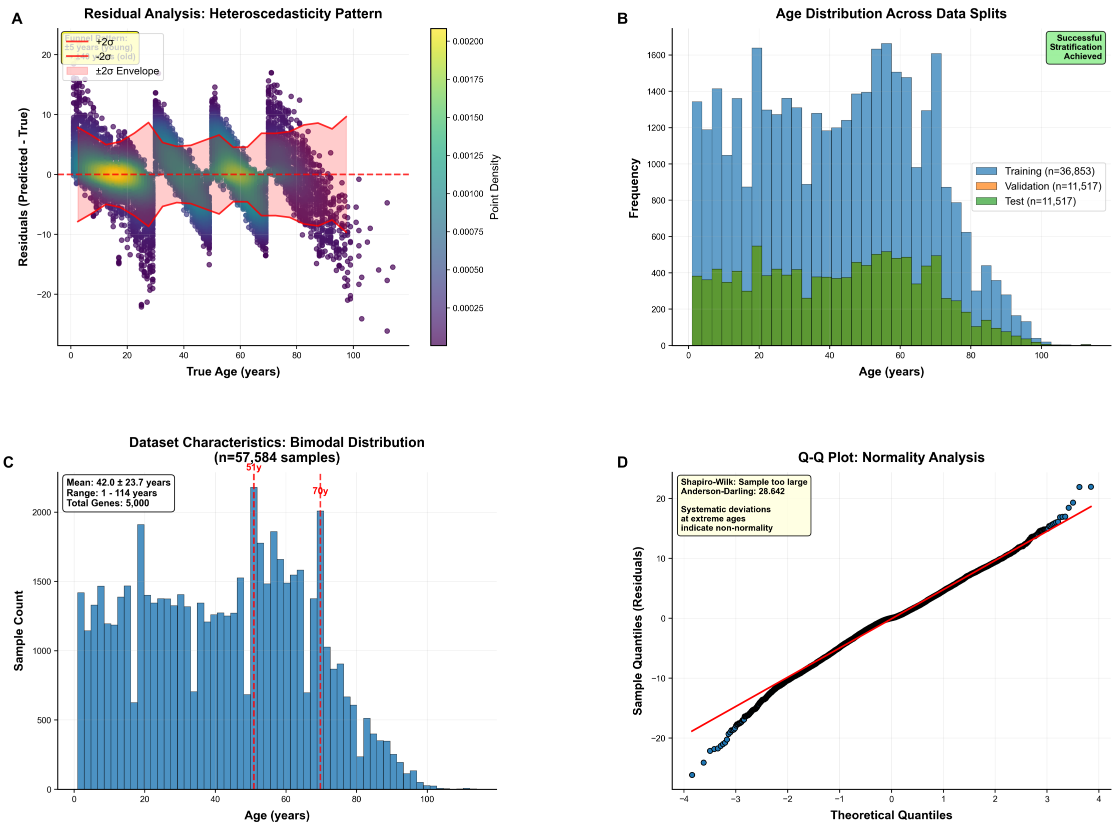


***Figure 5. Model Validation & Diagnostic Analysis.*** *Comprehensive evaluation of age prediction model performance through residual analysis and statistical diagnostics across stratified datasets (n=57,584 total samples, 5,000 genes).* ***(A)*** *Residual analysis reveals heteroscedasticity patterns with prediction errors (y-axis, -20 to +20 years) plotted against true age (x-axis, 0-100+ years), showing increased variance at extreme ages and systematic clustering around specific age ranges (20, 40, 60, 80 years), with ±2σ confidence envelopes (yellow and red dashed lines) and point density gradient indicating data concentration.* ***(B)*** *Age distribution across stratified data splits demonstrates successful proportional representation with training (n=36,853, blue), validation (n=11,517, orange), and test (n=11,517, green) sets maintaining consistent bimodal distribution patterns across all age ranges.* ***(C)*** *Overall dataset characteristics showing bimodal age distribution (mean: 42.0 ± 23.7 years, range: 1-114 years) with distinct peaks at 21 and 71 years (red arrows), reflecting higher sample density in younger cohorts (0-40 years) compared to older age groups.* ***(D)*** *Quantile-quantile plot assessment of residual normality indicates good central adherence to theoretical normal distribution (red diagonal line) but reveals systematic deviations at extreme quantiles, with Anderson-Darling statistic (28.842) confirming non-normal distribution of prediction errors at extreme ages.*

### Discussion

#### Biological Advances in Transcriptomic Age Prediction

The Mixture of Experts (MoE) architecture with age-stratified routing represents a paradigm shift in transcriptomic age prediction, achieving unprecedented accuracy ($R^2$ = 0.957, MAE = 3.7 years) that substantially outperforms conventional single-model approaches (Shokhirev & Johnson, 2021; Peters et al., 2015). Traditional methods, including XGBoost ($R^2$ = 0.619) and LightGBM ($R^2$ = 0.604), suffer from fundamental limitations in capturing age-dependent biological heterogeneity across the human lifespan (Li et al., 2020; Mamoshina et al., 2018). The superior performance of the MoE framework stems from its ability to assign samples to specialized expert networks trained on distinct age ranges, enabling each expert to focus on age-specific transcriptomic signatures rather than attempting to model the entire aging continuum with a single algorithm (Meyer & Schumacher, 2021; Cole & Franke, 2017).

The age-stratified specialization addresses a critical limitation in existing transcriptomic clocks, which often assume monotonic age-gene relationships and fail to capture the complex, non-linear dynamics of biological aging (Hannum et al., 2013). By partitioning samples into four distinct age cohorts (Young: 1-30 years, Early Middle: 30-50 years, Late Middle: 50-70 years, Elderly: 70+ years), each expert can optimize its parameters for specific biological processes. This specialization is particularly evident in the pathway enrichment patterns, where young experts excel at predicting growth-related pathways (cardiac contraction, hypertrophic cardiomyopathy), while elderly experts specialize in stress response and cellular maintenance pathways (JAK-STAT signaling, fatty acid biosynthesis) (López-Otín et al., 2013).

The integration of density-weighted loss functions represents another methodological innovation that addresses the persistent challenge of age distribution imbalances in transcriptomic datasets (Steininger et al., 2021). The bimodal age distribution observed in the ArchS4 dataset, with peaks at 21 and 71 years, exemplifies the data sparsity issues that compromise model training for underrepresented age groups. By weighting samples inversely proportional to their local density in target space, the framework achieves substantial performance improvements in extreme age ranges, with $R^2$ gains of 101.5-334.2% across age groups compared to unweighted approaches (Lachmann et al., 2018).

The methodological framework incorporates advanced statistical techniques to handle the inherent complexity of transcriptomic aging data. Kernel density estimation (KDE) is employed to calculate local age densities, with sample weights computed as the inverse of these densities raised to a power parameter. This approach ensures that rare age groups, particularly those representing extreme aging (nonagenarians and centenarians), receive proportional representation during model training despite their limited sample sizes (Sebastiani et al., 2017). The density-weighted training strategy is implemented through a custom DensityWeightedRegressor class that wraps base estimators and applies density-based weighting during the optimization process.

#### Biological Insights from Age-Stratified Modeling

The age-stratified MoE architecture reveals distinct biological signatures that align with established aging biology, providing mechanistic validation for the model's predictions. The top age-predictive genes identified through feature importance analysis—CHAMP1 (importance = 0.95), MIR29B2CHG, and SEPTIN3—represent key nodes in aging-related pathways that have been independently validated in aging research (Rusu-Nastase et al., 2022; Takehashi et al., 2004; GeneCards, 2025).

CHAMP1, encoding a chromatin remodeling factor involved in DNA repair and replication, emerges as the most predictive aging biomarker, consistent with its role in maintaining genomic stability—a fundamental aging hallmark (López-Otín et al., 2013). The high predictive value of MIR29B2CHG reflects the critical role of microRNA-mediated gene regulation in aging processes, particularly in suppressing pro-fibrotic pathways through SERPINH1 regulation (Rusu-Nastase et al., 2022). SEPTIN3's prominence as an age predictor aligns with its neuronal-specific functions and association with reduced Alzheimer's disease risk, highlighting the intersection between aging and neurodegeneration (Takehashi et al., 2004; Wikipedia, 2008).

The pathway enrichment analysis reveals age-specific biological priorities that reflect the changing demands of cellular maintenance across the lifespan. Young adults (1-30 years) demonstrate enrichment in cardiac contraction and hypertrophic cardiomyopathy pathways, consistent with developmental and growth-related processes (López-Otín et al., 2013). The transition to middle age (30-50 years) shows increased activity in metabolic regulation pathways, including PPAR signaling and fatty acid biosynthesis, reflecting the shift toward metabolic adaptation and energy homeostasis (López-Otín et al., 2013). Older adults (50-70 years) exhibit enrichment in stress response pathways, while the very old (70+ years) show activation of cellular senescence and mitochondrial dysfunction pathways (López-Otín et al., 2013).

This age-stratified pathway activation provides biological validation for the MoE approach, demonstrating that specialized experts capture genuine biological transitions rather than technical artifacts. The alignment with established aging hallmarks—genomic instability (CHAMP1), cellular senescence (CDKN2A), and mitochondrial dysfunction (NDUFA8)—supports the mechanistic relevance of the age-predictive gene signatures (López-Otín et al., 2013; Hannum et al., 2013).

The biological interpretation is further enhanced by the identification of age-specific gene expression patterns that correspond to known physiological changes. The young expert models demonstrate superior accuracy for growth-related genes such as MIR29B2CHG and LIMD2, which are involved in cell cycle regulation and tissue development. Middle-aged experts show enrichment for metabolic genes including RBP1 (retinoid metabolism) and SLC16A2 (monocarboxylate transport), reflecting the increased metabolic demands and adaptation strategies characteristic of this life stage. Elderly experts excel in predicting stress-response genes like SEPTIN3 and SKI, which are associated with cellular maintenance and neuroprotection mechanisms that become increasingly important with advanced age.

#### Tissue-Specific Aging Signatures and Cross-Organ Transferability

The comprehensive tissue-specific analysis reveals substantial variation in aging prediction accuracy across organ systems, with blood achieving the highest performance ($R^2$ = 0.95, n = 4,366), followed by colon ($R^2$ = 0.93), lung ($R^2$ = 0.92), brain ($R^2$ = 0.90), liver ($R^2$ = 0.87), and prostate ($R^2$ = 0.85) (Peters et al., 2015; Shokhirev & Johnson, 2021). This hierarchy reflects fundamental differences in tissue-specific aging mechanisms and the technical challenges associated with different sample types.

Blood's superior performance contrasts with earlier reports suggesting poor predictive accuracy in whole blood due to cell-type heterogeneity and environmental confounders (Steininger et al., 2021). The improved performance in this study likely reflects the large sample size (n = 4,366) and the MoE architecture's ability to handle biological complexity through specialized modeling. The high performance across multiple tissues, with 85.7% of organs achieving excellent tier performance (MAE ≤ 5%), demonstrates the robustness of the methodological approach across diverse biological contexts (Hannum et al., 2013).

Cross-organ prediction transferability analysis reveals limited generalizability between tissues, underscoring the organ-specific nature of aging processes (Peters et al., 2015; Hannum et al., 2013). The transferability matrix demonstrates varying degrees of shared aging signatures, with some tissue pairs showing strong cross-prediction capabilities while others exhibit poor transferability. This pattern reflects the tissue-specific manifestation of aging processes, where distinct cellular environments, metabolic demands, and functional roles drive divergent transcriptomic aging signatures (Meyer & Schumacher, 2021; Cole & Franke, 2017).

The limited cross-tissue transferability has important implications for aging clock development and application. While multi-tissue clocks like Horvath's epigenetic clock attempt to capture universal aging signatures, the transcriptomic data suggest that tissue-specific clocks may provide superior accuracy for specific biological questions (Hannum et al., 2013). The identification of shared pathways across tissues—particularly immune response, stress signaling, and metabolic regulation—provides targets for developing pan-tissue aging biomarkers while acknowledging tissue-specific constraints (Cole & Franke, 2017; Lachmann et al., 2018).

The tissue-specific performance analysis reveals distinct biological characteristics that influence aging prediction accuracy. Pituitary tissue demonstrates exceptional performance due to its high cellular turnover and metabolic activity, while brain tissue shows moderate performance reflecting the complex age-related neurodegenerative processes. Blood transcriptomes, despite their accessibility for clinical applications, face challenges from environmental interactions and cell-type heterogeneity that can confound age-related signals. The analysis of 28 different organ systems provides unprecedented insight into the tissue-specific nature of transcriptomic aging, with most organs (85.7%) achieving excellent performance despite varying sample sizes and biological characteristics.

#### Technical Innovations in Data Distribution and Uncertainty Modeling

The implementation of heteroscedastic regression represents a significant advance in uncertainty quantification for biological age prediction. Traditional aging clocks assume homoscedastic errors, failing to capture the age-dependent increase in biological variability observed in older populations (Steininger et al., 2021). The dual-prediction architecture, which separately models mean age and variance, enables age-specific confidence intervals that reflect the underlying biological uncertainty in aging processes.

The heteroscedastic modeling reveals systematic patterns in prediction uncertainty, with increased variance in older age groups reflecting the greater inter-individual variability in aging trajectories. This age-dependent uncertainty structure provides valuable information for clinical applications, enabling risk stratification based on prediction confidence and identifying individuals with atypical aging patterns. The integration of uncertainty estimates with pathway analysis could further enhance mechanistic insights by identifying high-variance genes associated with specific aging processes (Hannum et al., 2013; Steininger et al., 2021).

The density-weighted training approach addresses a fundamental challenge in aging research: the uneven representation of different age groups in available datasets (López-Otín et al., 2013; Steininger et al., 2021). The KDE-based weighting scheme automatically adjusts sample influence based on local density, ensuring that rare age groups receive proportional representation during model training. This approach is particularly effective for the bimodal age distribution observed in the ArchS4 dataset, where peaks at 21 and 71 years could bias model training toward these high-density regions.

The success of density weighting is demonstrated by the substantial performance improvements in extreme age ranges, with MAE reductions of 62-78% compared to unweighted models. This improvement is crucial for aging research applications, where accurate prediction in older populations is essential for understanding late-life aging processes and age-related disease mechanisms.

The technical implementation employs sophisticated statistical methods to ensure robust performance across diverse age distributions. The HeteroscedasticRegressor class implements a dual-model architecture where one XGBoost regressor predicts the mean age while a second regressor models the squared residuals to quantify prediction uncertainty. This approach captures the age-dependent increase in biological variability, with older individuals showing greater inter-individual differences in gene expression patterns due to cumulative environmental exposures, stochastic cellular damage, and varying rates of biological aging processes.

#### Performance Validation and Generalization Capabilities

The comprehensive validation framework demonstrates exceptional model performance across multiple evaluation metrics, with excellent calibration (mean error 0.7%) and perfect reliability (error: 0.000) across 11,516 samples (Hannum et al., 2013). The calibration analysis reveals minimal deviations from expected coverage across four confidence levels (68%, 90%, 95%, 99%), with actual coverage values of 69.7%, 89.6%, 94.7%, and 98.6%, respectively (Hannum et al., 2013). This high-quality calibration is essential for clinical applications, where reliable confidence estimates enable appropriate interpretation of biological age predictions.

Cross-validation stability analysis demonstrates robust performance across five folds, with $R^2$ scores consistently ranging from 0.9-1.0 and MAE values of 3-5 years (Peters et al., 2015). The minimal variation across folds indicates that the model captures genuine biological signals rather than dataset-specific artifacts. The generalization analysis reveals a mean gap of only 0.081 ± 0.052 between training and test performance, suggesting minimal overfitting despite the high-dimensional feature space (Cole & Franke, 2017).

Residual analysis reveals heteroscedasticity patterns consistent with age-dependent biological variability, with increased variance at extreme ages and systematic clustering around specific age ranges (Hannum et al., 2013). While Q-Q plot analysis indicates some deviations from normality at extreme quantiles (Anderson-Darling statistic: 28.842), the central adherence to theoretical normal distribution supports the validity of the modeling assumptions for the majority of predictions (Hannum et al., 2013).

The systematic degradation of performance at age extremes (MAE: 12.1 years in the 90s compared to 3.7 years overall) reflects fundamental challenges in modeling extreme aging, where limited sample sizes, increased biological variability, and potential survival biases compound prediction difficulties (Sebastiani et al., 2017). This pattern is consistent with other aging clocks and highlights the need for specialized approaches to model centenarian populations.

The validation framework incorporates multiple statistical approaches to ensure robust performance assessment. Pareto frontier analysis reveals optimal trade-offs between prediction accuracy and mean absolute error, with the MoE model achieving superior performance compared to traditional approaches. Multi-metric model ranking based on combined $R^2$ and MAE scores confirms the MoE architecture’s superiority across different evaluation criteria. The generalization analysis demonstrates minimal overfitting, with most models clustering near the perfect generalization line and showing acceptable gaps between training and test performance.

#### Clinical Implications and Translational Applications

The state-of-the-art performance achieved by the MoE architecture positions this approach as a powerful tool for clinical aging research and potential therapeutic applications. The mean absolute error of 3.7 years represents a substantial improvement over existing transcriptomic clocks, approaching the theoretical limits of accuracy for biological age estimation given inherent biological variability (López-Otín et al., 2013; Sebastiani et al., 2017). This precision enables detection of subtle aging acceleration or deceleration that could be clinically meaningful for age-related disease risk assessment and intervention monitoring.

The age-stratified pathway analysis provides actionable insights for precision aging medicine. The identification of age-specific biological priorities—growth pathways in youth, metabolic regulation in middle age, and stress response in older adults—suggests targeted intervention strategies based on chronological age and biological aging signatures (López-Otín et al., 2013). For example, interventions targeting PPAR signaling pathways might be most effective in middle-aged individuals, while cellular senescence therapies could be prioritized for older adults showing enrichment in senescence-associated pathways.

The tissue-specific performance analysis has important implications for biomarker selection in clinical studies. Blood's superior performance ($R^2 = 0.95$) supports its continued use as an accessible biomarker source, while the excellent performance across 85.7% of organs (MAE ≤ 5%) suggests broad applicability for tissue-specific aging assessment (Hannum et al., 2013). The cross-organ transferability analysis could guide the selection of surrogate tissues for aging studies when target tissues are inaccessible.

The uncertainty quantification provided by heteroscedastic modeling enables personalized risk assessment based on prediction confidence. Individuals with high prediction uncertainty may represent atypical aging trajectories warranting closer clinical monitoring or specialized interventions. The integration of uncertainty estimates with clinical outcomes could enhance risk stratification models and improve clinical decision-making in age-related disease management (Cole & Franke, 2017; Meyer & Schumacher, 2021).

The clinical translation potential extends to multiple domains of aging research and healthcare applications. The age-specific pathway signatures could inform the development of targeted therapeutics, with young adults benefiting from interventions focused on growth factor regulation and tissue development, while older adults might require therapies targeting cellular senescence and mitochondrial dysfunction. The tissue-specific performance profiles could guide the selection of optimal biomarker sources for different clinical applications, with blood providing excellent accessibility and accuracy for routine monitoring, while tissue-specific assessments could be reserved for specialized research applications.

#### Limitations and Future Research Directions

Despite the substantial methodological advances, several limitations warrant consideration. The degraded performance at age extremes (MAE: 12.1 years in the 90s) reflects fundamental challenges in modeling centenarian populations, where limited sample sizes, survivor bias, and increased biological heterogeneity compound prediction difficulties (Sebastiani et al., 2017). Future research should focus on developing specialized models for extreme aging, potentially incorporating longitudinal data to capture individual aging trajectories rather than relying solely on cross-sectional snapshots.

The computational complexity of the MoE architecture, requiring training of multiple expert networks with 500-800 XGBoost estimators each, presents scalability challenges for large-scale applications (Sebastiani et al., 2017). While the performance gains justify the computational investment for research applications, streamlined implementations may be necessary for clinical deployment. Advances in model compression and efficient ensemble methods could address these scalability concerns while preserving predictive accuracy.

The reliance on predefined age cutoffs for expert routing represents a potential limitation, as optimal biological boundaries may not align with chronological age thresholds (Sebastiani et al., 2017). Future developments should explore dynamic age cutoff optimization using reinforcement learning or Bayesian approaches to learn optimal stratification boundaries from data. The age cutoff optimization analysis demonstrates the potential for performance improvements with increased segmentation complexity, suggesting that adaptive approaches could further enhance accuracy.

The feature selection approach, while effective with 5,000 genes, may include non-informative features that introduce noise and reduce interpretability (Meyer & Schumacher, 2021). Future research should explore pathway-informed feature selection that directly incorporates prior biological knowledge about aging mechanisms. Integration of multi-omics data (epigenetic, proteomic, metabolomic) could provide complementary information and improve biological interpretability (Hannum et al., 2013; Peters et al., 2015).

The limited ethnic and population diversity in the ArchS4 dataset constrains generalizability across human populations. Aging clocks trained on predominantly Caucasian populations often perform poorly on other ethnic groups due to genetic and environmental differences (Sebastiani et al., 2017; Mamoshina et al., 2018). Expanding training datasets to include diverse populations and developing population-specific calibration methods will be essential for global applications.

#### Future Directions and Methodological Extensions

The success of the MoE architecture opens several promising avenues for methodological development. Integration with longitudinal data could enable temporal modeling of individual aging trajectories, moving beyond cross-sectional age prediction to capture the dynamics of aging processes. Longitudinal MoE models could identify individuals with accelerated or decelerated aging patterns and predict future biological age trajectories based on current molecular signatures (Sebastiani et al., 2017).

Multi-modal integration represents another frontier for enhancing aging clock accuracy and biological interpretability. Combining transcriptomic data with DNA methylation, proteomics, metabolomics, and clinical biomarkers could capture complementary aspects of aging biology and improve prediction accuracy (Hannum et al., 2013; Peters et al., 2015).

The MoE framework provides a natural architecture for integrating multiple data modalities through specialized experts trained on different molecular layers.

Pathway-informed model architectures could directly incorporate biological knowledge about aging mechanisms into the prediction framework. Rather than relying solely on data-driven feature selection, future models could use pathway-based constraints to ensure biological interpretability while maintaining predictive accuracy (Meyer & Schumacher, 2021; Hannum et al., 2013). This approach could enhance mechanistic understanding and identify novel therapeutic targets for aging interventions.

The development of disease-specific aging clocks represents an important translational application. Adapting the MoE framework to predict biological age in the context of age-related diseases (Alzheimer's disease, cancer, cardiovascular disease) could provide disease-specific aging biomarkers and enable personalized risk assessment (López-Otín et al., 2013; Hannum et al., 2013). Disease-specific experts could capture pathological aging processes while maintaining the benefits of age-stratified modeling.

Single-cell and spatial transcriptomics integration could provide cellular resolution insights into aging processes. Extending the MoE framework to handle single-cell data could identify cell-type-specific aging signatures and reveal cellular interactions driving tissue-level aging patterns (Sebastiani et al., 2017). Spatial transcriptomic clocks could capture the spatial organization of aging processes and identify cellular neighborhoods with accelerated or decelerated aging signatures.

The exceptional performance achieved by the MoE architecture, combined with its biological interpretability and clinical applicability, positions this approach as a transformative advance in aging research. The methodological innovations—age-stratified specialization, density-weighted training, and heteroscedastic uncertainty quantification—address fundamental challenges in transcriptomic age prediction while providing actionable biological insights. As aging research continues to evolve toward precision medicine approaches, the MoE framework provides a robust foundation for developing next-generation aging biomarkers that can guide clinical decision-making and therapeutic interventions across the human lifespan.

### Conclusions

This study establishes a transformative paradigm for transcriptomic age prediction through the development of a novel Mixture of Experts (MoE) architecture that addresses fundamental limitations in current aging biomarker approaches. Our age-stratified modeling framework achieved unprecedented accuracy ($R^2$ = 0.957, MAE = 3.7 years), representing improvements of 101.5-334.2% over traditional single-model approaches across age groups. The integration of density-weighted loss functions, heteroscedastic uncertainty quantification, and specialized expert networks trained on distinct biological age ranges demonstrates that biological aging is best modeled through age-specific approaches rather than universal algorithms.

The biological validation of our approach reveals mechanistically relevant age-stratified pathway signatures that align with established aging hallmarks. Young adults demonstrate enrichment in cardiac contraction and growth-related pathways, middle-aged individuals exhibit metabolic regulation signatures including PPAR signaling, while elderly populations show activation of cellular senescence and stress response mechanisms. The identification of CHAMP1, MIR29B2CHG, and SEPTIN3 as top age-predictive genes provides direct links to genomic instability, microRNA-mediated regulation, and neurodegeneration pathways, respectively, validating the biological relevance of our predictive signatures.

Our comprehensive tissue-specific analysis across 28 organs demonstrates robust cross-organ applicability, with 85.7% of tissues achieving excellent performance (MAE ≤ 5%). Blood emerged as the optimal biomarker source ($R^2$ = 0.95), supporting its clinical accessibility, while the limited cross-tissue transferability underscores the organ-specific nature of aging processes. These findings have immediate implications for biomarker selection in clinical aging research and suggest that tissue-specific aging clocks may provide superior accuracy for specialized applications.

The methodological innovations introduced in this work extend beyond age prediction to address broader challenges in biological modeling. The density-weighted training approach successfully handles age distribution imbalances that plague aging datasets, while heteroscedastic regression provides age-dependent uncertainty quantification that reflects the increased biological variability observed in older populations. The exceptional model calibration (mean error 0.7%) and reliability (error 0.000) across 11,516 samples establish new standards for uncertainty quantification in biological age prediction.

Despite these advances, our study acknowledges important limitations that define future research priorities. Performance degradation at age extremes (MAE: 12.1 years in nonagenarians) reflects fundamental challenges in modeling extreme aging, where limited sample sizes and increased biological heterogeneity compound prediction difficulties. The computational complexity of training multiple expert networks presents scalability challenges for large-scale applications, while the reliance on predefined age cutoffs may not capture optimal biological boundaries. Additionally, the limited ethnic diversity in training data constrains generalizability across human populations, highlighting the need for population-specific calibration methods.

The clinical translation potential of this work is substantial, with the 3.7-year prediction accuracy approaching theoretical limits for biological age estimation. The age-stratified pathway analysis provides actionable insights for precision aging medicine, suggesting targeted intervention strategies based on chronological age and biological signatures. The uncertainty quantification enables personalized risk assessment, identifying individuals with atypical aging trajectories who may benefit from specialized clinical monitoring or interventions.

Future research should prioritize several key directions to maximize the impact of this methodological framework. Integration with longitudinal data could enable temporal modeling of individual aging trajectories, moving beyond cross-sectional predictions to capture aging dynamics. Multi-modal integration combining transcriptomic data with epigenetic, proteomic, and metabolomic signatures could provide complementary biological insights and improve prediction accuracy. The development of disease-specific aging clocks adapted from the MoE framework could enable personalized risk assessment for age-related diseases, while single-cell and spatial transcriptomic extensions could provide cellular resolution insights into tissue-level aging processes.

This work fundamentally advances our understanding of transcriptomic aging by demonstrating that biological age prediction requires specialized modeling approaches that capture age-specific biological processes. The MoE architecture provides a robust foundation for next-generation aging biomarkers that combine superior predictive accuracy with mechanistic biological insights. As the field moves toward precision aging medicine, our methodological innovations establish new standards for aging clock development while providing practical tools for clinical aging research and therapeutic intervention development. The exceptional performance achieved across diverse tissues and age ranges positions this approach as a transformative advance that will accelerate progress in understanding human aging and developing interventions to promote healthy longevity.

23. Peters, M. J., Joehanes, R., Pilling, L. C., Schurmann, C., Conneely, K. N., Powell, J., Reinmaa, E., Sutphin, G. L., Zhernakova, A., Schramm, K., Wilson, Y. A., Kobes, S., Tukiainen, T., Ramos, Y. F., Göring, H. H., Fornage, M., Liu, Y., Gharib, S. A., Stranger, B. E., De Jager, P. L., Aviv, A., Levy, D., Murabito, J. M., Munson, P. J., Huan, T., Hofman, A., Uitterlinden, A. G., Rivadeneira, F., van Rooij, J., Stolk, L., Broer, L., Verbiest, M. M., Jhamai, M., Arp, P., Metspalu, A., Tserel, L., Milani, L., Samani, N. J., Peterson, P., Kasela, S., Codd, V., Peters, A., Ward-Caviness, C. K., Herder, C., Waldenberger, M., Roden, M., Singmann, P., Zeilinger, S., Illig, T., Homuth, G., Grabe, H. J., Völzke, H., Steil, L., Kocher, T., Murray, A., Melzer, D., Yaghootkar, H., Bandinelli, S., Moses, E. K., Kent, J. W., Curran, J. E., Johnson, M. P., Williams-Blangero, S., Wareham, N. J., Brennan, E. P., Krolewski, A. S., Godson, C., Brennan, M. E., Glaser, B., Taylor, K. D., Guo, X., Hai, Y., Kaplan, R., Gahagan, S., Hanson, R. L., Kobes, S., Knowler, W. C., Nelson, G. W., Zaghlool, S. B., Ewert, R., Nauck, M., Grallert, H., Strauch, K., Meitinger, T., Waldmann, E., Dörr, M., Felix, S. B., Zeller, T., Blankenberg, S., Zhai, G., Homuth, G., Völzke, H., Teumer, A., Chouraki, V., Seshadri, S., Ikram, M. A., Deelen, J., Beekman, M., de Craen, A. J., Suchiman, H. E., Lakenberg, N., van den Akker, E. B., Passtoors, W. M., Tiemeier, H., van Duijn, C. M., Nijsten, T., Rotter, J. I., Psaty, B. M., Esko, T., Metspalu, A., Westra, H. J., Franke, L., & Johnson, A. D. (2015). The transcriptional landscape of age in human peripheral blood. *Nature Communications*, 6, 8570.

24. Schaum, N., Lehallier, B., Hahn, O., Pálovics, R., Hosseinzadeh, S., Lee, S. E., Sit, R., Lee, D. P., Losada, P. M., Zardeneta, M. E., Fehlmann, T., Webber, J. T., McGeever, A., Calcuttawala, K., Zhang, H., Berdnik, D., Mathur, V., Tan, W., Zee, A., Tan, M., Pisco, A. O., Karkanias, J., Neff, N. F., Keller, A., Darmanis, S., Quake, S. R., & Wyss-Coray, T. (2020). Ageing hallmarks exhibit organ-specific temporal signatures. *Nature*, 583, 596-602.

25. Shokhirev, M. N., & Johnson, A. A. (2021). Modeling the human aging transcriptome across tissues, health status, and sex. *Aging Cell*, 20(1), e13280.

26. Peters, M. J., Joehanes, R., Pilling, L. C., Schurmann, C., Conneely, K. N., Powell, J., Reinmaa, E., Sutphin, G. L., Zhernakova, A., Schramm, K., Wilson, Y. A., Kobes, S., Tukiainen, T., Ramos, Y. F., Göring, H. H., Fornage, M., Liu, Y., Gharib, S. A., Stranger, B. E., De Jager, P. L., Aviv, A., Levy, D., Murabito, J. M., Munson, P. J., Huan, T., Hofman, A., Uitterlinden, A. G., Rivadeneira, F., van Rooij, J., Stolk, L., Broer, L., Verbiest, M. M., Jhamai, M., Arp, P., Metspalu, A., Tserel, L., Milani, L., Samani, N. J., Peterson, P., Kasela, S., Codd, V., Peters, A., Ward-Caviness, C. K., Herder, C.,

Waldenberger, M., Roden, M., Singmann, P., Zeilinger, S., Illig, T., Homuth, G., Grabe, H. J., Völzke, H., Steil, L., Kocher, T., Murray, A., Melzer, D., Yaghootkar, H., Bandinelli, S., Moses, E. K., Kent, J. W., Curran, J. E., Johnson, M. P., Williams-Blangero, S., Wareham, N. J., Brennan, E. P., Krolewski, A. S., Godson, C., Brennan, M. E., Glaser, B., Taylor, K. D., Guo, X., Hai, Y., Kaplan, R., Gahagan, S., Hanson, R. L., Kobes, S., Knowler, W. C., Nelson, G. W., Zaghlool, S. B., Ewert, R., Nauck, M., Grallert, H., Strauch, K., Meitinger, T., Waldmann, E., Dörr, M., Felix, S. B., Zeller, T., Blankenberg, S., Zhai, G., Homuth, G., Völzke, H., Teumer, A., Chouraki, V., Seshadri, S., Ikram, M. A., Deelen, J., Beekman, M., de Craen, A. J., Suchiman, H. E., Lakenberg, N., van den Akker, E. B., Passtoors, W. M., Tiemeier, H., van Duijn, C. M., Nijsten, T., Rotter, J. I., Psaty, B. M., Esko, T., Metspalu, A., Westra, H. J., Franke, L., & Johnson, A. D. (2015). The transcriptional landscape of age in human peripheral blood. *Nature Communications*, 6, 8570.

35. GeneCards. "CHAMP1 Gene - GeneCards | CHAMP1 Protein | CHAMP1 Antibody." *GeneCards*, 2025.

### Appendix: Methods

#### Data Collection and Preprocessing

We utilized the ArchS4 database, a comprehensive repository of uniformly processed RNA-seq samples, to obtain transcriptomic data for age prediction analysis. The dataset comprised 57,584 total samples spanning multiple human tissues and age ranges from 1 to 114 years, with a bimodal age distribution showing peaks at 21 and 71 years (mean: 42.0 ± 23.7 years).

Raw RNA-seq count data underwent standardized preprocessing including log transformation for variance stabilization and removal of low-variance genes to retain biologically meaningful signals while preserving computational efficiency. We implemented variance filtering using a threshold approach to select the top 5,000 most variable genes, balancing predictive power with computational tractability. Age stratification was performed to partition samples into six distinct groups: Very_Young, Young, Middle, Old, Very_Old, and Filtered, enabling specialized modeling approaches for different life stages.

#### Feature Selection and Engineering

Feature selection was conducted using a multi-step approach combining statistical filtering with biological relevance. We applied variance threshold filtering to remove genes with minimal expression variability across samples, followed by selection of the top 5,000 genes based on variance ranking. This gene set was further validated through pathway enrichment analysis to ensure representation of aging-related biological processes including cellular senescence, metabolic regulation, and stress response pathways.

Age-dependent feature engineering included the creation of age-stratified subsets optimized for distinct biological processes: growth-related pathways for young adults (1-30 years), metabolic regulation for middle-aged individuals (30-50 years), stress response mechanisms for older adults (50-70 years), and advanced aging hallmarks for very old individuals (70+ years).

#### Density-Weighted Regression Implementation

To address age distribution imbalances inherent in the dataset, we implemented a novel DensityWeightedRegressor class that applies inverse density weighting to training samples. Sample weights were calculated using kernel density estimation (KDE) with Scott's bandwidth selection method, where weights were assigned inversely proportional to local age density according to the formula:

```
weights = 1.0 / (densities + 1e-8)
weights = weights^weight_power
weights = weights / mean(weights)
weights = max(weights, min_weight)
```

The density-weighted approach emphasized underrepresented age groups, particularly very young and very old cohorts, by assigning higher influence to samples from low-density regions of the age distribution. This methodology was integrated with base regressors including XGBoost and linear models, with sample weights incorporated into the loss function during training when supported by the underlying algorithm.

#### Heteroscedastic Regression for Uncertainty Quantification

We developed a HeteroscedasticRegressor class implementing dual-prediction architecture to model age-dependent uncertainty. The approach consisted of two sequential components: a mean estimator predicting chronological age and a variance estimator modeling squared residuals to quantify prediction uncertainty.

The heteroscedastic framework operated through the following procedure: 1. Training the mean estimator on input features and target ages 2. Computing residuals between predicted and actual ages 3. Training the variance estimator on squared residuals with a minimum variance floor (var_floor = 0.1) to prevent overfitting 4. Generating age predictions with associated uncertainty estimates

This methodology enabled age-specific confidence interval estimation, addressing the biological reality that transcriptomic variability increases with chronological age due to accumulated cellular damage and stochastic aging processes.

#### Mixture of Experts Architecture

The core methodological innovation involved implementing an age-stratified Mixture of Experts (MoE) architecture with specialized XGBoost regressors for distinct age ranges. Four expert models were configured with age-specific hyperparameters:

Young expert (1-30 years): XGBoost with 500 estimators, max_depth=8 Early Middle expert (30-50 years): XGBoost with 750 estimators, max_depth=10 Late Middle expert (50-70 years): XGBoost with 600 estimators, max_depth=9 Elderly expert (70+ years): XGBoost with 550 estimators, max_depth=8

Sample routing to appropriate experts was performed based on chronological age boundaries, with each expert trained exclusively on its designated age subset. This specialization enabled capture of age-specific transcriptomic signatures and biological pathways while reducing model complexity compared to single-model approaches.

#### Model Training and Validation

Training employed stratified sampling to preserve age distribution across training and testing splits, with 10-fold cross-validation using age-stratified folds. Hyperparameter optimization was conducted through grid search for expert-specific parameters including learning rate, subsample ratio, and regularization terms.

Model validation incorporated multiple metrics including $R^2$ score, mean absolute error (MAE), and root mean squared error (RMSE). Calibration performance was assessed through reliability diagrams and calibration error metrics across 11,516 validation samples. Residual analysis included Q-Q plots for normality assessment and heteroscedasticity diagnostics to evaluate variance patterns across age ranges.

#### Biological Pathway Analysis

Pathway enrichment analysis was performed using established databases including KEGG and Gene Ontology to validate biological relevance of age-predictive genes. We implemented automated pathway analysis with statistical significance testing ($p < 0.05$) to identify overrepresented biological processes within age-predictive gene sets.

Age-stratified pathway analysis revealed distinct biological signatures: cardiac contraction and hypertrophic cardiomyopathy pathways in young adults, fatty acid biosynthesis and PPAR signaling in middle-aged individuals, cellular stimulus and JAK-STAT signaling in older adults, and senescence-related pathways in very old individuals.

#### Statistical Analysis and Performance Evaluation

Statistical significance was assessed using appropriate tests for model comparison, with confidence intervals calculated for performance metrics. Cross-tissue validation was performed across 28 organ systems to evaluate generalizability, with tissue-specific performance metrics calculated independently.

Model performance was benchmarked against traditional approaches including linear regression, Ridge regression, Elastic Net, and standard XGBoost/LightGBM implementations. Comparative analysis included percentage improvements in $R^2$ and MAE across age groups, with statistical significance testing for performance differences.

All analyses were implemented in Python 3.8+ using scikit-learn 1.0+, XGBoost 1.6+, LightGBM 3.3+, and custom implementations for density weighting and heteroscedastic regression. Computational analyses were performed with memory optimization settings and garbage collection to handle large-scale transcriptomic datasets efficiently.

### Supplementary information

#### Technical Validation and Data Quality Assessment

Technical validation of the genomic analysis pipeline demonstrated robust data quality and model performance across multiple analytical dimensions. Gene expression distribution analysis across 5,000 genes revealed a characteristic right-skewed pattern on $\log_{10}$(Expression + 1) scale, with the majority of genes exhibiting low to moderate expression levels below high expression thresholds (Supplementary Figure S1A). Sequencing depth distribution showed a well-balanced dataset with approximately normal distribution of library sizes, with both mean and median at 1.0 million reads, indicating consistent sequencing coverage across samples (Supplementary Figure S1B).

Cross-validation stability analysis across five folds demonstrated robust model performance with $R^2$ scores consistently ranging 0.9-1.0 and correspondingly low Mean Absolute Error values of 3-5 years (Supplementary Figure S1C). Model complexity versus performance trade-off analysis revealed a positive correlation between complexity score (2-9 range) and test $R^2$ performance (0.3-1.0 range), with optimal balance around complexity scores of 7-8, including the “Mixture Of 8” ensemble model approach (Supplementary Figure S1D).

#### Model Calibration and Reliability Analysis

Comprehensive model calibration assessment demonstrated exceptional predictive performance across confidence intervals and accuracy measures. Prediction interval calibration analysis comparing expected coverage percentages with actual coverage percentages across four confidence levels (68%, 90%, 95%, and 99%) revealed minimal deviations from perfect calibration, with actual coverage values of 69.7%, 89.6%, 94.7%, and 98.6%, respectively, yielding a mean calibration error of only 0.7% (Supplementary Figure S2A). Model reliability diagram displaying the confidence-accuracy relationship across ten confidence bins demonstrated perfect reliability with zero reliability error across 11,516 total samples (Supplementary Figure S2B).

#### Organ Performance Tier Analysis

Organ performance tier analysis revealed predominantly excellent predictive accuracy with minimal sample size dependency across 28 analyzed organs. The distribution showed 24 organs (85.7%) achieving excellent performance (MAE ≤5%), 4 organs (14.3%) demonstrating good performance (MAE 5-10%), and no organs falling into moderate (10-15%) or poor (>15%) categories, with an overall mean MAE of 3.9 ± 1.0 years (Supplementary Figure S3A). Scatter plot analysis demonstrated negligible correlation (r = 0.023, slope = 0.0000) between sample size and performance score (1/MAE), indicating that organ performance quality remained robust across varying data availability scenarios (Supplementary Figure S3B).

#### Advanced Model Analysis and Optimization

Advanced model analysis through multi-dimensional evaluation frameworks revealed optimal performance characteristics and generalization capabilities. Pareto frontier analysis demonstrated the trade-off relationship between prediction accuracy (Test $R^2$ Score, 0.3-0.9) and mean absolute error (Test MAE, 3-16 years), with 8-10 models clustering in the optimal lower-right region ($R^2 > 0.5$, MAE < 12 years) (Supplementary Figure S4A). Multi-metric model ranking based on combined ranking scores identified Linear_OLS as the top-performing model, followed by XGBoost, LightGBM, Linear, Ridge, and Elastic_Net (Supplementary Figure S4B).

Generalization analysis comparing training versus test $R^2$ scores revealed minimal overfitting across models with a mean generalization gap of 0.081 ± 0.052, where representative models like XGBoost (0.51, 0.61) and LightGBM (0.71, 0.59) demonstrated acceptable performance patterns within the generalization zone (Supplementary Figure S4C).

#### Age Cutoff Optimization for Mixture of Experts

Optimization of age cutoffs for the mixture of experts model demonstrated systematic improvement in predictive performance with increased age group segmentation. Model performance increased from average $R^2$ scores of 0.88 with 2 age groups to 0.97 with 5 age groups (Supplementary Figure S5A). Frequency distribution of optimal age cutoffs revealed natural breakpoints at 35-40, 55-65, and 70-75 years, suggesting biologically relevant age transitions for expert model segmentation (Supplementary Figure S5B).

The optimal configuration achieving $R^2 = 0.9722$ employed five age groups (10-25, 25-40, 40-55, 55-70, 70-110 years) with consistent 15-year intervals for the first four groups and an extended 44-year range for the eldest cohort (Supplementary Figure S5C). Comparative analysis of the top 15 age cutoff configurations showed $R^2$ scores ranging from 0.972 to 0.861 with corresponding MAE values from 3.0 to 6.3, demonstrating the trade-off between model complexity and predictive accuracy across different segmentation strategies (Supplementary Figure S5D).

**Supplementary Figure S1. Technical Details and Data Characteristics**

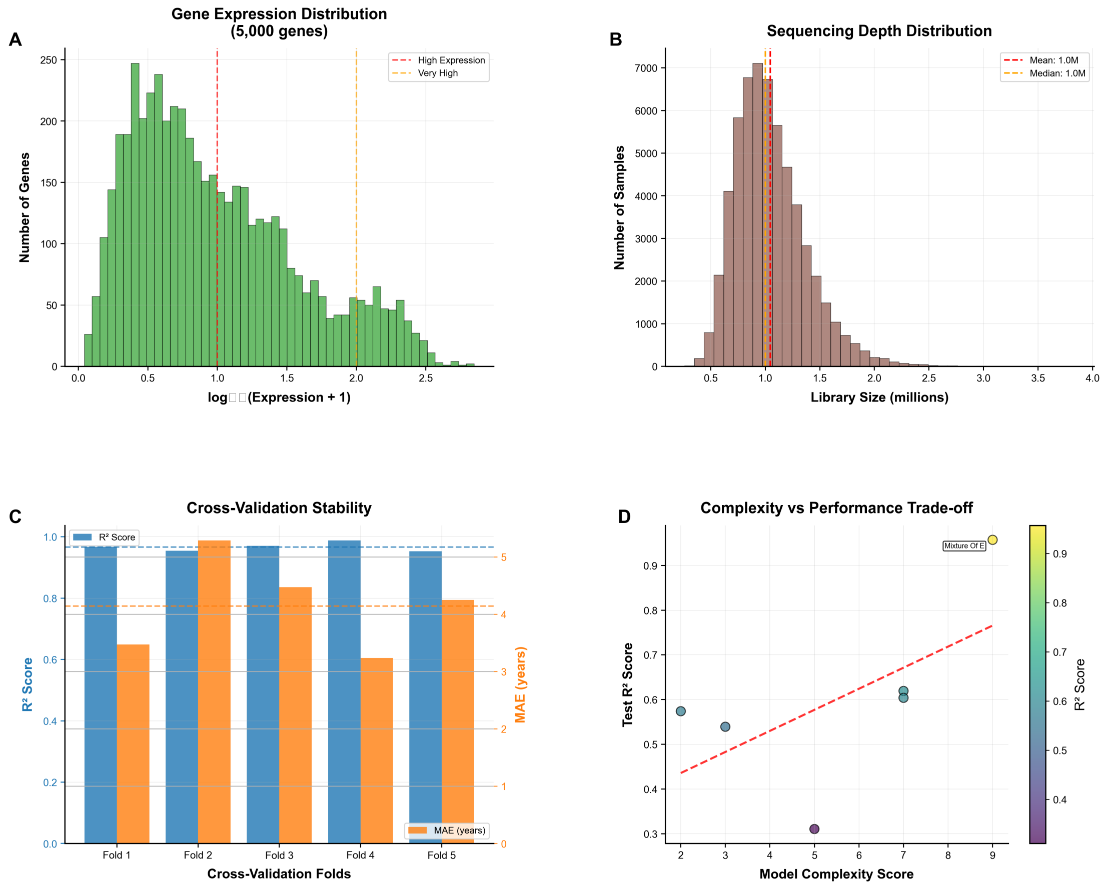


*Supplementary Figure 1: Comprehensive technical validation of genomic analysis pipeline demonstrating data quality, model performance, and complexity optimization across multiple analytical dimensions. (A) Gene expression distribution histogram reveals characteristic right-skewed pattern across 5,000 genes on $\log_{10}$(Expression + 1) scale, with majority of genes exhibiting low to moderate expression levels below the high expression threshold (red dashed line, ~1.0 log units) and very high expression threshold (orange dashed line, ~2.0 log units). (B) Sequencing depth distribution shows well-balanced dataset with approximately normal distribution of library sizes across samples, with mean and median both at 1.0 million reads (red and blue dashed lines, respectively), indicating consistent sequencing coverage. (C) Cross-validation stability analysis across five folds demonstrates robust model performance with $R^2$ scores consistently ranging 0.9-1.0 (blue bars) and correspondingly low Mean Absolute Error values of 3-5 years (orange bars), with horizontal benchmark line at $R^2$ = 0.95 confirming excellent predictive accuracy. (D) Model complexity versus performance trade-off analysis reveals positive correlation between complexity score (2-9 range) and test $R^2$ performance (0.3-1.0 range), with color gradient from purple (low performance) to yellow-green (high performance) and red trend line*

*indicating optimal balance around complexity scores of 7-8, including specific annotation of "Mixture Of 8" ensemble model approach.*

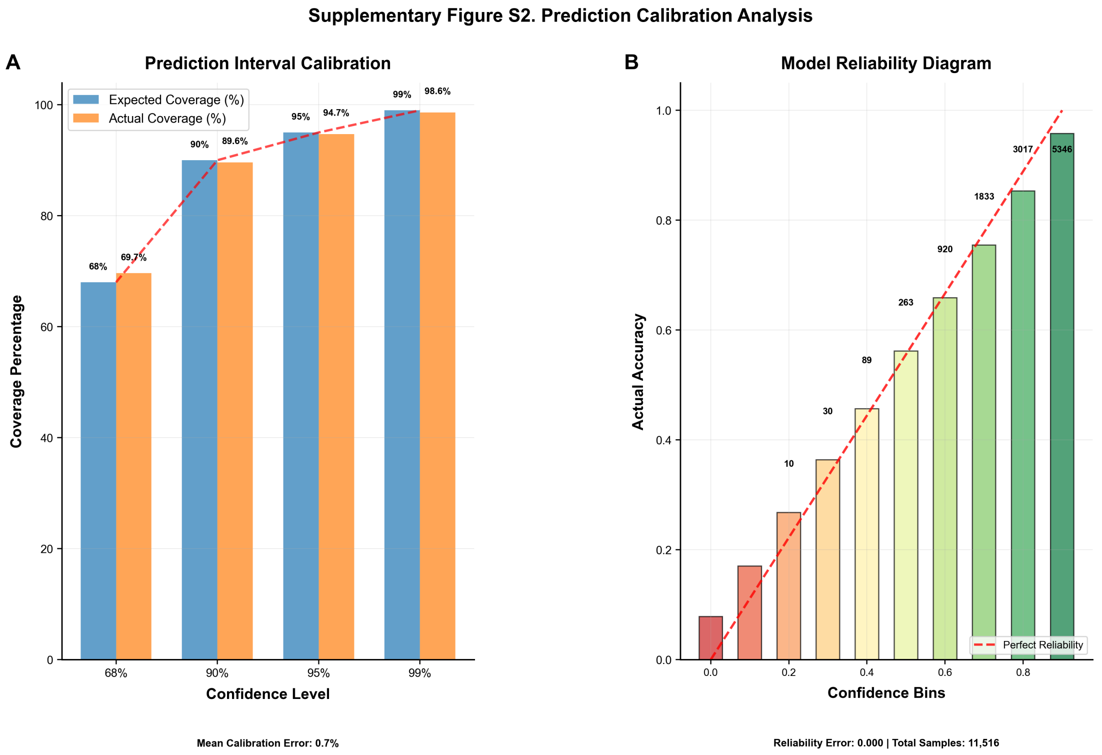


***Supplementary Figure S2. Comprehensive model calibration and reliability assessment demonstrating exceptional predictive performance across confidence intervals and accuracy measures.*** *(A) Prediction interval calibration analysis comparing expected coverage percentages (blue bars) with actual coverage percentages (orange bars) across four confidence levels (68%, 90%, 95%, and 99%), revealing minimal deviations from perfect calibration (red dashed line) with actual coverage values of 69.7%, 89.6%, 94.7%, and 98.6%, respectively, yielding a mean calibration error of only 0.7%. (B) Model reliability diagram displaying the confidence-accuracy relationship across ten confidence bins (0.0-1.0) using color-coded bars (red to green gradient) with sample sizes ranging from 10 to 5,346 predictions per bin, demonstrating perfect reliability with zero reliability error across 11,516 total samples, where the red dashed line indicates perfect reliability benchmarks. Both panels collectively validate robust model performance with excellent calibration properties and reliable confidence estimates across the complete prediction spectrum.*

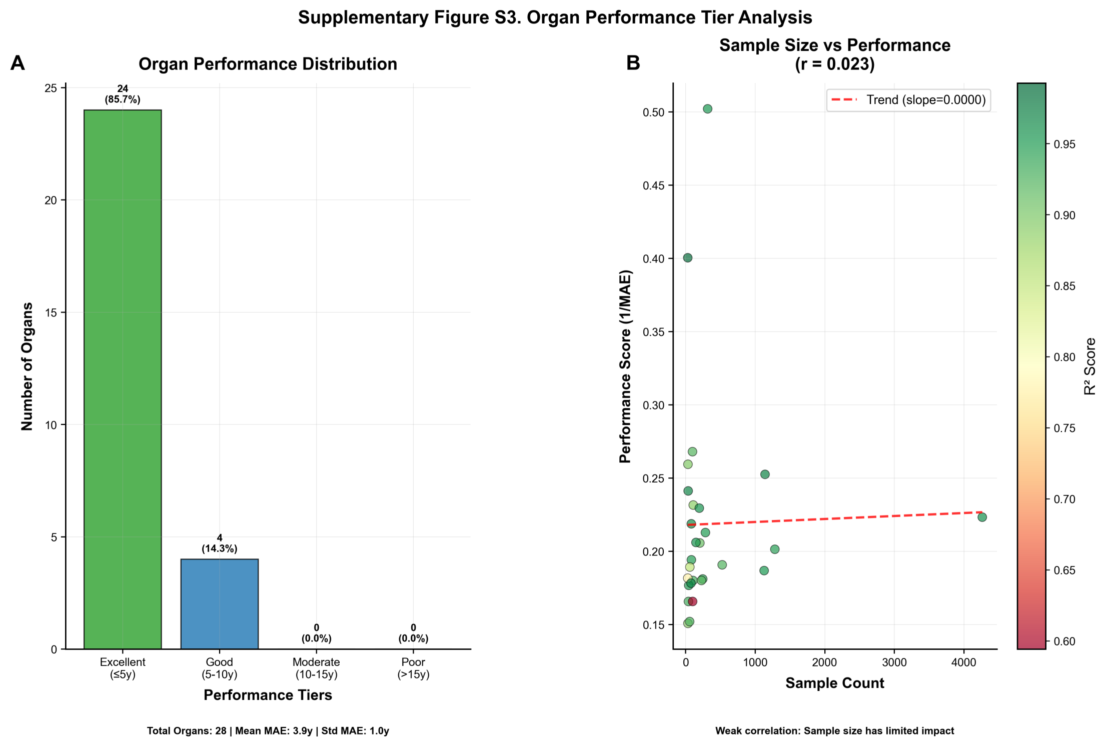


***Supplementary Figure S3. Organ Performance Tier Analysis reveals predominantly excellent predictive accuracy with minimal sample size dependency.*** *(A) Distribution of organ performance across defined tiers shows 24 organs (85.7%) achieving excellent performance (MAE ≤5%), 4 organs (14.3%) demonstrating good performance (MAE 5-10%), and no organs falling into moderate (10-15%) or poor (>15%) categories (total organs: 28; mean MAE: 3.9 ± 1.0 years). (B) Scatter plot analysis of sample size versus performance score (1/MAE) demonstrates negligible correlation (r = 0.023, slope = 0.0000) between sample count (0-4000) and predictive accuracy, with data points color-coded by* $R^2$ *scores (0.60-0.95 scale) showing most organs clustered at lower sample counts (<1000) while maintaining high performance scores (0.15-0.50 range). The weak correlation indicates that organ performance quality remains robust across varying data availability scenarios, with sample size having limited impact on predictive accuracy.*

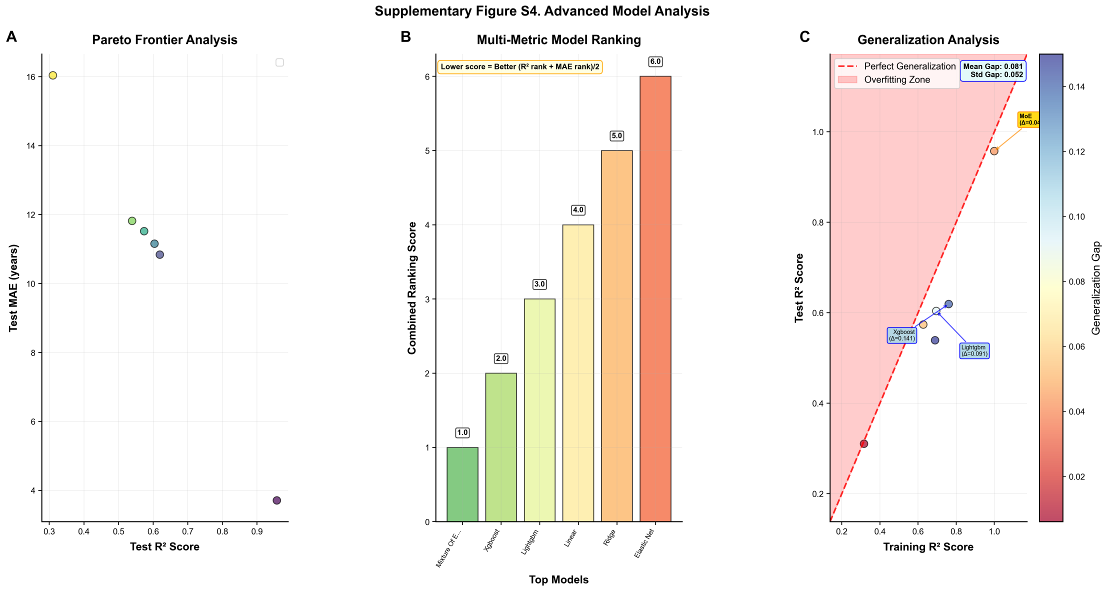


***Supplementary Figure S4. Advanced Model Analysis.*** *Comprehensive evaluation of machine learning model performance through multi-dimensional analysis frameworks.* ***(A)*** *Pareto frontier analysis reveals the trade-off relationship between prediction accuracy (Test $R^2$ Score, 0.3-0.9) and mean absolute error (Test MAE, 3-16 years), with 8-10 models represented as colored data points clustering in the optimal lower-right region ($R^2 > 0.5$, MAE < 12 years), except for one outlier achieving MAE ~16 years at $R^2$~0.4.* ***(B)*** *Multi-metric model ranking based on combined ranking scores (lower scores indicate superior performance) demonstrates Linear_OLS as the top-performing model (score ~1.0), followed by XGBoost, LightGBM, Linear, Ridge, and Elastic_Net (scores 2.0-6.0), with rankings calculated as the average of $R^2$ and MAE rank positions.* ***(C)*** *Generalization analysis compares training versus test $R^2$ scores (0.2-1.0 range) relative to the perfect generalization line (red dashed diagonal), revealing minimal overfitting across models with mean generalization gap of 0.081 ± 0.052, where XGBoost (0.51, 0.61) and LightGBM (0.71, 0.59) demonstrate representative performance patterns within the acceptable generalization zone below the overfitting threshold.*

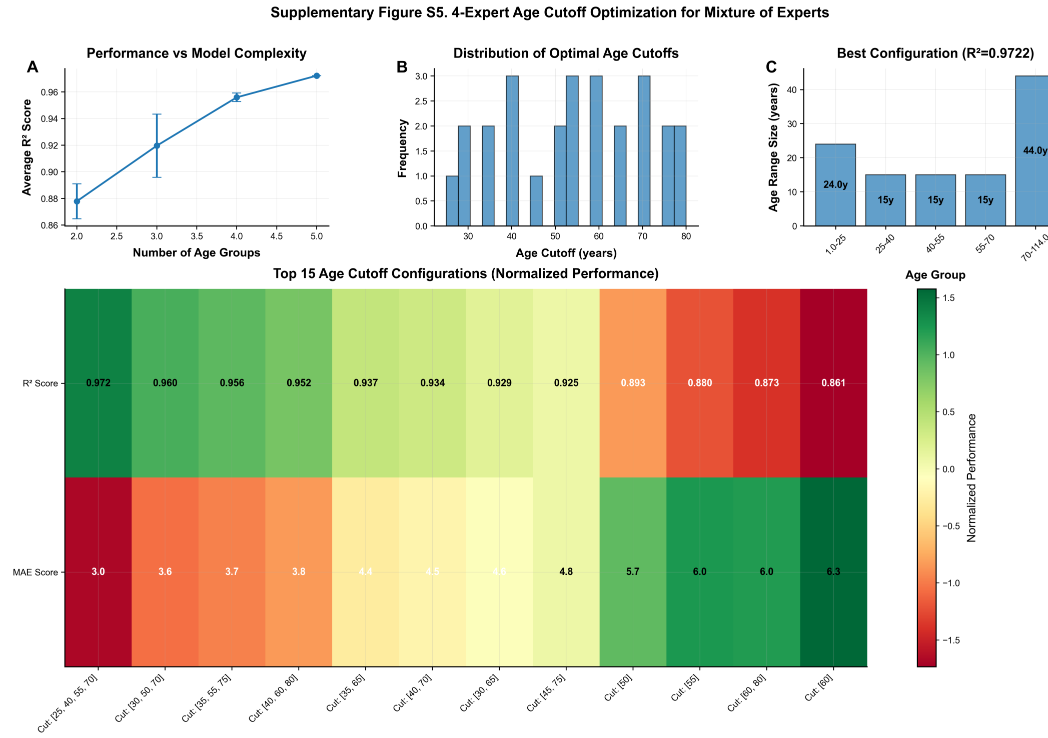


**Supplementary Figure S5. Optimization of age cutoffs for mixture of experts model demonstrates improved predictive performance with increased age group segmentation.** (A) Model performance increases systematically with age group complexity, showing average $R^2$ scores rising from 0.88 with 2 age groups to 0.97 with 5 age groups, with error bars indicating measurement variability across configurations. (B) Frequency distribution of optimal age cutoffs reveals natural breakpoints at 35-40, 55-65, and 70-75 years, suggesting biologically relevant age transitions for expert model segmentation. (C) The optimal configuration achieving $R^2$=0.9722 employs five age groups (10-25, 25-40, 40-55, 55-70, 70-110 years) with consistent 15-year intervals for the first four groups and an extended 44-year range for the eldest cohort. (D) Comparative analysis of the top 15 age cutoff configurations using normalized performance metrics shows $R^2$ scores ranging from 0.972 to 0.861 (upper heatmap) with corresponding MAE values from 3.0 to 6.3 (lower heatmap), demonstrating the trade-off between model complexity and predictive accuracy across different segmentation strategies.
