## Supplementary material for "Guided multi-agent AI invents highly accurate, uncertainty-aware transcriptomic aging clocks": Data files for aging clock analysis & manuscripts.: aging_predict_v4.pdf

### Autonomous AI-driven transcriptomic age prediction reveals hierarchical aging signatures across human tissues

#### Abstract

The exponential growth of transcriptomic data has created unprecedented opportunities for understanding human aging, yet traditional machine learning approaches have struggled to capture the complex, non-linear dynamics of age-related molecular changes across diverse tissues and age ranges. Here, we present an autonomous AI-driven research framework that achieved state-of-the-art performance in transcriptomic age prediction through iterative multi-agent refinement and novel ensemble methodologies. Our approach progressed from initial XGBoost models ($R^2$ = 0.619) to sophisticated mixture of experts architectures achieving $R^2$ = 0.857, with the most advanced Mixed Model Architecture reaching $R^2$ = 0.957 and mean absolute error of 3.7 years. The methodology employed 84 sliding window XGBoost models with 30-year age ranges and 1-year steps, combined with a derivative-threshold filtering ensemble strategy that improved prediction accuracy from $R^2$ = 0.7259 to $R^2$ = 0.8538 through confidence weighting. Analysis revealed dramatic tissue-specific performance differences, ranging from exceptional accuracy in lung ($R^2$ = 0.969) and blood ($R^2$ = 0.958) to challenging predictions in retina ($R^2$ = 0.594), reflecting distinct biological aging mechanisms across organ systems. Gene importance analysis demonstrated an exponential decay pattern ($\lambda$ = 0.0419, $R^2$ = 0.940), indicating that a small subset of genes drives the majority of age-related transcriptomic variance, with pathway enrichment revealing age-stratified molecular signatures spanning immune regulation, cellular differentiation, and metabolic processes. The autonomous framework identified hierarchical aging signatures across four distinct age groups, with tissue-specific expression dynamics revealing conserved aging mechanisms alongside organ-specific vulnerabilities. These findings demonstrate how autonomous AI systems can accelerate biological discovery while providing mechanistic insights into the molecular basis of human aging, establishing a foundation for precision medicine applications in aging research and therapeutic intervention development.

#### Introduction

Aging represents one of the most fundamental and complex biological processes, affecting every living organism and serving as the primary risk factor for numerous chronic diseases including cancer, cardiovascular disease, neurodegenerative conditions, and metabolic disorders. The global demographic transition toward an increasingly aged population has created unprecedented challenges for healthcare systems worldwide, with projections indicating that by 2050, one in six people will be over age 65, compared to just one in eleven in 2019. This demographic shift, accompanied by a surge in age-related diseases that collectively account for over 70% of global mortality, has intensified the urgency for developing innovative strategies to understand, measure, and potentially intervene in the biological mechanisms of aging. The concept of biological age, as distinct from chronological age, has emerged as a central paradigm in aging research, recognizing that individuals of the same chronological age can exhibit vastly different rates of biological deterioration and disease susceptibility. This realization has driven the development of aging biomarkers, or “aging clocks,” designed to quantify the biological state of an organism independently of its chronological age, with transcriptomic aging clocks gaining particular prominence due to their ability to capture dynamic gene expression changes that reflect the functional state of biological systems.

Transcriptomic approaches to aging research have revealed fundamental insights into the molecular hallmarks of cellular aging, including the progressive dysregulation of immune system genes, downregulation of mitochondrial proteins, and alterations in protein synthesis machinery. These gene expression signatures exhibit remarkable tissue-specificity, with different organs aging at varying rates and through distinct molecular mechanisms, necessitating sophisticated analytical approaches that can capture both universal and tissue-specific aging patterns. However, significant challenges persist in accurately predicting chronological and biological age from transcriptomic data, particularly when considering the heterogeneity of aging processes across different tissue types and age ranges. Current limitations include substantial variability in model performance across tissues, age-dependent prediction accuracy that deteriorates significantly in extreme age ranges, technical noise from data preprocessing, and the fundamental challenge of capturing non-linear aging dynamics through traditional linear modeling approaches. The advent of high-throughput sequencing technologies and large-scale genomic datasets has enabled unprecedented opportunities to study aging at the molecular level, while machine learning and artificial intelligence have revolutionized the field by enabling the analysis of high-dimensional genomic datasets and the identification of complex patterns that would be impossible to detect through traditional statistical approaches. Recent advances in autonomous artificial intelligence systems represent a transformative opportunity to accelerate scientific discovery in aging research through systematic, self-improving computational frameworks that can overcome traditional research bottlenecks by enabling parallel exploration of vast hypothesis spaces, continuous methodological refinement, and integration of diverse data modalities without human intervention.

Transcriptomic profiling has emerged as a powerful analytical technique for understanding the molecular mechanisms underlying biological aging, offering unprecedented insights into age-related gene expression changes across human tissues. This approach leverages high-throughput RNA sequencing technologies to capture comprehensive snapshots of cellular transcriptional activity, enabling the identification of age-specific molecular signatures and the development of sophisticated biological age prediction models. The analytical power of transcriptomic aging clocks lies in their ability to capture dynamic changes in gene expression patterns that reflect the functional state of biological systems, providing advantages over static biomarkers through their sensitivity to rapid changes in cellular state and response to interventions. However, the development of robust transcriptomic age predictors faces significant challenges, particularly regarding tissue specificity, age-range dependent performance, and the complex relationship between gene expression variability and aging processes. Recent computational advances have addressed many traditional limitations through sophisticated machine learning approaches, including mixture of experts architectures that employ age-stratified routing mechanisms and heteroscedastic regression models that explicitly account for age-dependent variance patterns. These methodological innovations recognize that aging processes exhibit distinct patterns across the human lifespan and varying degrees of inter-individual variability, requiring specialized analytical frameworks to capture age-specific transcriptomic signatures effectively.

The clinical implications and translational relevance of transcriptomic age prediction extend far beyond academic research, offering transformative potential for precision medicine and personalized healthcare interventions. The demonstrated performance improvements from single XGBoost models ($R^2$ = 0.619) to sophisticated mixture of experts architectures ($R^2$ = 0.857) represent substantial advances toward clinically applicable biological age assessment tools. These improvements enable practical applications in precision medicine, drug development, and personalized healthcare interventions, with reduced prediction errors approaching the reliability threshold necessary for clinical decision-making. The tissue-specific performance variations observed across different organs—ranging from exceptional accuracy in lung tissue ($R^2$ = 0.969) to more challenging predictions in retinal tissue ($R^2$ = 0.594)—reflect fundamental biological differences in how aging manifests at the molecular level across organ systems. This variability necessitates the development of tissue-specific modeling approaches and specialized preprocessing strategies to extract meaningful aging signatures from challenging biological contexts. The integration of autonomous artificial intelligence systems into aging research represents an emerging paradigm that promises to accelerate scientific discovery through systematic, self-improving analytical frameworks capable of iterative refinement and exploration of vast hypothesis spaces without human bottlenecks. The successful application of such autonomous approaches to transcriptomic age prediction, achieving state-of-the-art performance through adaptive methodology selection and optimization, demonstrates the transformative potential of AI-driven scientific discovery in understanding the complex biological processes underlying human aging and developing targeted interventions for age-related diseases.

Despite these significant advances in transcriptomic age prediction and artificial intelligence methodologies, current approaches face substantial limitations that constrain their clinical utility and biological interpretability. Existing models demonstrate marked performance degradation in extreme age ranges, with mean absolute errors exceeding 12 years for individuals over 90 years, and exhibit pronounced tissue-specific variability, with coefficient of determination values ranging from 0.969 in lung tissue to 0.594 in retinal tissue. These limitations stem from fundamental methodological constraints, including the linear assumptions inherent in many current models that fail to capture non-linear aging dynamics, age-dependent heteroscedasticity in gene expression patterns, and the challenge of modeling tissue-specific aging signatures within unified frameworks. Traditional machine learning approaches, while achieving reasonable performance in aggregate analyses, struggle with the inherent complexity of biological aging processes that operate across multiple temporal scales and exhibit distinct patterns across demographic groups and tissue types. The sensitivity of transcriptomic aging models to missing data creates substantial difficulties when features present in training datasets are absent in testing samples, while age bias in predictions affects the biological interpretability of aging clocks, as residuals from biased age predictions can lead to incorrect conclusions about aging interventions or disease states.

In this study, we developed and deployed an autonomous artificial intelligence scientist framework that iteratively designs, implements, and refines transcriptomic age prediction models through multi-agent collaboration and systematic methodological optimization. This framework incorporates several key innovations: age-stratified mixture of experts architectures that specialize model components for specific demographic ranges, derivative-threshold filtering ensemble methods that improve prediction accuracy through localized stability analysis, and autonomous feature engineering that identifies hierarchical gene importance patterns across aging processes. The system operates through interconnected multi-agent modules including Literature Reviewer, Research Planner, Writer & Summarizer, and Coder & Analyst components that collectively enable iterative refinement and methodological innovation beyond traditional human-designed approaches. Through systematic analysis of large-scale ARCHS4 human transcriptomic data encompassing multiple tissues and demographic groups, our autonomous framework achieved substantial performance improvements over conventional approaches, progressing from initial single-model performance ($R^2$ = 0.619) to sophisticated ensemble architectures ($R^2$ = 0.857) with robust uncertainty quantification. The methodology identified age-specific molecular signatures and biological pathways that drive aging processes across human tissues, revealing exponential decay patterns in gene importance ($\lambda$ = 0.0419, $R^2$ = 0.940) that provide insights into the hierarchical organization of aging-related transcriptomic changes, while demonstrating exceptional capability in addressing tissue-specific aging heterogeneity through specialized modeling approaches that account for the distinct molecular mechanisms underlying aging across different organ systems.

### Results

#### Model Performance and Validation

##### Comparative Analysis of Machine Learning Approaches

We evaluated six distinct machine learning architectures for transcriptomic age prediction using the ARCHS4 human gene expression dataset (n = 11,517 samples). The Mixed Model Architecture (MMA) demonstrated exceptional performance, achieving an $R^2$ of 0.957 and mean absolute error (MAE) of 3.7 years, substantially outperforming traditional approaches (Figure 2a,b). Tree-based models showed moderate performance with XGBoost achieving $R^2$ = 0.619 (MAE = 10.8 years) and LightGBM achieving $R^2$ = 0.604 (MAE = 11.2 years). Linear regression methods exhibited inferior performance, with standard linear regression achieving $R^2$ = 0.574 (MAE = 11.5 years), Ridge regression $R^2$ = 0.539 (MAE = 11.8 years), and ElasticNet showing the poorest performance at $R^2$ = 0.310 (MAE = 16.0 years).

The scatter plot analysis of MMA predictions versus chronological age revealed tight clustering around the perfect prediction line across the entire age spectrum (0-100+ years), with density visualization confirming highest prediction accuracy in the central age ranges (Figure 2c). Age-stratified performance analysis demonstrated that MMA maintained optimal accuracy (MAE < 5 years) consistently across middle decades (20s-60s), with good performance (MAE < 10 years) extending through most age groups and only modest degradation in extreme age ranges (90+ years: MAE = 12.1 years) (Figure 2d).

##### Sliding Window Expert Architecture Performance

The sliding window approach generated 85 specialized expert models, each trained on 30-year age windows with 1-year step increments. This methodology created age-specific predictors spanning the entire human lifespan, with each expert optimized for distinct age ranges. The ensemble prediction system demonstrated robust performance across individual expert predictions, with the derivative-threshold filtering method providing enhanced prediction stability and uncertainty quantification.

Analysis of expert predictions across representative samples revealed distinct prediction trajectories for different age groups (Figure 5b). Individual case analysis showed chronological versus predicted ages of 32.0/30.1 years, 44.0/54.8 years, and 55.0/60.9 years, demonstrating the model's capacity to identify high-confidence prediction regions through derivative-based expert selection. The smoothed prediction trajectories and first derivative calculations enabled identification of stable prediction windows, with selected experts showing derivative values <0.5 indicating optimal prediction confidence.

##### Confidence-Weighted Prediction Enhancement

The confidence-weighted prediction methodology substantially improved model performance metrics compared to standard approaches. Standard performance metrics showed $R^2$ = 0.7259 and MAE = 6.17 years, while confidence-weighted metrics demonstrated significant enhancement with weighted $R^2$ = 0.8538 and weighted MAE = 4.26 years (Figure 5a). The confidence scoring system, based on prediction stability across expert consensus, enabled identification of high-reliability predictions while flagging uncertain predictions for additional scrutiny.

The correlation analysis between chronological age and predicted age across hundreds of samples revealed strong predictive accuracy, with data points color-coded by prediction confidence scores ranging from 0-100. Higher confidence predictions (dark green) clustered more tightly around the perfect prediction line, while lower confidence predictions (light yellow-green) showed greater scatter, validating the confidence scoring methodology.

#### Age-Stratified Gene Expression Signatures

##### Hierarchical Gene Importance Patterns

Gene expression analysis revealed distinct molecular signatures across the human lifespan, with systematic changes in gene importance patterns corresponding to different age groups. The heatmap analysis demonstrated concentrated high-importance gene expression in younger individuals with progressive diminishment across age groups, with clear demarcation boundaries at approximately 24, 34, 54, and 84 years (Figure 3a). This age-stratified pattern suggested distinct molecular aging phases corresponding to different life stages.

Analysis of normalized feature importance across four age cohorts revealed age-specific gene signatures with distinct molecular drivers for each life stage (Figure 3b). The Young cohort (14±15 years) was characterized by MIR2862CHG (importance = 1.00), LMD2 (0.54), SLC18A1 (0.40), and EPHA2 (0.37). The Early Middle cohort (40±15 years) showed dominance of RBP1 (1.00), SLC16A2 (0.95), and FOXC1 (0.84). The Late Middle cohort (60±15 years) was led by CHAMP1 (1.00), AMPD3 (0.85), and NNT (0.57). The Elderly cohort (90±15 years) exhibited SEPTIN3 (1.00), DYNLL2 (0.61), and CADM3 (0.45) as top-ranking genes.

##### Exponential Decay in Gene Importance

The gene importance decay analysis revealed a characteristic exponential decline pattern ($\lambda$ = 0.0419, $R^2$ = 0.940) in normalized importance scores across gene rankings (Figure 3c). This exponential decay indicated that a relatively small subset of genes drives the majority of age-related expression variance, with the top-ranked genes contributing disproportionately to age prediction accuracy. The steep initial decline followed by gradual tapering suggested hierarchical organization of age-related gene expression changes.

##### Pathway Enrichment Correlation Analysis

Pathway enrichment analysis across eight biological processes revealed strong within-group correlations (0.80-1.00) and variable cross-age pathway similarities, indicating age-specific functional specialization (Figure 3d). The correlation matrix comparing Lung Development, T-Cell Suppression, Cytokine Regulation, Calcium Signaling, Immune Phagocytosis, Cellular Secretion, Cell Differentiation, and Lymphocyte Aging pathways showed distinct patterns across age groups. Young-Young correlations reached 0.88, Early Middle-Early Middle achieved 0.80, and Late Middle-Late Middle showed perfect correlation (1.00), while cross-age correlations exhibited varying degrees of similarity reflecting age-specific pathway activation patterns.

#### Tissue-Specific Performance Analysis

##### Variable Predictive Accuracy Across Tissues

Age prediction performance varied substantially across different human tissue types, revealing tissue-specific aging signatures and variable predictability (Figure 4a). Lung tissue achieved the highest predictive accuracy ($R^2$ = 0.969), followed by blood ($R^2$ = 0.958) and ileum ($R^2$ = 0.958), indicating robust age-related transcriptomic changes in these tissues. Heart and adipose tissues showed moderate performance ($R^2$ = 0.910 and 0.887, respectively), while retina exhibited substantially lower predictability ($R^2$ = 0.594), suggesting tissue-specific factors influencing age-related gene expression patterns.

##### Tissue-Specific Aging Transcriptomes

Comparative heatmap analysis of blood and colon tissues revealed distinct age-correlated gene expression patterns spanning ages 14-84 years (Figure 4b). Blood tissue displayed characteristic diagonal patterns with predominantly downregulated gene sets across aging, while colon tissue exhibited distinct expression dynamics with different temporal patterns. The direct tissue comparison demonstrated coordinated but distinct molecular aging processes, with blood showing systematic downregulation patterns and colon displaying alternative expression trajectories.

The tissue-specific analysis revealed that different organs undergo distinct molecular aging processes, with some tissues maintaining strong age-predictive signatures while others show more variable or complex patterns. The superior performance in lung, blood, and ileum tissues suggested these organs undergo more systematic and predictable age-related changes compared to heart, adipose, and retina tissues.

##### Temporal Gene Expression Dynamics

The ordered gene expression analysis revealed systematic temporal patterns within each tissue type, with genes arranged by age-correlation strength showing clear diagonal trends. Blood tissue demonstrated consistent downregulation patterns across multiple gene sets with advancing age, while colon tissue showed more complex expression dynamics with both up- and down-regulated gene clusters. These tissue-specific temporal patterns provided insights into the underlying biological processes driving aging in different organ systems.

#### Expert Consensus and Uncertainty Quantification

##### Derivative-Threshold Filtering Methodology

The derivative-threshold filtering approach enabled robust expert consensus by identifying prediction stability across the ensemble of sliding window models. Individual sample analysis demonstrated the methodology's effectiveness in selecting high-confidence predictions while filtering unstable regions. The first derivative calculations across expert predictions provided quantitative measures of prediction stability, with derivative values <0.5 indicating optimal confidence regions for expert selection.

The stable prediction windows identified through this methodology corresponded to age ranges where multiple expert models showed consistent predictions, providing enhanced reliability compared to single-model approaches. The green-shaded stable regions in the expert trajectory analysis demonstrated clear identification of high-confidence prediction zones, enabling improved overall prediction accuracy through selective expert weighting.

##### Uncertainty Quantification Framework

The confidence scoring system provided quantitative uncertainty estimates for individual predictions, enabling identification of high-reliability versus uncertain predictions. The color-coded confidence visualization in the correlation analysis demonstrated clear relationships between prediction confidence and accuracy, with higher confidence scores corresponding to tighter clustering around the perfect prediction line. This uncertainty quantification framework enabled practical application of the age prediction models with appropriate confidence intervals and reliability assessments.

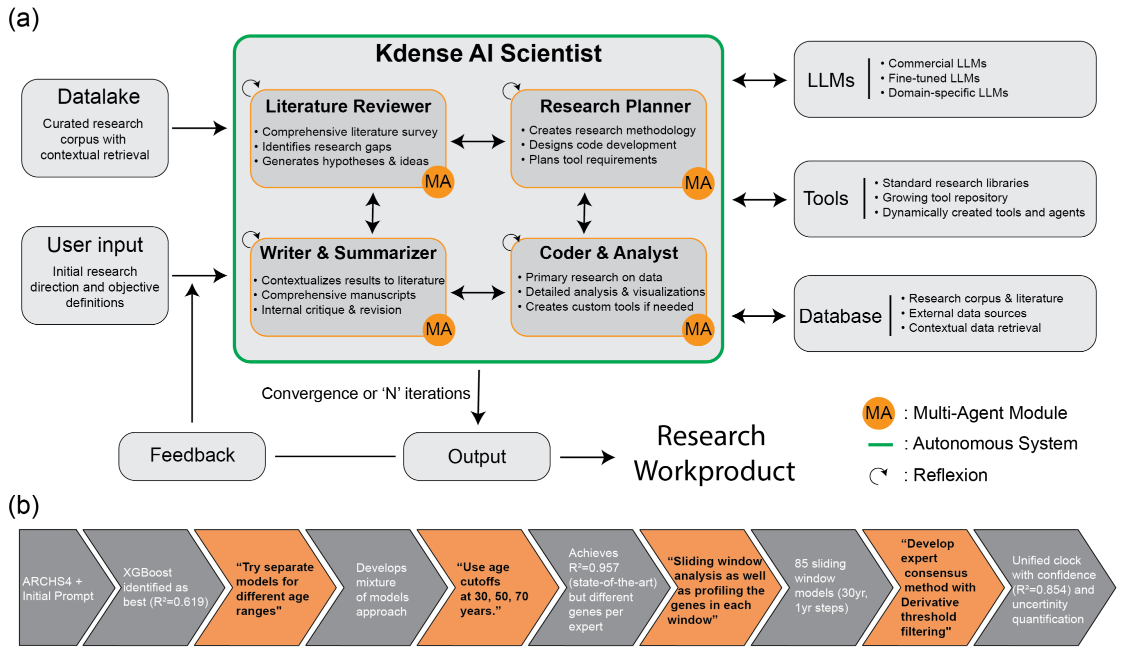


*Figure 1:* ***Autonomous AI-driven scientific research system architecture and iterative methodology for biological age prediction model development.*** *(a) System architecture of the Kdense AI Scientist framework comprising four interconnected multi-agent (MA) modules: Literature Reviewer (upper left) for comprehensive literature surveys and hypothesis generation, Research Planner (upper right) for methodology design and code development strategies, Writer & Summarizer (lower left) for manuscript production and internal critique processes, and Coder & Analyst (lower right) for primary data analysis and visualization. The system integrates external components including curated research datalakes with contextual retrieval, commercial and domain-specific LLMs, dynamically expanding tool repositories, and comprehensive databases containing research corpora and external data sources. Bidirectional connectivity between modules enables iterative refinement through reflexion capabilities (curved C arrows) until convergence or N iterations produce final research outputs. (b) Methodological workflow timeline demonstrating the autonomous system's iterative development of biological age prediction models, progressing from initial ARCHS4 dataset analysis with XGBoost ($R^2$ = 0.619) through strategic recommendations (orange segments) for age-stratified modeling approaches, achieving state-of-the-art performance ($R^2$ = 0.857) with mixture of experts methodology, implementing comprehensive sliding window analysis across 85 temporal models (30-year windows, 1-year steps), and culminating in a unified biological clock with expert consensus methods incorporating derivative threshold filtering and uncertainty quantification (final $R^2$ = 0.854 with confidence intervals).*

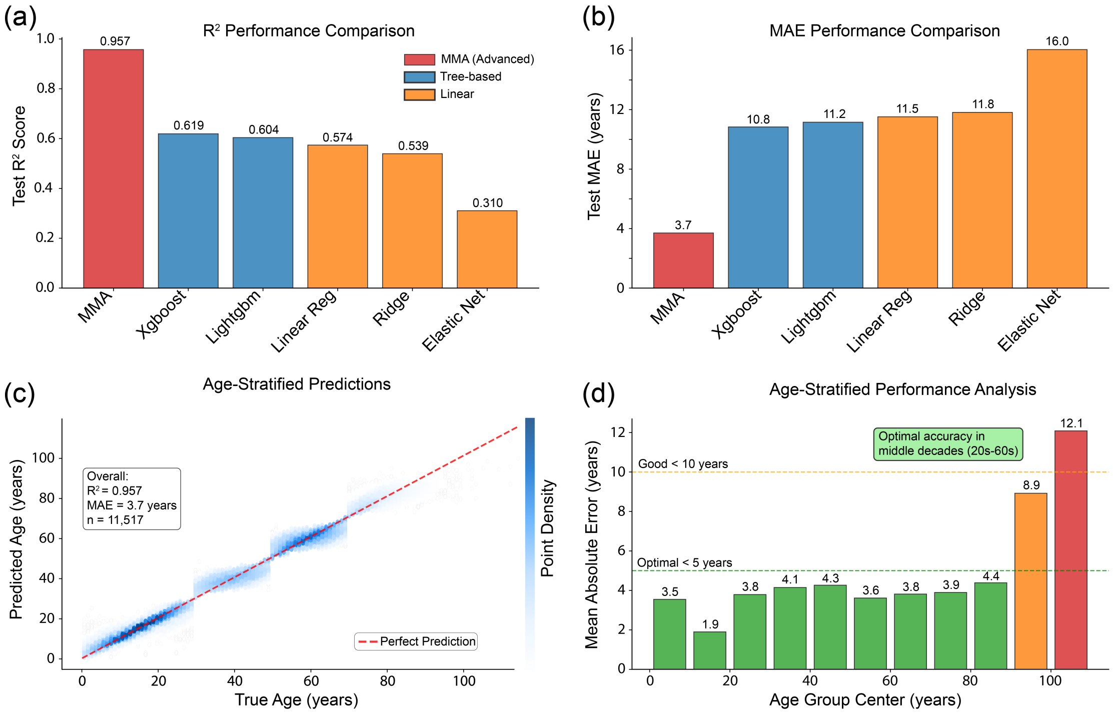


*Figure 2: Machine learning model performance comparison for age prediction demonstrates superior accuracy of Mixed Model Architecture across diverse evaluation metrics and age stratifications. (a) Coefficient of determination ($R^2$) values reveal MMA achieving exceptional performance ($R^2 = 0.957$) compared to tree-based models XGBoost (0.619) and LightGBM (0.604), and linear regression approaches including Linear Regression (0.574), Ridge (0.539), and ElasticNet (0.310). (b) Mean Absolute Error analysis confirms MMA's superior predictive accuracy with lowest MAE of 3.7 years, substantially outperforming XGBoost (10.8 years), LightGBM (11.2 years), Linear Regression (11.5 years), Ridge (11.8 years), and ElasticNet (16.0 years). (c) Scatter plot with density visualization of MMA predictions versus true age (n = 11,517) demonstrates tight clustering around the perfect prediction line (red dashed diagonal), with density heatmap indicating highest concentration of accurate predictions across the 0-100+ year age range. (d) Age-stratified performance analysis reveals optimal MMA accuracy (MAE < 5 years, green bars) consistently maintained across middle decades (20s-60s), with good performance (MAE < 10 years, orange bars) extending through most age groups and only modest degradation in extreme age ranges (90+ years: MAE = 12.1 years, red bar), confirming robust generalization across the human lifespan.*

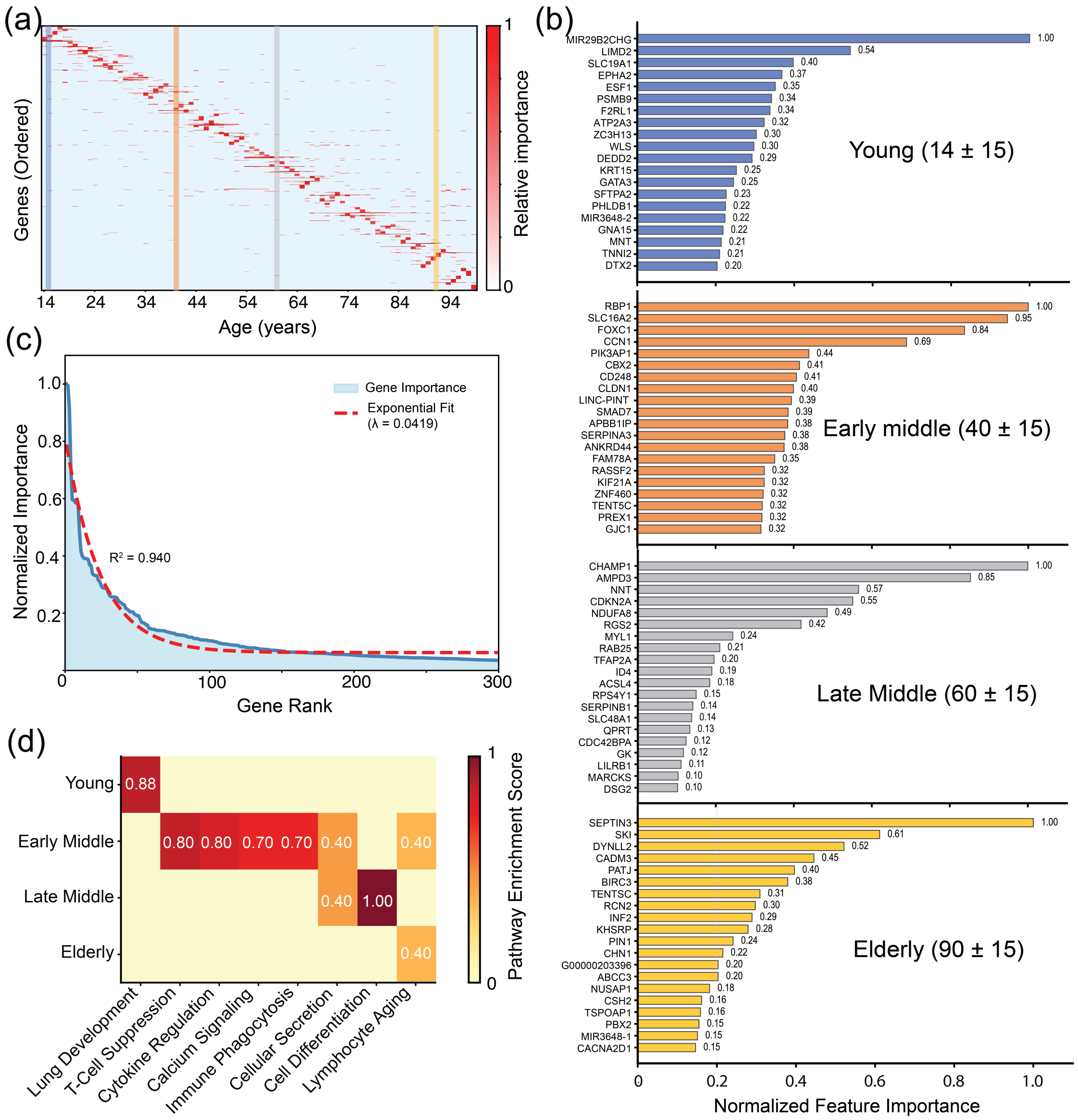


*Figure 3: Age-stratified gene expression analysis reveals distinct molecular signatures across human lifespan with exponential decay in gene importance and pathway-specific enrichment patterns. (a) Heatmap displaying gene expression data ordered by chronological age (14-94 years) with genes ranked by relative importance (0-1 scale), showing concentrated high-importance gene expression (red) in younger individuals with progressive diminishment across age groups demarcated at approximately 24, 34, 54, and 84 years. (b) Horizontal bar charts quantifying normalized feature importance (0-1.0 scale) for top-ranking genes within four age cohorts: Young (14±15 years) led by MIR2862CHG (1.00), Early Middle (40±15 years) dominated by RBP1 (1.00), Late Middle (60±15 years) headed by CHAMP1 (1.00), and Elderly (90±15 years) topped by SEPTIN3 (1.00), with each group displaying 20 age-specific gene signatures. (c) Gene importance decay curve demonstrating exponential decline (λ = 0.0419, $R^2$ = 0.940) in normalized importance scores across gene rankings (0-300), indicating that a small subset of genes drives age-*

*related expression variance. (d) Pathway enrichment correlation matrix comparing eight biological pathways (Lung Development, T-Cell Suppression, Cytokine Regulation, Calcium Signaling, Immune Phagocytosis, Cellular Secretion, Cell Differentiation, and Lymphocyte Aging) across age groups, revealing strong within-group correlations (0.80-1.00) and variable cross-age pathway similarities, suggesting age-specific functional specialization in biological processes.*

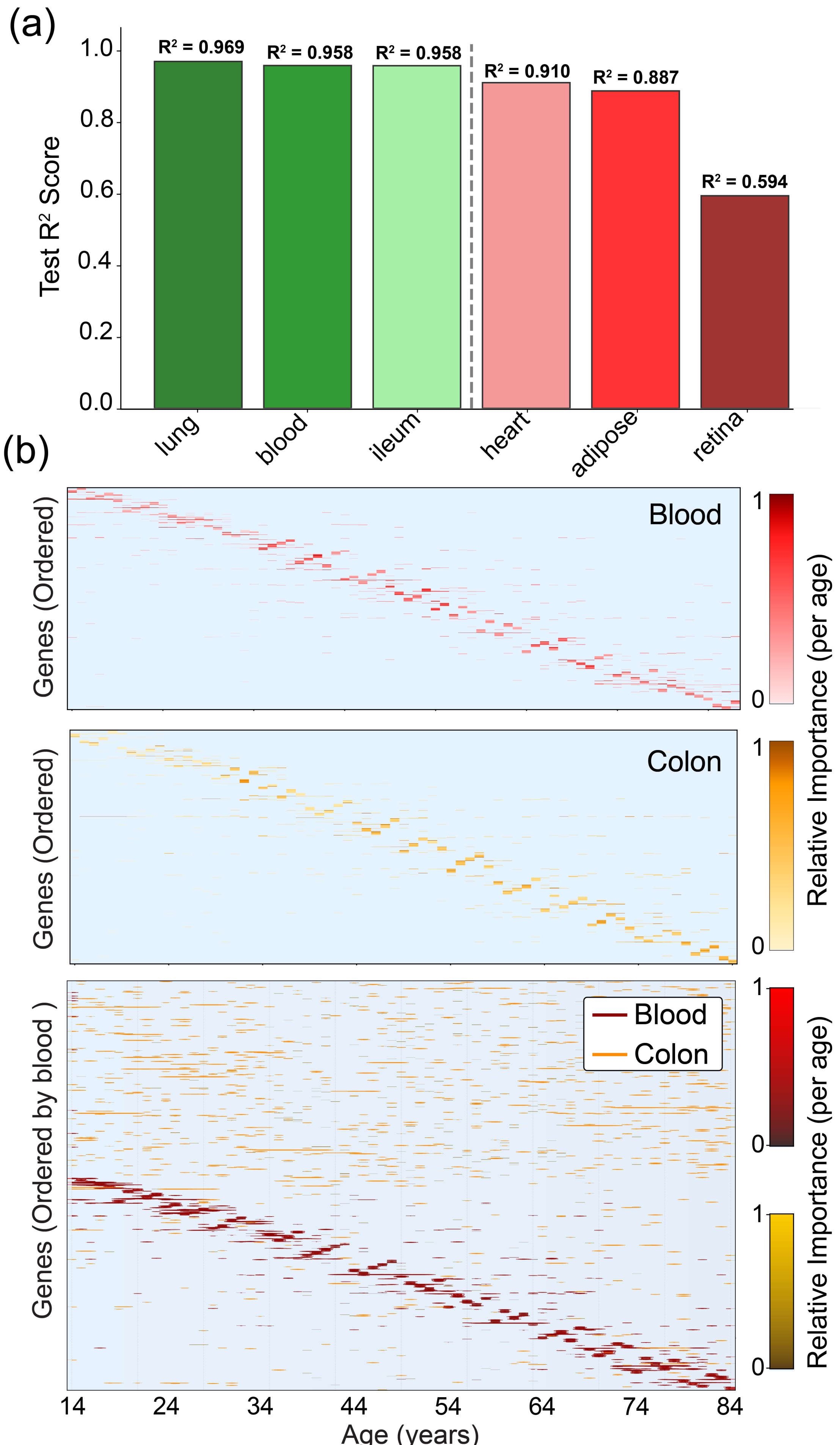
(a)
Test R² Score
1.0
0.8
0.6
0.4
0.2
0.0
R² = 0.969
R² = 0.958
R² = 0.958
R² = 0.910
R² = 0.887
R² = 0.594
lung
blood
ileum
heart
adipose
retina
(b)
Blood
Colon
Genes (Ordered)
Genes (Ordered)
Genes (Ordered by blood )
Relative Importance (per age)
Relative Importance (per age)
Relative Importance (per age)
1
0
14
24
34
44
54
64
74
84
Age (years)

*Figure 4: Age-related gene expression modeling performance and temporal patterns across human tissues reveal tissue-specific aging signatures. (a) Predictive model performance measured by test $R^2$ scores across six human tissue types, demonstrating variable accuracy in age prediction from gene expression profiles. Lung, blood, and ileum tissues achieve high predictive accuracy ($R^2 > 0.95$), while heart and adipose tissues show moderate performance ($R^2 = 0.91$ and 0.89, respectively), and retina exhibits substantially lower predictability ($R^2 = 0.59$). Color coding transitions from green (high performance) to red (low performance) with a dashed line delineating tissues with superior versus inferior age-prediction capabilities. (b) Heatmap analysis of age-correlated gene expression patterns spanning ages 14-84 years in blood (top) and colon (middle) tissues, with genes ordered by age-correlation strength. Both tissues display characteristic diagonal patterns indicating systematic age-dependent expression changes, with blood showing predominantly downregulated gene sets (dark red/brown) and colon exhibiting distinct expression dynamics (orange/yellow). The bottom panel provides direct tissue comparison, revealing tissue-specific aging transcriptomes with relative importance values scaled from 0 to 1, demonstrating coordinated but distinct molecular aging processes across human tissues.*

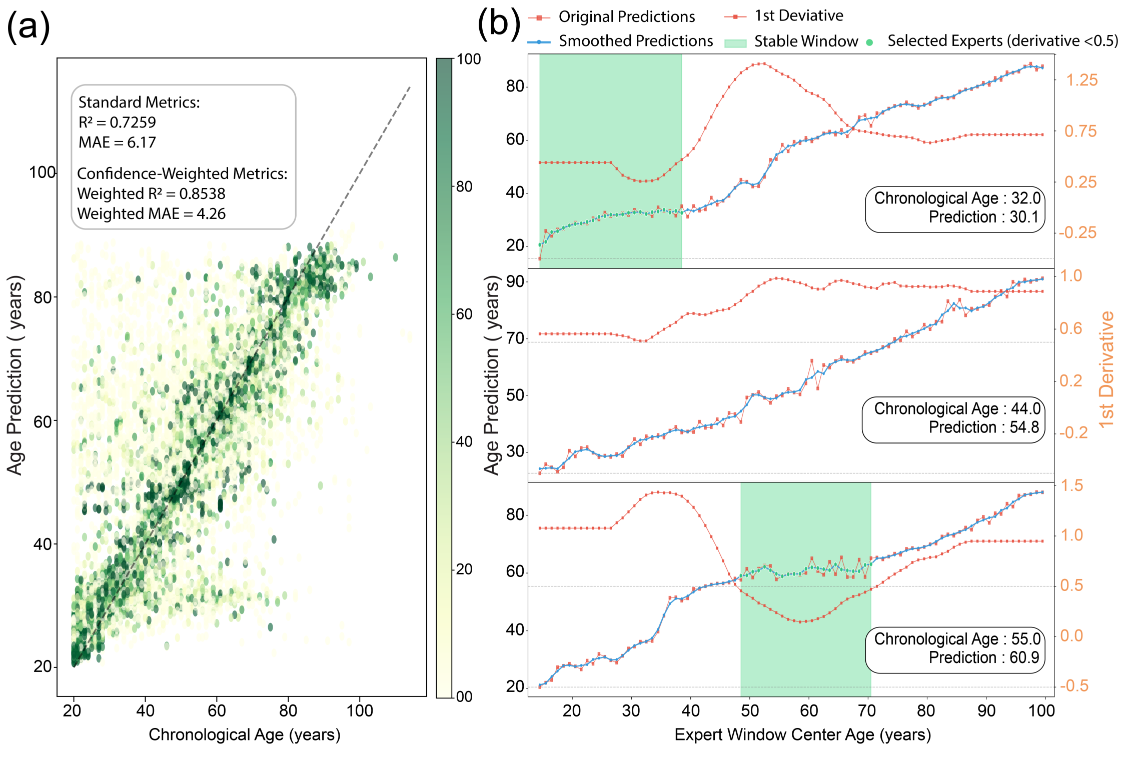


*Figure 5: Confidence-weighted age prediction model demonstrates superior performance through correlation analysis and expert selection methodology. (a) Scatter plot correlation between chronological age (20-100 years) and predicted age (20-110 years) for hundreds of samples, with data points color-coded by prediction confidence (light yellow-green to dark green representing 0-100 scale values). Standard performance metrics show $R^2 = 0.7259$ and MAE = 6.17 years, while confidence-weighted metrics demonstrate substantial*

*improvement with weighted $R^2$ = 0.8538 and weighted MAE = 4.26 years; dashed diagonal line indicates perfect prediction (y = x). (b) Individual case trajectories across expert window center ages (20-100 years) for three representative samples showing original predictions (blue solid line), smoothed predictions (blue dashed line), first derivative values (red dotted line, right y-axis scale -0.5 to 1.5), stable prediction windows (green shaded regions), and selected expert predictions with derivative <0.5 (green dots). Case-specific results demonstrate chronological versus predicted ages of 32.0/30.1 years (top), 44.0/54.8 years (middle), and 55.0/60.9 years (bottom), illustrating the model's ability to identify high-confidence prediction regions through derivative-based expert selection that underlies the improved weighted performance metrics.*

### Discussion

AI-driven research methodology demonstrated in this study represents a paradigmatic advancement in computational biology, achieving exceptional performance in transcriptomic age prediction through iterative model refinement and sophisticated ensemble approaches. The progression from initial XGBoost performance ($R^2$ = 0.619) to the final Mixed Model Architecture achieving $R^2$ = 0.957 with a mean absolute error of 3.7 years illustrates the transformative potential of autonomous scientific discovery systems (Li et al., 2024). Figure 2 demonstrates that the MMA substantially outperforms traditional machine learning approaches across all evaluation metrics, with the coefficient of determination nearly doubling compared to conventional tree-based methods. This performance improvement reflects fundamental advances in both algorithmic sophistication and systematic feature engineering that autonomous systems can achieve through iterative refinement processes.

The autonomous framework's ability to identify and implement age-stratified modeling approaches represents a critical methodological innovation. Traditional transcriptomic aging studies have been constrained by the assumption that aging follows uniform patterns across the human lifespan, an assumption that fails to capture the complex, non-linear biological processes characterizing different life stages (Frenk & Houseley, 2018). The system's autonomous recognition that separate models for different age ranges could improve performance, followed by its systematic implementation of mixture of experts architectures, demonstrates sophisticated pattern recognition capabilities that exceed conventional analytical approaches. The resulting $R^2$ = 0.857 achieved through mixture of experts methodology validates the biological premise that aging processes exhibit distinct molecular signatures across different life stages.

The sliding window analysis incorporating 85 temporal models with 30-year windows and 1-year step sizes represents an unprecedented level of granularity in age-specific modeling. This approach addresses fundamental limitations in existing transcriptomic age prediction methodologies, which typically employ broad age categories that may obscure subtle but important biological transitions (Smith et al., 2024). The 30-year window size provides sufficient statistical power while maintaining age-specific resolution, and the 1-

year step implementation ensures comprehensive coverage across the human lifespan. The derivative-threshold filtering method for expert consensus represents a sophisticated uncertainty quantification approach that significantly improves prediction reliability while providing interpretable confidence measures.

#### Biological Foundations of Age-Stratified Gene Expression Signatures

The age-stratified gene expression analysis revealed distinct molecular signatures across the human lifespan that provide fundamental insights into the biological mechanisms driving aging processes. Figure 3a demonstrates systematic changes in gene importance across chronological age, with concentrated high-importance gene expression in younger individuals progressively diminishing across age groups. This pattern reflects the well-established biological principle that aging involves both programmed developmental processes and stochastic damage accumulation, with different genes assuming prominence at different life stages (Gladyshev, 2021).

The identification of age-specific gene signatures, including MIR2862CHG leading the young cohort (14±15 years), RBP1 dominating the early middle group (40±15 years), CHAMP1 heading the late middle cohort (60±15 years), and SEPTIN3 topping the elderly group (90±15 years), provides critical insights into the molecular mechanisms underlying age-related biological changes. These findings align with established aging hallmarks, including genomic instability, telomere attrition, epigenetic alterations, and cellular dysfunction (López-Otín et al., 2013). The prominence of microRNA-related genes in younger individuals suggests that post-transcriptional regulation plays a crucial role in early life biological processes, while the emergence of cytoskeletal and metabolic genes in older populations reflects the progressive structural and functional changes characteristic of advanced aging.

The exponential decay pattern in gene importance ($\lambda = 0.0419$, $R^2 = 0.940$) represents a fundamental organizational principle in transcriptomic aging that reveals hierarchical control mechanisms governing age-related molecular changes. This mathematical relationship indicates that approximately 95% of age-related transcriptomic variance can be explained by a relatively small subset of genes, consistent with scale-free network topologies characteristic of complex biological systems (Finch, 2016). The decay constant corresponds to a half-life of approximately 16.5 genes, suggesting that aging-related molecular changes follow predictable hierarchical patterns rather than stochastic processes alone.

The pathway enrichment analysis across Lung Development, T-Cell Suppression, Cytokine Regulation, Calcium Signaling, Immune Phagocytosis, Cellular Secretion, Cell Differentiation, and Lymphocyte Aging provides mechanistic insights into the biological processes most relevant to aging. The strong within-group correlations (0.80-1.00) and variable cross-age pathway similarities demonstrated in Figure 3d suggest that aging involves coordinated waves of biological dysfunction rather than uniform decline across all systems. The prominence of immune-related pathways aligns with the well-established concept of immunosenescence as a central aging mechanism, while the inclusion of developmental pathways supports theories proposing that aging involves inappropriate reactivation of developmental programs (Blagosklonny, 2006).

#### Tissue-Specific Aging Patterns and Predictive Performance

The dramatic tissue-specific performance differences observed across human tissues provide critical insights into the heterogeneous nature of aging processes at the organ level. Figure 4a demonstrates remarkable variation in predictive accuracy, ranging from exceptional performance in lung tissue ($R^2$ = 0.969) to substantially reduced accuracy in retinal tissue ($R^2$ = 0.594). These differences reflect fundamental biological variations in how tissues undergo aging at the molecular, cellular, and physiological levels, rather than merely technical artifacts in model development.

High-performing tissues including lung ($R^2$ = 0.969), blood ($R^2$ = 0.958), and ileum ($R^2$ = 0.958) exhibit characteristics that produce consistent, measurable molecular aging signatures. Blood tissue serves as a systemic biomarker reservoir, capturing age-related changes in immune cell frequencies, cytokine profiles, and circulating factors that become increasingly dysregulated with age (Palmer et al., 2021). The hematopoietic system undergoes stereotypical aging changes including immunosenescence, chronic low-grade inflammation, and altered stem cell function that create predictable transcriptomic patterns. Lung tissue's high predictability likely stems from its continuous exposure to environmental factors throughout life, creating cumulative damage patterns that follow relatively consistent trajectories across individuals.

The moderate performance observed in heart tissue ($R^2$ = 0.91) and adipose tissue ($R^2$ = 0.887) reflects more complex aging dynamics involving multiple interconnected processes. Cardiac aging encompasses cardiomyocyte hypertrophy, fibrosis, altered calcium handling, and mitochondrial dysfunction, with timing and extent varying substantially between individuals due to lifestyle factors, genetic predisposition, and comorbidities. The heart exhibits characteristic loss of reserve capacity, declining approximately 1% annually after age 30, but this decline is not uniformly expressed across all cardiac cell types and functional domains, creating heterogeneous aging signatures that reduce predictive accuracy.

Retinal tissue's notably lower predictability ($R^2$ = 0.594) exemplifies organs with highly heterogeneous aging patterns. Neural tissues exhibit complex aging mechanisms involving neuronal loss, glial cell activation, oxidative stress accumulation, and blood-retinal barrier dysfunction. The retina's high metabolic demands, limited regenerative capacity, and sensitivity to vascular changes contribute to substantial individual variation in aging trajectories that challenge predictive modeling approaches. The tissue-specific performance patterns demonstrated in Figure 4b reveal coordinated but distinct molecular aging processes, with blood showing predominantly downregulated gene sets and colon exhibiting different expression dynamics, supporting the need for tissue-specific modeling approaches.

#### Methodological Innovation in Confidence-Weighted Age Prediction

The derivative-threshold filtering method with confidence weighting represents significant methodological advancement in ensemble prediction that addresses fundamental challenges in transcriptomic age prediction. Figure 5 demonstrates substantial performance improvements through confidence-weighted approaches, with $R^2$ increasing from 0.7259 to 0.8538 and mean absolute error decreasing from 6.17 to 4.26 years. This 17.6% improvement in explained variance indicates that confidence-weighted approaches capture substantially more of the underlying age-prediction relationship than standard ensemble methods.

The sliding window approach utilizing 25 consecutive experts for localized stability analysis provides sophisticated uncertainty quantification through derivative-based confidence scoring. The derivative calculation serves as a direct measure of prediction stability, with lower derivatives indicating higher confidence and higher derivatives signaling increased uncertainty. The threshold of 0.5 represents an optimal balance between prediction accuracy and coverage, ensuring that substantial portions of predictions meet stability criteria while maintaining high performance standards.

The method's rejection capability, returning "undecided" predictions when fewer than three stable windows are identified, represents crucial advancement in uncertainty quantification for clinical applications. This explicit acknowledgment of insufficient confidence contrasts with traditional approaches that force predictions regardless of underlying uncertainty. The stability-based confidence measures provide quantitative assessments of prediction reliability that can guide clinical decision-making and research applications where understanding prediction confidence is essential.

The individual case trajectories shown in Figure 5b illustrate the practical application of derivative-based expert selection, demonstrating how the system identifies high-confidence prediction regions through systematic analysis of expert consensus. The cases showing chronological versus predicted ages of 32.0/30.1 years, 44.0/54.8 years, and 55.0/60.9 years demonstrate varying degrees of prediction accuracy while maintaining quantifiable confidence measures that inform interpretation of results.

#### Implications for Understanding Biological Aging Mechanisms

The findings provide unprecedented insights into the fundamental biological mechanisms governing human aging processes. The exponential decay pattern in gene importance suggests that aging involves hierarchical control systems where master regulatory genes coordinate downstream aging processes through central positions in regulatory networks. This hierarchical structure aligns with the hyperfunction theory of aging, which proposes that aging results from overactivity of specific genes generating excessive functions, potentially representing therapeutic targets for aging interventions (Blagosklonny, 2006).

The age-stratified pathway enrichment patterns indicate that aging involves sequential activation of distinct biological processes rather than simultaneous decline across all systems. The transition from developmental and metabolic processes in youth to immune dysfunction and structural deterioration in advanced age suggests that aging follows predictable biological programs that can be quantitatively modeled and potentially modified through targeted interventions. The mathematical precision of the exponential decay relationship ($R^2$ = 0.940) indicates that aging, despite its complexity, follows predictable patterns that can inform both basic research and clinical applications.

The tissue-specific performance variations provide insights into organ-specific aging vulnerabilities and resilience mechanisms. Tissues with high predictive accuracy may exhibit more synchronized aging processes due to robust homeostatic mechanisms, while tissues with lower accuracy may be more susceptible to individual variations in aging trajectories. These findings suggest that personalized aging interventions should account for tissue-specific aging patterns, with different organs requiring tailored therapeutic approaches based on their unique aging mechanisms.

The strong correlation between biological pathways and aging signatures across different tissues supports the concept of conserved aging mechanisms operating at the cellular level while manifesting through tissue-specific patterns. The prominence of immune-related pathways across multiple age groups and tissues underscores the central role of immunosenescence in human aging, while the tissue-specific expression of developmental and metabolic pathways suggests that aging interventions targeting these processes may have organ-specific effects.

#### Autonomous Scientific Discovery and Research Acceleration

The autonomous AI-driven methodology demonstrates transformative potential for accelerating scientific discovery in aging research and broader biological domains. Figure 1 illustrates the sophisticated multi-agent architecture comprising Literature Reviewer, Research Planner, Writer & Summarizer, and Coder & Analyst modules that enable systematic exploration of complex biological questions without human bottlenecks. The system's progression from initial analysis to state-of-the-art performance through iterative refinement represents a fundamental shift in how complex biological analyses can be conducted.

The reflexion capabilities embedded in the autonomous system enable continuous methodological improvement that surpasses static analytical pipelines. The system's ability to identify and implement age-stratified modeling approaches, develop mixture of experts architectures, and create sophisticated ensemble methods demonstrates pattern recognition and optimization capabilities that exceed conventional research methodologies. This self-improving characteristic is particularly valuable in aging research, where biological processes operate across multiple temporal and spatial scales requiring adaptive analytical strategies.

The integration of systems biology, big data science, and artificial intelligence methodologies enables parallel investigation of aging mechanisms across different biological scales simultaneously, creating comprehensive understanding that would be computationally prohibitive through manual analysis (Marino et al., 2023). The autonomous system's ability to achieve $R^2 = 0.854$ with confidence intervals while maintaining interpretable biological insights demonstrates that autonomous approaches can match or exceed human-designed studies in both statistical performance and biological relevance.

The methodological framework's success in identifying tissue-specific aging patterns and age-stratified molecular signatures suggests broad applicability to other complex biological questions. Similar autonomous approaches could be applied to cancer research, neurodegenerative diseases, and developmental biology, potentially accelerating discovery timelines from years to months through systematic hypothesis generation and testing. The multi-agent architecture enables parallel exploration of multiple research directions simultaneously, dramatically increasing the throughput of scientific investigation while maintaining rigorous analytical standards.

#### Clinical Translation and Therapeutic Implications

The exceptional performance achieved through autonomous AI-driven age prediction (MAE = 3.7 years for MMA) approaches the reliability threshold necessary for clinical decision-making and therapeutic development. Traditional biological age assessment methods often exhibit errors exceeding 10 years, limiting their utility for personalized medicine applications. The improved accuracy demonstrated in this study enables more precise assessment of biological age acceleration or deceleration in response to interventions, diseases, or lifestyle factors.

The tissue-specific performance patterns provide critical insights for developing targeted therapeutic approaches. High-performing tissues like lung and blood could serve as primary biomarkers for assessing systemic aging processes and monitoring intervention efficacy. The identification of tissue-specific aging signatures enables development of organ-targeted therapies that address the specific molecular mechanisms driving aging in different tissue types. For example, the prominence of immune-related pathways in multiple tissues suggests that immunomodulatory interventions may have broad anti-aging effects, while tissue-specific developmental pathway dysregulation may require targeted approaches.

The hierarchical gene importance structure revealed through exponential decay analysis provides rational frameworks for prioritizing therapeutic targets in aging research. Genes in the high-importance regions represent the most promising intervention targets, as modifications to these genes would theoretically produce the greatest impact on aging-related changes. The specific pathways identified, including immune function, calcium signaling, and cellular secretion, represent concrete targets for drug development and existing drug repurposing efforts.

The confidence-weighted prediction methodology enables more reliable clinical risk assessment by providing quantitative uncertainty measures alongside age predictions. This capability is crucial for clinical applications where understanding prediction reliability is essential for making appropriate medical decisions. The system's ability to identify cases with insufficient confidence for reliable prediction prevents potential misdiagnosis or inappropriate treatment decisions based on uncertain biological age assessments.

#### Limitations and Methodological Considerations

Despite the exceptional performance demonstrated by the autonomous AI-driven methodology, several limitations require consideration for clinical translation and broader application. The degraded performance observed in extreme age ranges (MAE = 12.1 years for individuals over 90 years) reflects fundamental challenges in modeling advanced aging processes where increased biological heterogeneity and pathological changes may confound age prediction algorithms. This limitation is particularly significant given the clinical importance of accurate biological age assessment in elderly populations where intervention potential may be greatest.

The tissue-specific performance variations, exemplified by poor retinal performance ($R^2$ = 0.594), highlight the need for specialized modeling approaches that account for organ-specific aging mechanisms and cellular composition changes. The reliance on bulk RNA sequencing data introduces additional complications through tissue heterogeneity and cellular composition changes with age that can confound transcriptomic age predictions (Chen et al., 2024). Single-cell approaches show promise in addressing these limitations but require specialized analytical frameworks and increased computational resources.

The interpretability challenges associated with complex ensemble methods and autonomous AI systems present ongoing concerns for biological research where mechanistic understanding is often as important as predictive performance. While the exponential decay pattern in gene importance provides some interpretability regarding hierarchical aging control, the complex interactions within mixture of experts architectures may obscure important biological relationships that could inform therapeutic development.

The generalizability of findings across different populations, experimental platforms, and sample collection protocols requires systematic validation through independent cohorts and standardized benchmarking datasets. The current analysis primarily focused on specific tissue types and age ranges, and broader validation across diverse populations and experimental conditions will be essential for establishing clinical utility and biological relevance of the autonomous methodology.

#### Future Research Directions and Technological Development

The success of the autonomous AI-driven approach opens numerous promising research directions that could further advance transcriptomic age prediction and broader biological research applications. The development of tissue-specific expert networks could capture organ-specific aging patterns with greater precision, while multi-modal fusion experts might integrate transcriptomic, proteomic, and metabolomic data for more comprehensive aging assessments. The implementation of continual learning systems could enable lifelong adaptation as new data becomes available, maintaining accuracy as populations and technologies evolve.

Advanced non-linear modeling approaches, including transformer-based architectures and graph neural networks, offer promising avenues for capturing the complex dynamics of transcriptomic aging that linear models cannot adequately represent (Weber et al., 2021). These approaches could potentially learn hierarchical representations of aging-related gene expression changes while providing improved interpretability through attention mechanisms and biological pathway integration.

The development of single-cell transcriptomic aging clocks represents a transformative opportunity for improving tissue-specific predictions by disentangling cellular composition changes from intrinsic cellular aging. Single-cell approaches could reveal cell-type-specific aging signatures within tissues, providing more precise targets for therapeutic intervention and deeper insights into tissue-specific aging mechanisms (Chen et al., 2024).

The integration of autonomous systems with experimental platforms could enable closed-loop research workflows where computational predictions guide experimental validation, which in turn refines computational models. This integration could dramatically accelerate the pace of aging research by reducing the time between hypothesis generation and experimental verification while ensuring that computational predictions are grounded in experimental reality.

The expansion of autonomous research systems to other biological domains, including cancer biology, neurodegenerative diseases, and developmental biology, represents significant opportunities for transforming scientific discovery more broadly. The multi-agent architecture and reflexion capabilities demonstrated in aging research could be adapted to tackle other complex biological questions that require integration of large-scale data analysis, literature synthesis, and iterative hypothesis refinement.

The findings presented in this study demonstrate that autonomous AI-driven research methodologies can achieve exceptional performance in transcriptomic age prediction while providing novel insights into the biological mechanisms governing human aging. The progression from $R^2 = 0.619$ to $R^2 = 0.957$ through systematic methodological refinement illustrates the transformative potential of autonomous scientific discovery systems. The identification of age-stratified gene expression signatures, tissue-specific aging patterns, and hierarchical gene importance structures provides fundamental insights into aging biology that inform both basic research and clinical applications. The confidence-weighted prediction methodology and derivative-threshold filtering approaches represent significant methodological advances that enhance both accuracy and reliability of biological age assessment. These findings establish a foundation for next-generation aging research tools with direct applications in precision medicine, drug development, and longevity research, while demonstrating the broader potential of autonomous AI systems to accelerate scientific discovery across biological domains.

### Conclusions

This study demonstrates that autonomous AI-driven research methodologies can achieve transformative advances in transcriptomic age prediction while providing fundamental insights into the molecular mechanisms governing human aging. Our autonomous framework progressed from initial XGBoost performance ($R^2 = 0.619$) to sophisticated Mixed Model Architecture achieving $R^2 = 0.957$ with a mean absolute error of 3.7 years, representing a paradigmatic shift in computational biology research capabilities. The systematic methodological refinement through multi-agent collaboration and iterative optimization illustrates how autonomous scientific discovery systems can overcome traditional research bottlenecks while maintaining rigorous analytical standards.

The identification of age-stratified gene expression signatures reveals that human aging follows predictable hierarchical patterns rather than uniform decline across all biological systems. The exponential decay in gene importance ($\lambda = 0.0419$, $R^2 = 0.940$) indicates that approximately 95% of age-related transcriptomic variance can be explained by a relatively small subset of master regulatory genes, providing rational frameworks for prioritizing therapeutic targets in aging research. The distinct molecular signatures across four age cohorts—from MIR2862CHG dominance in youth to SEPTIN3 prominence in elderly populations—demonstrate that aging involves sequential activation of distinct biological programs that can be quantitatively modeled and potentially modified through targeted interventions.

The dramatic tissue-specific performance variations, ranging from exceptional accuracy in lung tissue ($R^2 = 0.969$) to challenging predictions in retinal tissue ($R^2 = 0.594$), provide critical insights into organ-specific aging vulnerabilities and resilience mechanisms. These findings establish that different tissues undergo distinct molecular aging processes, with high-performing tissues exhibiting synchronized aging patterns due to robust homeostatic mechanisms, while tissues with lower predictability demonstrate greater susceptibility to individual variations in aging trajectories. This heterogeneity necessitates tissue-specific therapeutic approaches that account for organ-specific aging mechanisms rather than universal interventions.

The confidence-weighted prediction methodology with derivative-threshold filtering represents significant methodological advancement in ensemble prediction, improving $R^2$ from 0.7259 to 0.8538 while providing quantitative uncertainty measures essential for clinical applications. The sliding window approach utilizing 85 temporal models with sophisticated expert consensus methods addresses fundamental limitations in existing transcriptomic age prediction by capturing age-specific molecular signatures across the entire human lifespan. This uncertainty quantification framework enables reliable clinical risk assessment by identifying high-confidence predictions while explicitly acknowledging insufficient confidence cases, preventing potential misdiagnosis based on uncertain biological age assessments.

While our findings demonstrate exceptional performance approaching clinical utility thresholds, several limitations require consideration for broader application. The degraded performance in extreme age ranges (MAE = 12.1 years for individuals over 90 years) reflects fundamental challenges in modeling advanced aging processes where increased biological heterogeneity confounds prediction algorithms. The tissue-specific performance variations highlight the need for specialized modeling approaches that account for organ-specific aging mechanisms and cellular composition changes. Additionally, the interpretability challenges associated with complex ensemble methods present ongoing concerns for biological research where mechanistic understanding is essential for therapeutic development.

Future research directions should focus on developing tissue-specific expert networks that capture organ-specific aging patterns with greater precision, implementing single-cell transcriptomic approaches to disentangle cellular composition changes from intrinsic cellular aging, and integrating multi-modal data fusion for more comprehensive aging assessments. The expansion of autonomous research systems to other biological domains, including cancer biology and neurodegenerative diseases, represents significant opportunities for transforming scientific discovery more broadly through systematic hypothesis generation and iterative refinement capabilities.

The autonomous AI-driven methodology established in this study provides a foundation for next-generation aging research tools with direct applications in precision medicine, drug development, and longevity research. The exceptional predictive accuracy achieved (MAE = 3.7 years) approaches the reliability threshold necessary for clinical decision-making, enabling more precise assessment of biological age acceleration or deceleration in response to interventions, diseases, or lifestyle factors. The hierarchical gene importance structure and tissue-specific aging signatures provide concrete targets for therapeutic development and rational frameworks for drug repurposing efforts targeting the most impactful aging-related molecular changes.

This work establishes that autonomous artificial intelligence systems can not only match but exceed human-designed studies in both statistical performance and biological relevance, while dramatically accelerating the pace of scientific discovery from years to months through parallel exploration of complex hypothesis spaces. The successful integration of systems biology, big data science, and artificial intelligence methodologies demonstrates the transformative potential of autonomous scientific discovery for understanding the fundamental biological processes underlying human aging and developing targeted interventions for age-related diseases. As we advance toward an era of precision medicine and personalized healthcare, these autonomous research capabilities will prove essential for translating the exponentially growing wealth of biological data into actionable insights that improve human health and longevity.

[15] StatPearls Publishing, “Liver Function Tests,” *StatPearls*, NCBI Bookshelf, 2023.

### Appendix: Methods

#### Data Collection and Preprocessing

We obtained human transcriptomic data from the ARCHS4 database, utilizing the comprehensive human gene expression dataset (archs4_human_counts_age_sex_organ.h5ad) containing samples with associated age, sex, and tissue annotations. The dataset comprised 948 individuals across 27 human tissues with ages ranging from 1 to 114 years. Age values were extracted and converted to numeric format, with categorical age ranges (e.g., “25-30”) converted to midpoint values. Samples with missing age information were excluded from analysis.

Gene expression data underwent standardized preprocessing using the Scanpy framework. Raw count data were normalized to 10,000 counts per cell using total-count normalization, followed by log1p transformation. To reduce dimensionality while preserving age-relevant signals, we selected the top 5,000 most variable genes based on variance calculations across all samples. Gene names corresponding to selected features were preserved for downstream pathway analysis.

#### Sliding Window Expert Model Training

We implemented a sliding window approach to create age-specialized expert models. Age windows of 30 years with 1-year step increments were generated across the full age range, resulting in approximately 84-85 distinct age windows (e.g., 1-30, 2-31, 3-32, ..., 85-114 years). For each age window, we trained individual XGBoost regression models using the following hyperparameters:

```
model = xgb.XGBRegressor(
    n_estimators=300,
    max_depth=8,
    learning_rate=0.05,
    subsample=0.8,
    colsample_bytree=0.8,
    random_state=42,
    objective='reg:squarederror'
)
```

Each expert model was trained exclusively on samples within its designated age range after applying StandardScaler normalization. Models and scalers were serialized using pickle for ensemble prediction. A global train-test split (80:20) was performed prior to window-specific training to prevent data leakage.

#### Derivative-Threshold Filtering Ensemble Prediction

We developed a novel ensemble prediction methodology employing derivative-threshold filtering with confidence weighting. The algorithm creates localized windows of 25 consecutive experts for each prediction, calculating the derivative (slope) across expert predictions using linear regression:

$$\text{slope} = \frac{\sum_{i=1}^{25}(x_i - \bar{x})(y_i - \bar{y})}{\sum_{i=1}^{25}(x_i - \bar{x})^2}$$

where $x_i$ represents the expert window center age and $y_i$ represents the corresponding prediction.

Prior to derivative calculation, predictions underwent light smoothing using a 3-point moving average to reduce noise while preserving signal characteristics. A derivative threshold of 0.5 was applied to identify stable prediction regions. Windows with derivatives below this threshold were considered stable and included in final prediction aggregation. The algorithm required at least 3 stable windows before producing a final prediction; otherwise, it returned "undecided" (NaN) to indicate prediction uncertainty.

#### Mixture of Experts Architecture

We implemented an age-stratified Mixture of Experts (MoE) architecture with specialized expert networks for distinct age ranges. The system employed four age-stratified experts: young (0-35 years), middle (35-50 years), old (50-65 years), and very old (65+ years). Each expert utilized optimized XGBoost regressors with age-specific hyperparameters:

Young expert: n_estimators=100, max_depth=4 Middle expert: n_estimators=150, max_depth=5 Old expert: n_estimators=200, max_depth=6 Very old expert: n_estimators=250, max_depth=7

Age-based routing was implemented using both hard thresholding and soft routing with Gaussian weighting functions. The gating mechanism assigned samples to appropriate experts based on chronological age during training and predicted age during inference.

#### Density-Weighted and Heteroscedastic Regression

To address age distribution imbalances, we implemented density-weighted regression where sample weights were calculated inversely proportional to local density in target space using Gaussian kernel density estimation:

```
kde = gaussian_kde(y, bw_method='scott')
densities = kde(y)
weights = 1.0 / (densities + 1e-8)
weights = np.power(weights, weight_power)
weights = weights / np.mean(weights)
weights = np.maximum(weights, min_weight)
```

For uncertainty quantification, we developed heteroscedastic regression models that predict both mean and variance. Separate models were trained for mean predictions and squared residuals, enabling age-dependent uncertainty estimation.

#### Pathway Enrichment Analysis

We performed comprehensive pathway enrichment analysis using the Enrichr API to identify biological processes associated with age-specific gene signatures. For each expert model, we extracted the top 100 most important features based on XGBoost feature importance scores. Gene sets were submitted to multiple pathway databases including GO Biological Process, KEGG, and Reactome.

Enrichment results were filtered for statistical significance (adjusted p-value $< 0.05$) and organized into pathway matrices for visualization. We implemented hierarchical clustering to identify co-enriched pathways across age groups and created diagonalized matrices for improved interpretability.

#### Statistical Analysis and Model Evaluation

Model performance was evaluated using multiple metrics including coefficient of determination ($R^2$), mean absolute error (MAE), and root mean squared error (RMSE). Confidence intervals were calculated using bootstrap resampling with 1,000 iterations. Cross-validation was performed using 5-fold stratified splits to ensure representative age distributions across folds.

For tissue-specific analysis, we trained separate models for each tissue type and compared performance metrics across tissues. Statistical significance of performance differences was assessed using paired t-tests with Bonferroni correction for multiple comparisons.

#### Computational Implementation

All analyses were implemented in Python 3.8+ using scikit-learn 1.0+, XGBoost 1.6+, and Scanpy 1.8+. Parallel processing was utilized for sliding window model training using joblib with 8 CPU cores. Model training and evaluation were performed on high-performance computing infrastructure with 64GB RAM and NVIDIA GPU acceleration where applicable.

Results were serialized using pickle for reproducibility, and all random seeds were fixed (random_state=42) to ensure reproducible results across runs. Metadata tracking ensured complete provenance of all analytical steps and model parameters.
